# The human GID complex engages two independent modules for substrate recruitment

**DOI:** 10.1101/2021.04.07.438752

**Authors:** Weaam I. Mohamed, Sophia L. Park, Julius Rabl, Alexander Leitner, Daniel Boehringer, Matthias Peter

**Author notes:** equal contribution (Sophia L. Park, Julius Rabl).

## Abstract

The human GID (hGID) complex is an evolutionary conserved E3 ubiquitin ligase regulating diverse biological processes including glucose metabolism and cell cycle progression. However, the biochemical function and substrate recognition of the multi-subunit complex remains poorly understood. While the yeast GID complex recognizes Pro/N-end rule substrates via yeast Gid4, the human GID complex requires a WDR26/Gid7-dependent module to trigger proteasomal degradation of mammalian HBP1. Here, using biochemical assays, crosslinking-mass spectrometry and cryo-electron microscopy, we show that hGID unexpectedly engages two distinct modules for substrate recruitment, dependent on either WDR26 or GID4. WDR26 together with RanBP9 cooperate to ubiquitinate HBP1 *in vitro*, while GID4 is dispensable for this reaction. In contrast, GID4 functions as an adaptor for the substrate ZMYND19, which surprisingly lacks a Pro/N-end rule degron. GID4 substrate binding and ligase activity is regulated by ARMC8α, while the shorter ARMC8β isoform assembles into a stable hGID complex that is unable to recruit GID4. Cryo-EM reconstructions of these hGID complexes reveal the localization of WDR26 within a ring-like, tetrameric architecture and suggest that GID4 and WDR26/Gid7 utilize different, non-overlapping binding sites. Together, these data advance our mechanistic understanding of how the hGID complex recruits cognate substrates and provide insights into the regulation of its ligase activity.

## Introduction

The ubiquitin-proteasome system (UPS) is required for cells to adjust to different nutrient conditions such as limiting carbon sources. Changing metabolic flux is often controlled by regulating the relative abundance of rate-limiting enzymes that function in distinct exergonic pathways (Nakatsukasa *et al*, 2015). In yeast, gluconeogenesis and glycolysis are intermittently coordinated to prevent simultaneous glucose production and break-down. This is achieved in part by the <u>G</u>lucose <u>i</u>nduced <u>d</u>eficient degradation (GID) complex (Santt *et al*, 2008), a multi-subunit E3 ligase that specifically targets the surplus of gluconeogenic enzymes for proteasomal degradation, including the conserved Fructose-1,6-bisphosphatase 1 (Fbp1). Adequate glucose levels induce expression of its critical subunit Gid4 (Santt *et al*, 2008), which is otherwise degraded by autoubiquitination. Interestingly, Gid4 functions as a substrate receptor recognizing a Pro/N-end degron motif (Chen, 2017; Dong *et al*, 2018; Qiao *et al*, 2019). Gid4 is partially redundant with Gid10, which is upregulated by heat and osmotic stress conditions (Qiao *et al*, 2019; Melnykov *et al*, 2019). Moreover, Gid11/Ylr149c was recently identified as a GID substrate receptor recognizing proteins with N-terminal threonine residues (Edwin Kong *et al*, 2021), thus expanding the specificity of the GID complex. Interestingly, these substrate receptors are recruited to the GID-complex by binding to Gid5, which, in turn, interacts with the catalytic core composed of Gid8 and the RING-containing subunits Gid2 and Gid9. Structural analysis of the monomeric GID-complex also identified an essential role of Gid1, which interacts with Gid8 and Gid5. In contrast to these subunits, Gid7 is not required to degrade gluconeogenic enzymes (Menssen *et al*, 2018). Indeed, Gid7 does not stably incorporate into the yeast GID complex (Qiao *et al*, 2019), and the role of Gid7 in yeast thus remains unclear.

Interestingly, the GID E3 ligase complex is highly conserved, and all seven yeast GID subunits have homologous counterparts in humans. RanBP9 (Gid1), RMND5a (Gid2), ARMC88 (Gid5), TWA1 (Gid8) and MAEA (Gid9) are ubiquitously expressed and assemble into a high-molecular weight complex localizing to the nucleus and cytoplasm (Kobayashi *et al*, 2007). The human GID complex (hGID) is also referred to as C*-*terminal to LisH (CTLH) complex after a sequence motif shared between five subunits (Kobayashi *et al*, 2007). Like in yeast, the two RING-domain containing subunits RMND5a and MAEA1 linked by TWA1 form the catalytic core of the E3 ligase (Lampert *et al*, 2018). Besides this catalytic trimer, the hGID complex assembles with other subunits such as WDR26 (Gid7), RanBP9/RanBP10 (Gid1), MKLN1, GID4, ARMC8 and YPEL5 (Kobayashi *et al*, 2007; Lampert *et al*, 2018). WDR26 contains a WD40-domain, which typically folds into a characteristic beta-propeller and frequently exits in substrate receptors of the Cullin 4 RING E3 ubiquitin ligase family (CRL4) (Higa *et al*, 2006). RanBP9 and RanBP10 contain a SPRY beta-domain, which is commonly present in TRIM RING E3 ligases (DCruz *et al*, 2013), and ARMC8 contains armadillo-like domains, which also serve as platform for various protein-protein interactions (Huber *et al*, 1997). Interestingly, mammalian cells express two ARMC8 isoforms, ARMC8α and ARMC8β, resulting from alternative splicing of the same gene (Kobayashi *et al*, 2007; Maitland *et al*, 2019; Tomaru *et al*, 2010). Both ARMC8α and ARMC8β incorporate into the hGID complex (Kobayashi *et al*, 2007; Maitland *et al*, 2019), but the structural and functional differences between the two remain poorly explored. Therefore, although the different subunits are evolutionary conserved and the catalytic core of hGID resembles the yeast complex, further work is required to understand the assembly and structural organization of this intricate E3 ligase in mammalian cells.

The biological functions of the mammalian GID E3 ligase are only beginning to emerge and to date there is no evidence that links hGID ligase function to glucose metabolism. Although the binding pocket in human GID4 is conserved, endogenous substrates governed by the Pro/N-end degron motif have not been identified. Despite this, the GID complex has been linked to cell proliferation in human cells, at least in part by targeting the transcriptional repressor HMG box protein 1 (HBP1) for proteasomal degradation (Lampert *et al*, 2018). HBP1 inhibits cell cycle progression by regulating the retinoblastoma tumor suppressor (Rb), and also regulates the expression of genes involved in differentiation and apoptosis. Interestingly, this role of the hGID complex in regulating cell cycle progression and HBP1 stabilization requires not only the catalytic core subunits, but also WDR26/Gid7.

Consistent with this role in cell proliferation, numerous studies have reported significantly increased expression of multiple GID subunits across a variety of human tumor cells and tissues (Both *et al*, 2016; Jiang *et al*, 2016; 2015b; 2015a; Liang *et al*, 2016; Zhao *et al*, 2016; Zhou *et al*, 2016). Most notably, elevated WDR26 protein levels correlate with poor disease prognosis in many cancers, where available large cancer datasets highlighted gene amplification of WDR26 with a remarkable prevalence of up to 55% in breast, ovarian, and prostate cancers (Cerami *et al*, 2012; Gao *et al*, 2013). Additionally, ARMC8α, but not ARMC8β, was found to promote cell proliferation and invasion of non-small cell lung cancer cells (Xie *et al*, 2014). ARMC8α was also shown to bind and target α-catenin for proteasomal degradation, and may interact with <u>H</u>epatocyte growth factor-<u>R</u>egulated tyrosine kinase <u>S</u>ubstrate (HRS). However, little is known about the ARMC8β subunit and its role in the function and regulation of the hGID E3 ligase complex.

Several subunits of the hGID complex, namely RanBP9, RanBP10, WDR26 and MKLN1, have been linked to neurodegeneration and amyloid β (Aβ) pathologies (Her *et al*, 2017; Woo *et al*, 2015), intellectual disability (Skraban *et al*, 2017), and early onset bipolar diseases and schizophrenia (Nassan *et al*, 2017; BAE *et al*, 2015). Moreover, suppression of RMND5a in *Xenopus laevis* leads to malformations in the fore and midbrain (Pfirrmann *et al*, 2015), suggesting that the GID complex may regulate brain development and neuronal functions. RanBP9 is ubiquitously expressed and the majority of knock-out mice die immediately after birth (Puverel *et al*, 2011). The few survivors are significantly smaller in size and cannot undergo spermatogenesis or oogenesis, suggesting that the GID complex may function in growth control and meiosis.

Despite the multitude of evidence supporting a role of the hGID complex in multiple biological processes, few critical substrates have been identified that can explain the underlying phenotypes. Moreover, it remains unclear whether these diverse cellular functions of the complex require its E3 ligase activity, and whether they involve all or just a subset of the known hGID subunits. Therefore, it is crucial to better understand the function and regulation of the different hGID subunits and, in particular, elucidate the mechanism of substrate recruitment.

Previous AP-MS studies not only identified novel hGID subunits, but also sub-stoichiometrically associated proteins such as HBP1, ZMYND19 and HTRA2 (Boldt *et al*, 2016; Lampert *et al*, 2018). HBP1 binds the hGID complex preferentially in proteasome-inhibited cells, consistent with being a physiological substrate (Lampert *et al*, 2018). HTRA2 encodes a mitochondrial serine protease that induces cell death by regulating cytosolic inhibitors of apoptosis (IAPs), leading to increased caspase activity. Zinc finger MYND domain-containing protein 19 (ZMYND19) interacts with multiple hGID subunits, including TWA1, ARMC8 and RMND5a (Boldt *et al*, 2016). Although ZMYND19 protein levels are upregulated in hepatocellular carcinoma (Zhu *et al*, 2018), its biological functions remain unclear.

In this study, we combined cell biology, biochemistry and cryo-electron microscopy to elucidate the assembly and molecular mechanisms of the hGID E3 ligase, with a particular emphasis on subunits involved in substrate recruitment. Interestingly, we found that the hGID E3 ligase engages two independent modules for substrate recruitment, comprised of either WDR26/RanBP9 or GID4/ARMC8. We identified and characterized the minimal hGID complex required for HBP1 degradation *in vitro*, composed of WDR26 together with the catalytic core subunits MAEA, RMDN5a and TWA1. We further show that ZMYND19 is targeted for degradation by hGID in a GID4-dependent manner, although it lacks a Pro/N-end rule degron motif. Finally, we propose distinct roles for the ARMC8 isoforms; while both ARMC8α and ARMC8β assemble stable hGID complexes, only ARMC8α is able to recruit GID4.

## Results

### The hGID complex uses distinct substrate modules to target different substrates

In order to identify subunits within the hGID complex that are involved in substrate recruitment, we generated siRNA against ARMC8, GID4, RanBP9 and WDR26. While siRNA-depletion of ARMC8 and GID4 expression did not affect endogenous protein levels of HBP1, reduction of RanBP9 and WDR26 lead to an accumulation of HBP1 in HeLa Kyoto cells (**Fig. 1A)**. Consistently, ectopic co-expression of WDR26 and HBP1 prominently decreased HBP1 levels in a MG132-dependent manner, which was not the case when HBP1 was co-expressed with GID4 (**Fig. 1B**). Conversely, overexpression of GID4, but not WDR26, substantially decreased ZMYND19 levels (**Fig. 1C**). Taken together, these data suggest that HBP1 is targeted for proteasomal degradation in a WDR26/RanBP9-dependent manner, while ZMYND19 is a GID4/ARMC8-dependent substrate of the hGID complex (**Fig. 1D**).

**Figure 1:**
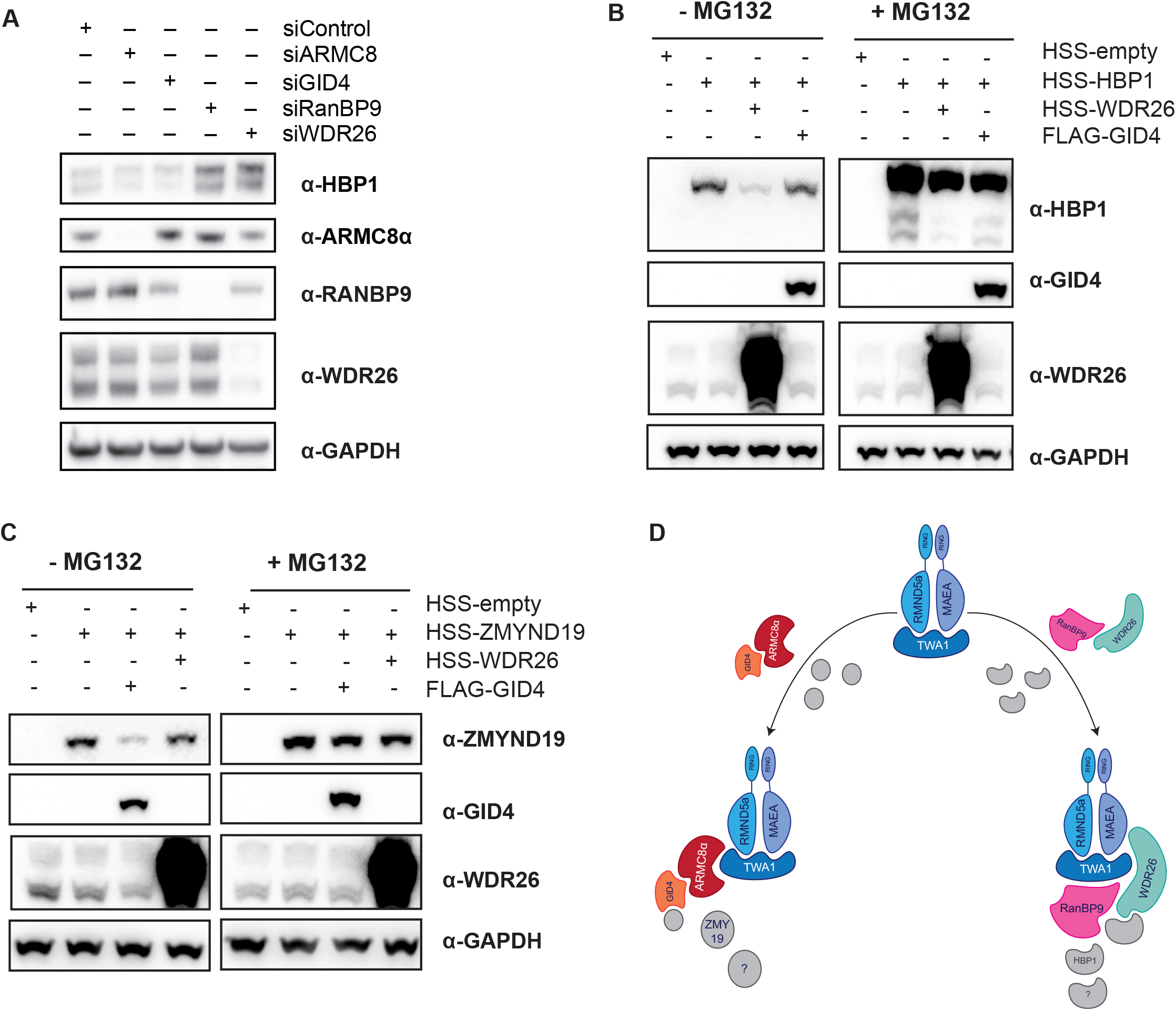
The hGID complex uses distinct substrate modules to target different substrates. **A.** Immunoblot of samples following depletion of WDR26, RanBP9, ARMC8, and GID4 using pools of siRNAs for 72 hr in HeLa Kyoto cells, and monitoring the endogenous protein levels of HBP1. **B.** Western blotting of samples after ectopic overexpression in HEK-293T of HBP1 either alone, or with WDR26 or GID4. The stability of HBP1 was monitored after treatment of MG132 or DMSO for 10 hr. **C.** Immunoblot of samples following ectopic overexpression in HEK-293T of ZMYND19 either alone, or with WDR26 or GID4. The stability of ZMYND19 was monitored after treatment of MG132 or DMSO for 10 hr. **D.** A schematic representation visualizing the hGID E3 ligase complex recruiting two distinct modules for substrate recruitment.

To biochemically test this hypothesis, we conducted *in vitro* ubiquitination assays for HBP1 and ZYMND19 in the presence of hGID complexes with defined subunit composition. Different hGID sub-complexes and full-length GID4 were purified from *Sf9* insect cells using a multi-step column purification (**Fig. 2A**, **Suppl. Fig. 1A-D**), and likewise, the substrates HBP1 and ZMYND19 were expressed and purified to homogeneity (**Fig. 2B**, **Suppl. Fig. 1E** **and** **F**). Interestingly, the minimal hGID complex required to achieve efficient HBP1 ubiquitination was composed of the catalytic core (MAEA, RMND5a and TWA1) together with WDR26 (**Fig. 2C**). Although RanBP9 forms a stable complex with WDR26 (**Suppl. Fig. 1G**), addition of RanBP9 only slightly enhanced HBP1 ubiquitination (**Fig. 2C**). hGID complexes lacking both WDR26 and RanBP9, but containing ARMC8, were unable to ubiquitinate HBP1 (**Fig. 2C**), Likewise, hGID complexes composed of the core subunits MAEA, RMND5a and TWA1, together with ARMC8 and GID4 only poorly ubiquitinated HBP1 *in vitro* (**Fig. 2D**). Addition of GID4 and/or ARMC8 to complexes containing WDR26/RanBP9 had no effect (**Fig. 2D**). Thus, we conclude that WDR26/RanBP9, but not the GID4/ARMC8 module, promotes the E3 ligase activity of the hGID complex towards HBP1.

**Figure 2:**
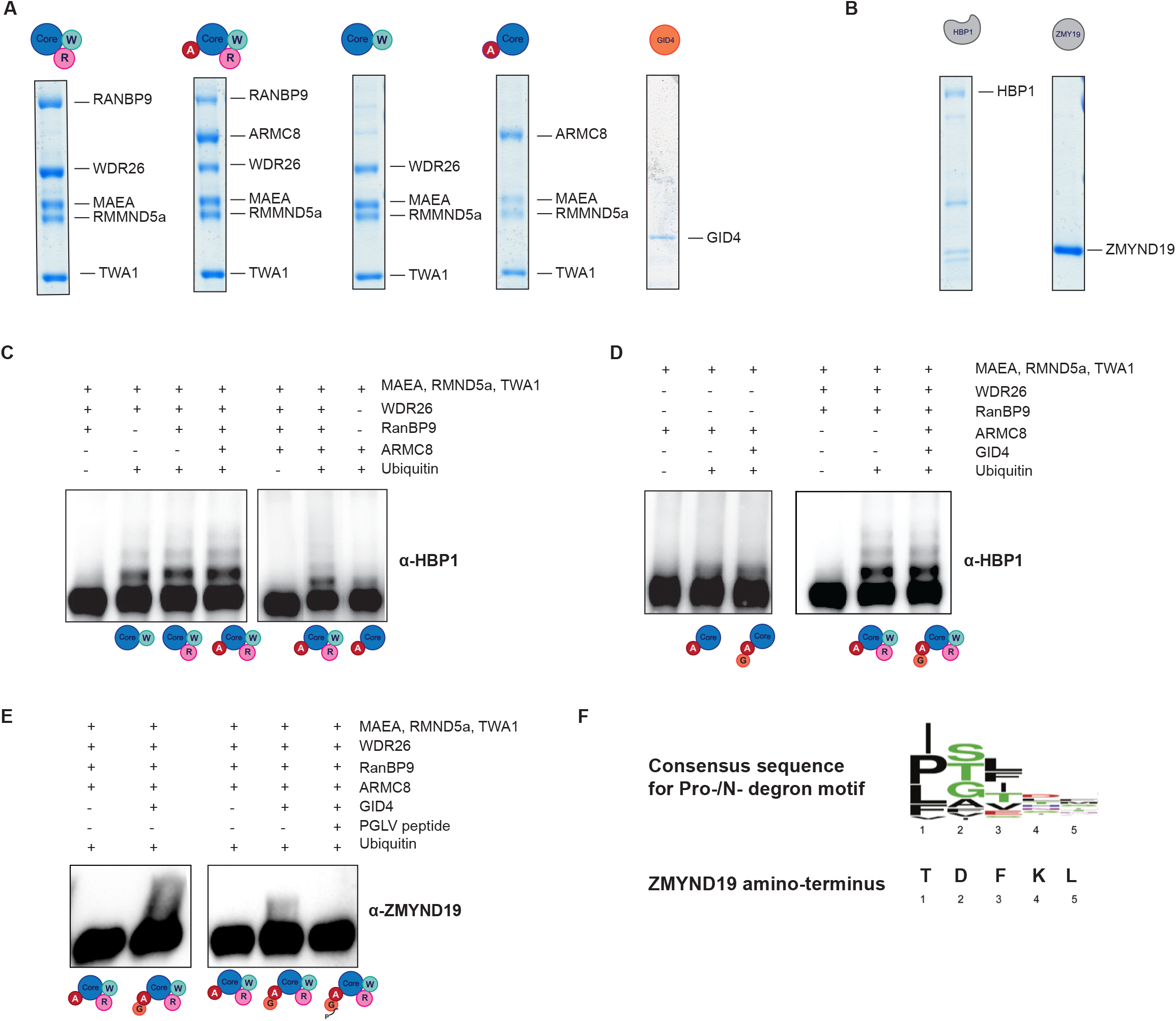
Distinct substrate recruitment modules are required to ubiquitinate HBP1 and ZMYND19 *in vitro*. **A.** Coomassie-stained SDS-PAGE showing purified hGID sub-complexes used for *in vitro* ubiquitination assays. The catalytic core composed of MAEA, RMND5a, and TWA1 is colored in blue, WDR26 in dark cyan, RanBP9 in light magenta, ARMC8 in dark red and GID4 in orange. **B.** Coomassie-stained SDS-PAGE showing the purified hGID substrates, HBP1 and ZMYND19. **C.** and **D.** Western blot analysis of the *in vitro* ubiquitinated HBP1, which was performed by mixing purified HBP1 with ubiquitin E1, UBCH5a, and ubiquitin in the presence of the indicated hGID sub-complexes. **E.** Immunoblots of *in vitro* ubiquitination of ZMYND19, which was performed by mixing purified ZMYND19 with ubiquitin E1, UBE2H, ubiquitin and the 6-subunit hGID complex (ARCM8, RanBP9, WDR26, MAEA, RMND5a, and TWA1) in the presence or absence of GID4 and a 10-fold excess of the PGLV GID4-specific peptide. **F.** Comparison of the N-terminal sequences of the first five amino acids of the Pro-/N-degron consensus motif (Dong et. al. 2020) and human ZMYND19 (Q96E35).

Conversely, ZMYND19 ubiquitination *in vitro* was dependent on the GID4 subunit, as a hGID complex containing the core subunits along with WDR26, RanBP9 and ARMC8 was not capable of ubiquitinating ZMYND19 (**Fig. 2E**). This ubiquitination was substantially inhibited in the presence of a 10-fold excess of a GID4-specific peptide (Dong *et al*, 2018), consistent with a role of GID4-mediated targeting of ZMYND19. Surprisingly, ZMYND19 does not contain a Pro/N-end rule-degron (**Fig. 2F**), implying that the GID4 binding pocket may also recognize substrates via internal degron motifs.

### WDR26/RanBP9-containing hGID complexes assemble ring-shaped tetramers

Size exclusion purification of the HBP1-targeting hGID complex (MAEA, RMND5a, TWA1, WDR26 and RanBP9) by Superose 6 column showed one predominant peak with an elution profile much larger than the expected monomeric size of 260 kDa. Consistently, oligomerization of hGID was confirmed by SEC-MALS analysis, where the five-subunit hGID complex eluted in a broad peak largely at 1.1 MDa, indicative of a tetrameric assembly (**Fig. 3A**). In contrast, hGID complexes lacking RanBP9 revealed two peaks with identical subunit composition (**Suppl. Fig. 1C** **and** **D**), suggesting an equilibrium between two oligomeric states. Oligomerization of the hGID complex also occurs *in vivo*, as shown by co-immunoprecipitation of differentially-tagged subunits (Kobayashi *et al*, 2007). Moreover, MAEA, RMND5a, TWA1, WDR26 and RanBP9 are found in the same peak fraction with a proposed molecular weight of more than 1.6 MDa in the SECexplorer web platform (**Suppl. Fig. 1H**) (Heusel *et al*, 2019).

**Figure 3:**
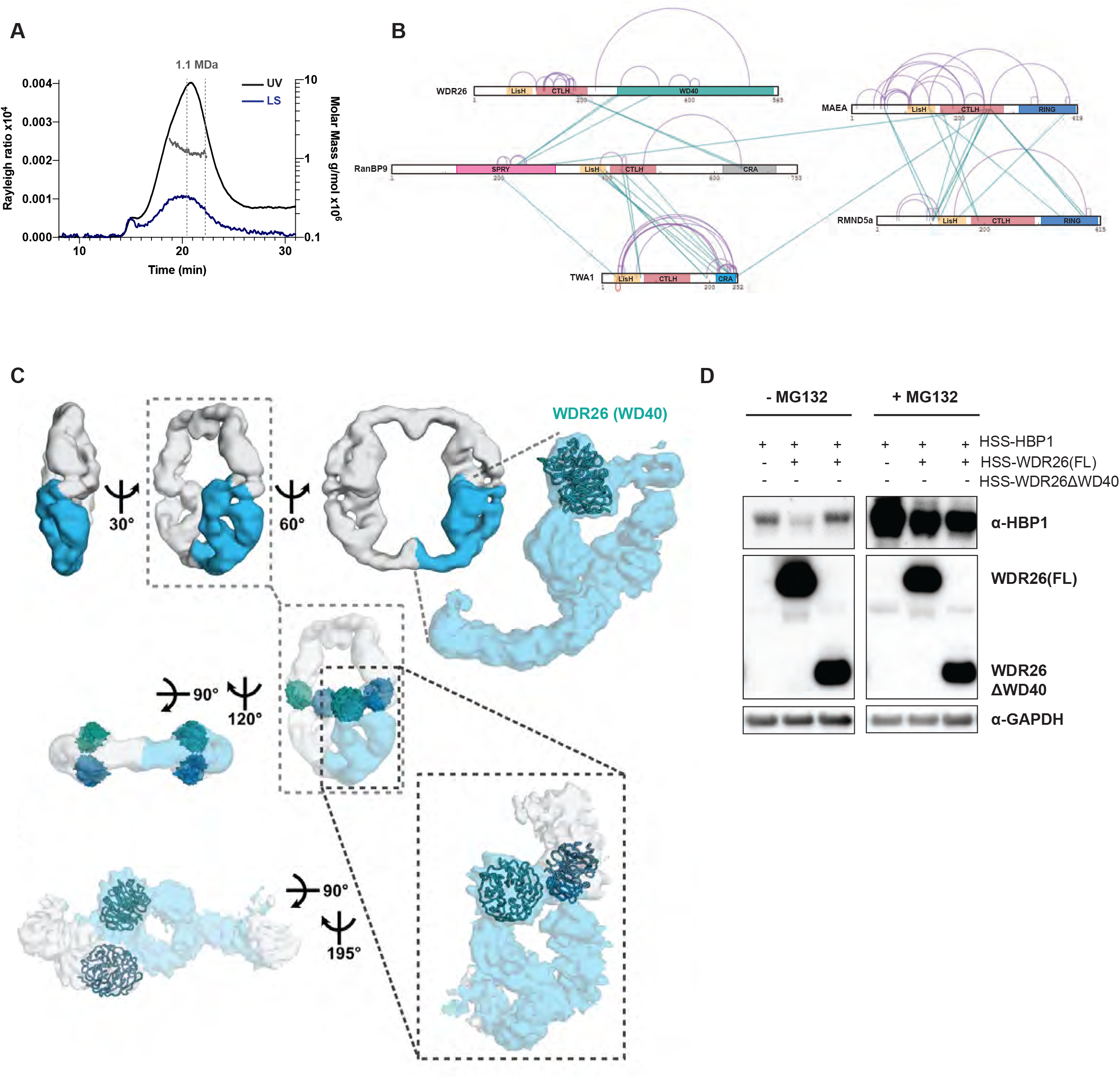
WDR26/RanBP9-containing hGID complexes assemble ring-shaped tetramers. **A.** Chromatogram of the SEC-MALS analysis at a flow rate of 0.5 ml/min, showing the UV curve and the Rayleigh ratio (1/cm) at a scattering angle of 90 degrees (left y-axis), together with the molar mass (MDa) of the peaks determined by MALS (right y-axis). The peak fraction showing a homogenous size distribution at around 1.1 MDa is labeled with gray dotted lines. **B.** XL-MS analysis of the 5-subunit hGID complex (RanBP9, WDR26, RMND5a, MAEA, and TWA1). Cross-links within different complex subunits are indicated by green lines, and cross-links within the same subunit are indicated with purple lines. The predicted domain boundaries of the different subunits are colored as follows: LisH domain in light orange, CTLH domain in dark orange, RING domains in blue, TWA1’s CRA domain in light blue, WD40 in dark cyan, and SPRY in light magenta. **C.** Rotational views of the cryo-EM map of the 5-subunit hGID complex (RanBP9, WDR26, RMND5a, MAEA, and TWA1) at 12 Å resolution. The higher resolution cryo-EM map at 9 Å produced by particle symmetry expansion is shown in blue. The dotted rectangle highlights the positions of the fitted WD40 domains from two asymmetric units. **D.** Western blotting of samples following ectopic overexpression of HBP1 either alone, or with full-length (FL) or WD40-truncated WDR26 (ΔWD40) in HEK-293T. HBP1 levels were monitored in cells treated with MG132 or DMSO for 12 hr.

To gain better molecular insight into the assembly and oligomerization of the hGID-RanBP9/WDR26 complex, we performed cross-linking mass spectrometry analysis (XL-MS) (**Fig. 3B**). As expected, extended interactions were detected between the two RING-domain containing subunits (MAEA and RMND5a) via their LisH and CTLH domains. LisH and CTLH domains form thermodynamically stable dimers (Gerlitz *et al*, 2005), and are thus expected to be involved in assembly. A dense cross-linking pattern was also detected between RanBP9’s LisH and CTLH domains and the CRA domain of TWA1. RanBP9’s SPRY domain also interacts with the WD40 domain of WDR26, while no cross-links could be observed between WDR26 and the other subunits (**Fig. 3B**). Based on these data, we speculate that RanBP9 adopts an elongated structure characteristic of a scaffolding function.

To corroborate these interactions, we pursued single particle cryo-electron microscopy (cryo-EM) analysis of the stable hGID complex composed of its catalytic core (MAEA, RMND5a, TWA1) bound to the WDR26/RanBP9 substrate module (**Suppl. Fig. 2A)**. Single particle analysis of the five subunit GID complex (**Suppl. Table 3**) revealed that hGID assembles into a ring-shaped complex with a diameter of ~270 Å. 2D classification of the particles showed circular class averages with twofold symmetry (Scheres, 2016) **(Suppl. Fig. 2B**). The circular scaffold is approximately 25 Å wide, and is decorated with inward facing protrusions. Comparing the 2D classes revealed that the ring diameter varies slightly, which indicates flexibility of the scaffold ring. Initial model generation with CryoSPARC (Punjani *et al*, 2017) uncovered a D2 symmetric arrangement, consistent with a tetramer of five-subunit GID complexes. To address the conformational flexibility of the scaffold ring, we employed 3D classification after symmetry expansion to refine a cryo-EM map of the asymmetric unit to subnanometer resolution (**Fig. 3C**). In the cryo-EM map of the asymmetric unit, we could locate the WD40-propeller of WDR26, which represents the largest protein fold present in the GID subunits. The resolution did not allow an unambiguous assignment of the alpha-helical modules or other domains. The WD40-propeller of WDR26 protrudes from an elongated scaffold-like density. In the tetramer, two WDR26 subunits contact each other via their WD40 propellers, suggesting a possible role in oligomerization. Moreover, the WD40 propeller seems to be in a conformation primed for substrate recruitment (**Fig. 3C**). To investigate a possible role of the WD40 propeller in substrate recruitment, we over-expressed a WDR26-mutant lacking its WD40 domain, together with HBP1 in HEK-293T cells. Interestingly, this mutant was unable to degrade HBP1 (**Fig. 3D**), suggesting that the WD40 domain of WDR26 is functionally relevant *in vivo*. Taken together, these data demonstrate that the HBP1-degrading hGID complex composed of MAEA, RMND5a, TWA1, WDR26 and RanBP9 forms a ring-like, tetrameric structure, possibly stabilized by interactions with the WD40 domains of WDR26.

### ARMC8α but not ARMC8β recruits GID4 to the core complex, but does not prevent binding of the WDR26/RanBP9 module

Previous cryo-EM structural analysis of the yeast Gid4-containing GID complex (Gid1, Gid2, Gid4, Gid5, Gid8 and Gid9) lacking WDR26/Gid7 revealed a monomeric assembly in which Gid4 binds Gid5, the yeast ARMC8 homologue (Qiao *et al*, 2019). Consistently, human GID4 also requires ARMC8 to be recruited into the GID complex *in vitro* (**Suppl. Fig. 3A**). Mammalian cells express two main ARMC8 isoforms, ARMC8α (residues 1-673) and ARMC8β (residues 1-385) (**Fig. 4A**), which are both expressed at comparable levels in HEK-293T cells (**Fig. 4B**). Interestingly, ARMC8β lacks the conserved C-terminal domain, which in yeast Gid5 has been implicated in Gid4 binding (Qiao *et al*, 2019) (**Suppl. Fig. 3B**). This suggests that ARMC8α, but not ARMC8β, is able to recruit GID4. To test this hypothesis, we first performed immunoprecipitation assays using HSS-tagged WDR26 and FLAG-tagged GID4. While ARMC8α readily co-purified with GID4 complexes, ARMC8β fails to interact with human GID4 *in vivo* (**Fig. 4C**). In contrast, both ARMC8α and ARMC8β co-immunoprecipitate with WDR26, suggesting that their binding does not compete with the WDR26/RanBP9 module.

**Figure 4:**
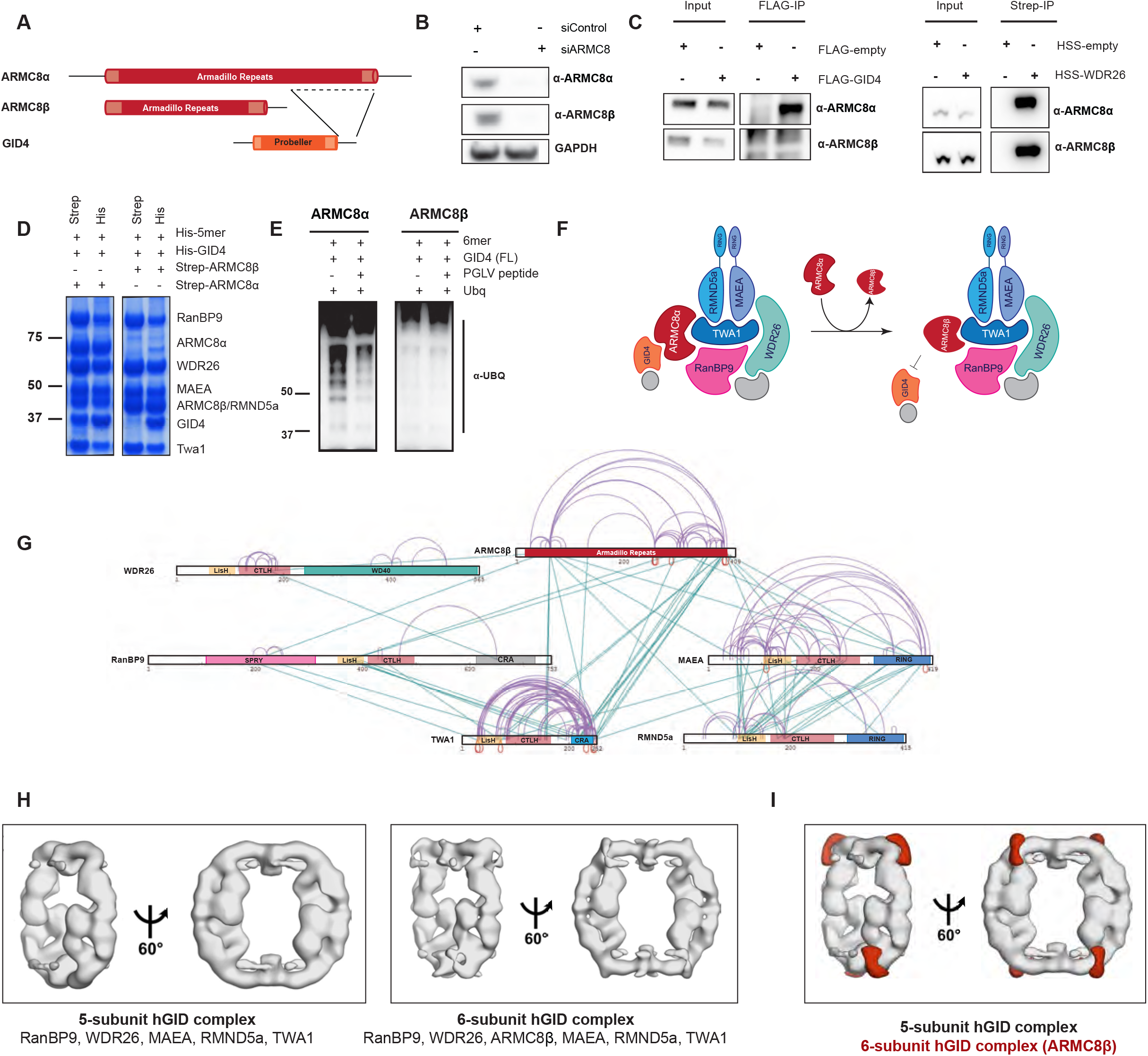
Armc8α but not Armc8β recruits GID4 to the core complex in an assembly that does not prevent binding of the Wdr26/RanBP9 module. **A.** Schematic representation of ARMC8α (FL) (Q8IUR7-1), ARMC8β (Q8IUR7-6) and GID4 (Q8IVV7-1) proteins. **B.** Western blot analysis showing the levels of ARMC8α and ARMC8β in HeLa Kyoto cells after 72 hr treatment with control siRNA or siRNA pools against ARMC8. **C.** Transiently expressed FLAG-GID4 (left panels) or HSS-WDR26 (right panels) was immunoprecipitated from HEK-293T cells and probed by immunoblotting for the presence of ARMC8 isoforms. **D.**Baculoviral co-expression in *Sf9* cells of the 5-subunit hGID complex (5mer; Strep-RanBP9, His-WDR26, FLAG-MAEA, His-RMND5a and His-TWA) with His-GID4 in the presence of ARMC8α or Armc8β Strep- or His-pulldowns revealed the presence of GID4 in ARMC8α but not in Armc8β, complexes. **E.** Immunoblot analysis of *in vitro* ubiquitination of GID4-dependent complexes in the presence of ARMC8α or ARMC8β Where indicated, the reaction was carried out in the presence of 20-fold molar excess of the PGLV GID4-binding peptide. **F.** Schematic representation illustrating that in contrast to ARMC8α, incorporation of ARMC8β prevents hGID activity towards GID4 substrates. **G.** XL-MS analysis of the 6-subunit hGID complex (RanBP9, WDR26, RMND5a, MAEA, TWA1 and ARMC8β). Cross-links within the different complex subunits are indicated by green lines, and cross-links within the same subunit by purple lines. The predicted domain boundaries within the different subunits are colored as follows: LisH domain in light orange, CTLH domain in dark orange, RING domains in blue, TWA1’s CRA domain in light blue, WD40 in dark cyan, ARMC8β in dark red and the SPRY domain in light magenta. **H.** Comparison of the cryo-EM maps of the 5-subunit hGID complex (RanBP9, WDR26, RMND5a, MAEA and TWA1) and the 6-subunit hGID complex (RanBP9, WDR26, RMND5a, MAEA, TWA1 and ARMCβ). **I.** The difference map (red) shows the extra density corresponding to ARMC8β

To directly test assembly of these ARMC8 isoforms with GID4 and other members of the GID core complex *in vitro*, we reconstituted hGID complexes containing either ARMC8α or ARMC8β (**Fig. 4D**). Indeed, while both ARMC8α and ARMC8β readily integrate into the complex, GID4 was only present in ARMC8α-containing complexes (**Fig. 4D**). Consistent with this observation, ARMC8β-containing hGID complexes showed a prominent reduction in GID4-dependent ubiquitination activity compared to ARMC8α controls (**Fig. 4E**). Finally, purified ARMC8α, but not ARMC8β, was able to bind GID4 *in vitro* (**Suppl. Fig. 3C**). Taken together, these results suggest an isoform-dependent regulation of hGID activity, where ARMC8β-bound hGID is not able to bind the GID4 substrate receptor and has reduced ubiquitination activity towards GID4 substrates (**Fig. 4F**).

To gain deeper molecular insights into the ARMC8β-containing hGID complex, we analyzed the 6-subunit hGID assembly (MAEA, RMND5a, TWA1, WDR26, RanBP9 and ARMC8β) by XL-MS and single particle cryo-EM. Size exclusion purification of this complex by Superose 6 column showed one main peak, indicative of a stable complex of similar size as compared to the 5-subunit complex lacking ARMC8β (**Suppl. Fig. 3D**). ARMC8β showed prominent cross-links with the C-terminal CRA domain of TWA1 and RanBP9’s LisH and CTLH domains (**Fig. 4G**). ARMC8β also connects to MAEA and RMND5a by several cross-links, and forms a dense network of cross-links within the core subunits and RanBP9, suggesting that it closely binds and stabilizes these subunits. Oligomerization was further supported by cross-links between the same lysine residues in MAEA, ARMC8β and TWA1. Indeed, cryo-EM demonstrated that ARMC8β-containing hGID complexes maintain the tetrameric ring-like architecture of the 5-subunit hGID complex (**Fig. 4H**, **Suppl. Fig. 3E**). However, ARMC8β-containing hGID complexes appeared more rigid with an extra density near the interface of the subunits, suggesting that ARMC8β stabilizes the oligomeric assembly (**Fig. 4I**, **Suppl. Fig. 3F**).

Identifying the position of ARMC8β in the hGID assembly (**Fig. 4I**, **and Fig. 5A**), and fitting a hGID homology model based on the yeast structure, facilitated the assignment of the remaining GID subunits and domains, such as RanBP9 and TWA1, in the cryo-EM map of the complex (**Fig. 5B**). We generated homology models for ARMC8β, RanBP9 (SPRY and LisH domain) and TWA1 (LisH, CTLH, and CRA domains) based on the structure of Gid1 and homology modelling of Gid8, respectively (**Fig. 5B**). Consistent with the XL-MS data (**Fig. 3B and Fig. 4G)**, RanBP9 and TWA1 mediate major interactions via the LisH and CTLH/CRA domains, respectively (**Fig. 5C**). Furthermore, the SPRY domain of RanBP9 approaches the WD40 domain of WDR26, as also confirmed by several cross-links between these domains (**Fig. 3B**, **Suppl. Fig. 4A**). Moreover, in our fitted model, the WD40 domain of WDR26 is positioned far from ARMC8β and the RING module, suggesting that WDR26 does not contact these subunits directly. Based on the yeast GID structure, MAEA (homologue of Gid9) localizes next to TWA1 (**Fig. 5B** **and** **C**), which places the catalytic RING module (MAEA and RMND5a) at the second dimerization interface. *In vitro* pull-down assays did not show any direct interaction between the RING module (MAEA and RMND5a) and ARMC8-GID4 nor with RanBP9 (**Suppl. Fig. 4B** **and** **C**). Rather, TWA1 was necessary to link RanBP9 and ARMC8-GID4 to the catalytic module. Finally, fitting the yeast Gid5-Gid4 module into the cryo-EM map of the human GID complex shows no steric clashes between the two substrate recruitment subunits, GID4 and WDR26 (**Fig. 5D**). This suggests that the hGID complex may simultaneously engage the two substrate recruitment receptors.

**Figure 5:**
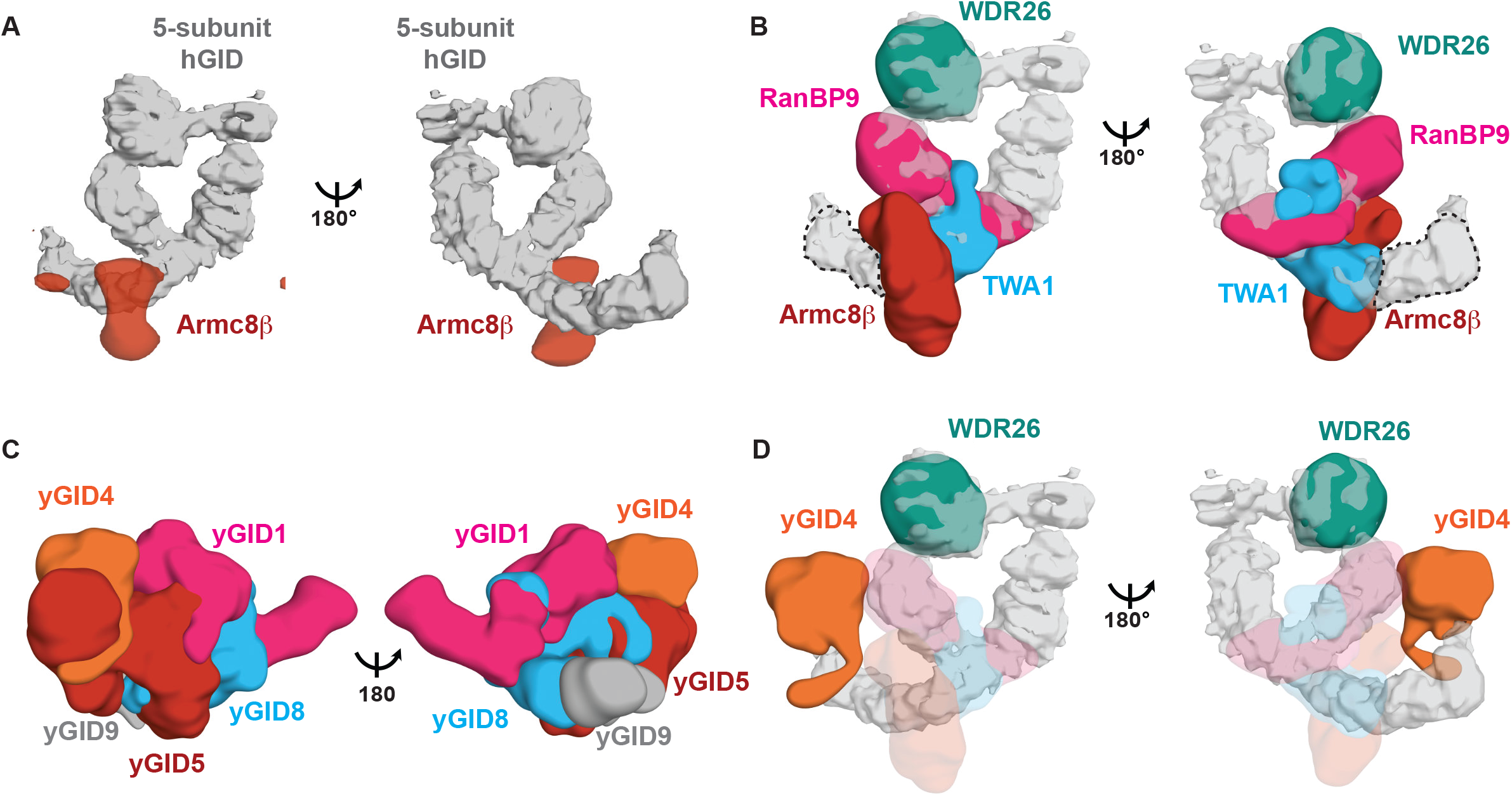
Comparison and organization of the human and yeast GID complexes. **A.** ARMC8β (difference map shown as red surface) binds to the scaffold, distal from the WDR26 WD40 propeller (the 5-subunit hGID map is shown in grey). **B.** Homology models of RanBP9 (SPRY and LisH domains) (magenta), TWA1 (blue) and the WD40 domain of WDR26 (green) are shown fitted into the map of the 5-subunit hGID complex. A homology model of ARMC8β fitted into the difference density is shown in red. The approximate position of the RING-domain containing subunits, MAEA and RMND5a, is indicated with a dotted line. **C.** The yeast GID complex was superimposed on the cryo-EM map of the 5-subunit hGID complex. The Gid4 (orange), Gid1 (magenta), Gid8 (blue), Gid5 (dark red) and Gid9 (gray) subunits of the yeast structure are shown in the same orientation as the hGID complex. **D.** Spatial arrangement of yeast Gid4 with respect to WDR26 is shown in context of the hGID complex.

## Discussion

This study provides a molecular framework for how the human GID E3 ligase recruits its substrates. Several multi-subunit E3 ligase complexes use dedicated subunits for catalytic activity and substrate recruitment. For example, Cullin-RING ligases (CRL) engage one out of a large family of substrate receptors, and their assembly is regulated by substrate availability and the exchange factor CAND1 (Pierce *et al*, 2013). Our results identified ARMC8α-GID4 and RanBP9-WDR26 as distinct substrate-recruitment modules of human GID complexes. In addition to Gid4, yeast cells express two alternative substrate receptors, Gid10 and Gid11, which all interact with the GID E3 ligase complex through Gid5/ARMC8 (Melnykov *et al*, 2019, Edwin Kong *et al*, 2021). Gid4 and Gid10 bind substrates containing an N-Pro degron motif (Dong *et al*, 2020), although systematic screening identified many candidates that do not fulfill these criteria (Edwin Kong *et al*, 2021). Similarly, human GID4 may also recognize substrates like ZMYND19 that lack N-Pro degron motifs. Nevertheless, GID4-dependent ubiquitination of ZMYND19 *in vitro* required a functional N-Pro binding pocket, and it will thus be interesting to determine how this substrate class is recognized. Using bioinformatic criteria, no mammalian GID4-like substrate receptors have been detected, and it may thus be interesting to screen for ARMC8α-interacting proteins to expand the substrate-receptor family.

We previously found that WDR26/Gid7 regulates the cell cycle by targeting the tumor suppressor HBP1 (Lampert *et al*, 2018). Indeed, WDR26 is overexpressed in many human tumors, and, intriguingly, our results suggest that overexpression is sufficient to trigger HBP1 degradation. This activity requires its WD40 domain, which may be involved in substrate recognition, analogous to DCAF substrate adaptors of CRL4 (Angers *et al*, 2006). Interestingly, the cell cycle function of WDR26 requires RanBP9 and the catalytic core subunits, but not ARMC8α or GID4. Similarly, yeast Gid7 is not necessary to degrade N-Pro substrates (Qiao *et al*, 2019), and thus further work is needed to identify cognate WDR26/Gid7 targets.

Fitting the available yeast GID structure (Qiao *et al*, 2019) into the hGID cryo-EM map confirms that the overall structural fold of the GID complex is conserved between yeast and human. Indeed, biochemical data demonstrates that the hGID E3 ligase complex uses ARMC8-GID4 as a substrate-recognition module. We observed no direct binding of either ARMC8 nor RanBP9 with the catalytic RING-containing subunits, suggesting that the central scaffold TWA1 may bridge these interactions. However, the described yeast GID complex lacks WDR26/Gid7, which we show in the human counterpart directly interacts with RanBP9. Thus, consistent with the *in vivo* data, ARMC8 and RanBP9 may function as adaptors to recruit distinct substrate receptors, WDR26 or GID4, respectively. Unlike CRL complexes, the spatial organization of the hGID complex suggests that both WDR26 and GID4 can be recruited at the same time (**Fig. 5D**), as they interact through distinct surfaces. The hGID complex may therefore function as a single unit with separate substrate recruitment modules, or exist as individual complexes that may favor one substrate recruitment module over the other.

Interestingly, while the yeast GID complex lacking WDR26/Gid7 is monomeric, the human GID complex assembles into a stable tetramer, with WDR26 and two catalytic RING modules forming oligomerization interfaces at both ends. This means that four RING domains may be positioned next to each other. Since hGID tetramers are active, it is possible that the bundled catalytic subunits cooperate with each other to increase poly-ubiquitination of cognate substrates. Alternatively, tetramerization may stabilize hGID complexes and favorably position bound substrates and the catalytic core units to allow efficient ubiquitin transfer from the E2 enzymes. However, the relative assembly and arrangement of the distinct substrate-recruiting modules in the tetramer remains to be explored. Finally, analogous to other multimeric complexes, sequestration of subunits may increase their half-life by protecting against auto-ubiquitination and self-destruction, presumably by burying ubiquitination sites and disordered regions required for proteasomal recognition (Mallik & Kundu, 2018).

Although the functional importance of hGID oligomerization remains unclear, it is interesting to note that similar properties have recently been described for other multi-subunit E3 ligases. For example, DCAF1 promotes oligomerization of CRL4 (Mohamed *et al*, 2021), and the Cul3-BTB adaptor SPOP polymerizes these CRL complexes and drives phase separation in cells (Cuneo & Mittag, 2019). Some E3 ligases are inhibited by oligomerization, while others oligomerize to increase catalytic activity (Balaji & Hoppe, 2020). Thus, further work will be required to understand the mechanism and function of oligomerization of hGID complexes.

Several E3 RING ligases regulate their catalytic activity by posttranslational modifications, such as phosphorylation, as in the cases of c-Cbl (Levkowitz *et al*, 1999), MDM2 (Khosravi *et al*, 1999) and NEDD4 (Debonneville *et al*, 2001). In addition, CRL activity is activated by covalent attachment of NEDD8, which promotes ubiquitin transfer to bound substrates (Duda *et al*, 2008). Interestingly, neddylation also prevents CAND1-mediated exchange of substrate adaptors, which is critical to dynamically assemble the required repertoire of cellular CRL complexes. Here, we uncovered an unconventional mechanism for how hGID complexes regulate their activity towards ARMC8/GID4 or WDR26/RanBP9-dependent substrates. We showed that human GID complexes can prevent GID4 recruitment by incorporating the shorter ARMC8β isoform (**Fig. 4F**). ARMC8β was previously described as an integral part of the hGID complex (Maitland *et al*, 2019; Kobayashi *et al*, 2007), and our results demonstrate that ARMC8β incorporation neither affects the oligomeric state (**Suppl. Fig. 3D)** nor the overall shape (**Fig. 4I**), but rather stabilizes the tetramer. Regulating the cellular levels or assembly of ARMC8α and ARMC8β into the complex may thus alter the stability of GID4 substrates *in vivo*. It will be interesting to determine whether hGID tetramers have variable ARMC8α and ARMC8β ratios and if cellular factors are needed to exchange these stably bound subunits to differentially modulate hGID-dependent substrate degradation.

## Materials and Methods

### Cell culture, immunoprecipitation and western blot experiments

HeLa Koyoto, HEK-293T, and RPE cells were grown in NUNC cell culture dishes in Dulbecco’s modified medium (DMEM) from Invitrogen supplemented with 10% FBS and 1% Penicillin-Streptomycin-Glutamine 100x (PSG, Life Technologies). ON-TARGETplus SMARTpool siRNA reagents targeting corresponding genes (ARMC8 #L-018876-00; hGID4 #L-017343-02; RanBP9 #L-012061-00; WDR26 #L-032006-01; Non-targeting Pool #D-001810-10) were purchased from Horizon Discovery. Briefly, HeLa Kyoto cells were transfected with 50nM of siRNA reagents using Lipofectamine 2000 (Thermo Fisher Scientific) according to the manufacturer’s specifications. Cells were harvested after 72h in denaturing urea/SDS buffer, and protein levels of corresponding hGID subunits or HBP1 were detected by immunoblotting.

To co-express HBP1 or ZMYND19 with WDR26, WDR26-ΔWD40 or GID4, 10 cm dishes of HEK-293T cells were transfected with either 6 μg of pcDNA5-HA-Strep-Strep (HSS)-HBP1 or pcDNA5-HSS-ZMYND19 alone, or together with 6 μg pcDNA5-HSS-WDR26, pcDNA5-HSS-WDR26-ΔWD40 or pcDNA5-FLAG-GID4. The media was changed after 14-16h and treated for 10 hours with 5 μM MG132 or for control DMSO. Cells were harvested ~48h post transfection, and lysed in 50 mM Tris-HCl pH 8.0, 150 mM NaCl, 1% NP-40, 0.5% sodium deoxycholate, 0.1% SDS and Complete Protease Inhibitor Cocktail (Roche). Lysates were cleared by centrifugation for 5 mins at 5000 rpm, and protein concentrations were normalized to 1 mg total protein using buffer containing Tris pH 7.7, 200 mM NaCl, and 0.5 mM tris(2-carboxyethyl)phosphine (TCEP).

For immunoprecipitation experiments, lysates were loaded on Strep or Flag beads, and incubated for 1 h at 4 °C. Beads were then washed three times with the lysis buffer (40 mM Tris-HCl pH 7.4, 120 mM NaCl, 1 mM EDTA, 0.3% CHAPS, 1 mM PMSF, 10% Glycerol, 0.5 mM TCEP, 1x PhosSTOP, and 1x Complete Protease Inhibitor Cocktail (Roche)), eluted with SDS-loading dye and incubated 5 mins at 95 °C, before analyzing bound proteins by immunoblotting.

Proteins were resolved by standard SDS-PAGE or NuPAGE 4-12% Bis-Tris Protein Gels (Invitrogen) before transfer onto Immobilon-PVDF or Nitrocellulose transfer membranes (Millipore). Before incubation with the respective primary antibodies, membranes were blocked in 5% milk-PBST (MIGROS) for 1h. For protein detection primary antibodies against ZMYND19 (ab86555, Abcam), HBP1 (11746-1-AP, Protein Tech Group), ARMC8 (sc-365307, SantaCruz), WDR26 (A302-244A, Bethyl Laboratories), TWA1 (5305, Prosci-Inc), MAEA (AF7288-SP, R&D Systems Europe Ltd), RanBP9 (A304-779A, Bethyl Laboratories), FLAG (M2, F3165, Sigma-Aldrich or F7425, Sigma-Aldrich), ubiquitin conjugates (P4D1, sc-8017, Santa Cruz), and GADPH (G-8795, Sigma-Aldrich) were used. Secondary antibodies were goat anti-mouse IgG HRP (170-6516, Bio-Rad), goat anti-rabbit IgG HRP (170-6515, Bio-Rad). Proteins were visualized with SuperSignal™ West Pico PLUS Chemiluminescent Substrate solution (Thermo Fisher) and scanned on a Fusion FX7 imaging system (Witec AG). For re-probing, blots were stripped in ReBlot Plus stripping buffer (2504 Millipore), and washed several times in PBST.

### *Sf9* protein expression and purification

cDNAs encoding human ARMC8⍺, ARMC8β, RanBP9, TWA1, MAEA, RMND5a, HBP1, GID4, ZMYND19, and WDR26 (121-661), were cloned into pAC8 vector, which is derived from the pBacPAK8 system (ClonTech). Recombinant baculoviruses were prepared in *Spodoptera frugiperda* (*Sf9*) cells using the Bac-to-Bac system (Life Technologies). Recombinant protein complexes were expressed in *Spodoptera frugiperda* by co-infection of single baculoviruses. For the 5-subunit hGID complex (RanBP9, MAEA, RMND5a, WDR26, and TWA1), RanBP9 was expressed with N-terminal Strep (II) tag, MAEA with N-terminal FLAG tag, and RMND5a, WDR26, and TWA1 with N-terminal His tag. For the 6-subunit hGID complex (ARMC8, RanBP9, MAEA, RMND5a, WDR26, and TWA1), ARMC8⍺ or ARMC8β were expressed with an N-terminal Strep (II) tag, MAEA with N-terminal FLAG tag, RanBP9, and RMND5a, WDR26, and TWA1 with an N-terminal His tag. For the 4-subunit hGID complexes MAEA, RMND5a, WDR26, and TWA1, or MAEA, RMND5a, ARMC8, and TWA1, WDR26 or ARMC8 were expressed with N-terminal Strep (II) tag, MAEA with N-terminal FLAG tag, RMND5a and TWA1 with N-terminal His tag. Full-length HBP1 was expressed with an N-terminal glutathione S-transferase (GST) tag, and ZMYND19 and GID4 with an N-terminal Strep (II) tag. Cells were harvested 36-48 h after infection and lysed by sonication in Tris-HCl pH 7.7, 200 mM NaCl, 0.5 mM TCEP, including 0.1% Triton X-100, 1x protease inhibitor cocktail (Roche Applied Science) and 1 mM phenylmethanesulfonyl fluoride (PMSF). Lysates were cleared by ultracentrifugation for 45 min at 40,000 *g*. The supernatant was loaded on Strep-Tactin (IBA life sciences) affinity chromatography beads in buffer containing Tris-HCl pH 7.5, 200 mM NaCl and 0.5 mM TCEP. The Strep (II) elution fractions were further purified via ion exchange chromatography (Poros HQ 50 μm, Life Technologies) and subjected to size-exclusion chromatography in a buffer containing 50 mM HEPES pH 7.4, 200 mM NaCl and 0.5 mM TCEP. For GID4 and HBP1, 10% of glycerol was added to all buffers. GID4 was purified by size exclusion chromatography in a buffer containing 50 mM MES pH 6.5, 200 mM NaCl and 0.5 mM TCEP. Pure fractions, as judged by SDS-PAGE, were collected and concentrated using 10,000 MWT cut-off centrifugal devices (Amicon Ultra) and stored at −80°C.

### Size Exclusion Chromatography-Multi-Angle Light Scattering (SEC-MALS)

The oligomeric state of the 5-subunit hGID complex (RanBP9, WDR26, MAEA, RMND5a and TWA1) was investigated by multiangle light scattering (MALS) coupled with size exclusion chromatography (SEC). SEC was performed on an Agilent 1200 HPLC system equipped with a diode array detector (DAD) using a Superose 6 10/300 column (Cytiva) in 50 mM HEPES pH7.4, 200 mM NaCl and 1mM TCEP. Data from the DAD and miniDAWN Treos-II (Wyatt Technology) were processed with the Astra V software to determine the weight averaged molar mass of the protein complex in the main eluting peak, where the calculated protein extinction coefficient of 1000 ml/(g cm) and the average protein dn/dc of 0.185 ml/g were used.

### *In vitro* ubiquitination and pull-down assays

*In vitro* ubiquitination assays were performed by mixing 0.35 μM hGID complexes and 0.2 μM HBP1 or 0.35 μM ZMYND19 with a reaction mixture containing 0.1 μM E1 (UBA1, BostonBiochem), 1 μM E2 (UBCH5a and UBCH5c, or UBE2H, BostonBiochem) and 20 μM Ubiquitin (Ubiquitin, BostonBiochem). Where indicated, 2 μM GID4 and 20 μM of synthetic GID4-binding peptide (PGLV) were added. Reactions were carried out in 50 mM Tris pH 7.7, 200 mM NaCl, 10 mM MgCl_2_, 0.2 mM CaCl_2_, 3 mM ATP, 2 mM DTT, 0.1x TritonX, 10% glycerol, and 0.1 mg ml^−1^ BSA, and incubated for 120 min at 35 °C. Reactions were stopped with SDS loading dye, and analyzed by western blot using anti-HBP1 (11746-1-AP, Protein Tech Group, 1:500) or anti-ZMYND19 antibody (ab86555, Abcam, 1:500).

For GID4-dependent *in vitro* ubiquitination reactions, 0.35 μM hGID complexes (RANBP9, MAEA, RMND5a, WDR26, and TWA1) with either ARMC8⍺ or ARMC8β were mixed with 0.2 μM ZMYND19 and 2 μM GID4, in the presence or absence of 40 μM GID4-binding synthetic peptide (PGLV). Reactions were carried out in 50 mM Tris pH 7.7, 200 mM NaCl, 10 mM MgCl_2_, 0.2 mM CaCl_2_, 3 mM ATP, 2 mM DTT, 0.1x TritonX, 10% glycerol, and 0.1 mg ml^−1^ BSA, and incubated for 120 min at 33 °C. Reactions were then analyzed by western blot using anti-Ubiquitin (P4D1) primary antibody (Santa Cruz).

For pull-down assays in *Sf9 cells*, 100 μL of baculoviruses of the 5-subunit hGID complex: Strep-RanBP9, His-WDR26, FLAG-MAEA, His-RMND5a, and His-TWA, with His-GID4 and His-ARMC8α or His-Armc8β were co-infected in 10 ml of *Sf9* cells. Infected cells were incubated at 27 °C for 48 h, and lysed by sonication in Tris-HCl pH 7.7, 200 mM NaCl, 0.5 mM TCEP, including 0.1% Triton X-100, 1x protease inhibitor cocktail (Roche Applied Science), and 1 mM PMSF. Lysates were cleared by centrifugation at 14,000 *g* for 30 minutes, and 1 ml of soluble protein fractions were incubated for 1 h at 4 °C with 20 μL Strep-Tactin Macroprep beads (IBA lifesciences). Beads were washed three times with lysis buffer, and bound proteins were eluted in 20 μL of SDS loading dye and heated at 95 °C for 2 min.

### Cross-linking mass spectrometry

Two different cross-linking protocols were used in this work, based on the amine-reactive disuccinimidyl suberate (DSS) (Leitner *et al*, 2013) and a combination of pimelic dihydrazide (PDH) and the coupling reagent 4-(4,6-dimethoxy-1,3,5-triazin-2-yl)-4-methylmorpholinium (DMTMM) chloride (Mohammadi *et al*, 2021; Leitner *et al*, 2014). DSS was obtained as a 1:1 mixture of “light” (d_0_) and “heavy” (d_12_) isotopic variants from Creative Molecules, Inc. Light (d_0_) PDH was obtained from ABCR, heavy (d_10_) PDH from Sigma-Aldrich; DMTMM chloride was also obtained from Sigma-Aldrich.

Cross-linking conditions were optimized in screening experiments on the 5-subunit hGID complex using SDS-PAGE as a readout and 1 mM DSS (d_0_/d_12_) and 22 mM PDH (d_0_/d_10_) + 4.4 mM DMTMM were selected as the optimal conditions. The low concentration of DMTMM relative to PDH results in the dominant formation of zero-length cross-links over the integration of the dihydrazide linker (Mohammadi *et al*, 2021). For XL-MS, protein complexes were prepared at a total protein concentration of 1 mg/ml in a buffer containing 50 mM HEPES pH 7.4, 200 mM NaCl, 1mM TCEP and cross-linked at 50 μg scale. DSS cross-linking was performed at 37 °C for 30 min, followed by a quenching step (50 mM NH_4_HCO_3_) for 30 min at the same temperature. PDH+DMTMM cross-linking was performed for 45 min at 37 °C followed by removal of the reagents by gel filtration (Zeba spin desalting columns, ThermoFisher Scientific).

After quenching or gel filtration, samples were dried in a vacuum centrifuge and redissolved in 8 M urea solution for reduction (2.5 mM tris-2-carboxyethyl phosphine, 37 °C, 30 min) and alkylation (5 mM iodoacetamide, 23 °C, 30 min in the dark) steps. Samples were then diluted to ~5.5 M urea with 150 mM NH_4_HCO_3_ before addition of endoproteinase Lys-C (Wako, 1:100, 37 °C, 2 h), followed by a second dilution step to ~1 M urea with 50 mM NH_4_HCO_3_ and addition of trypsin (Promega, 1:50, 37 °C, overnight). After overnight incubation, samples were acidified to 2% (v/v) formic acid and purified by solid-phase extraction (SepPak tC18 cartridges, Waters). Purified samples were fractionated by peptide-level size-exclusion chromatography (SEC) (Leitner *et al*, 2013; 2012)) using Superdex Peptide PC 3.2/300 (for the 5-subunit hGID complex) or Superdex 30 Increase 3.2/300 (for the 6-subunit hGID complex) columns (both GE Healthcare). Three high-mass fractions enriched in cross-linked peptide pairs were collected for MS analysis.

Liquid chromatography-tandem mass spectrometry (LC-MS/MS) was performed on an Easy nLC 1200 HPLC system connected to an Orbitrap Fusion Lumos mass spectrometer (both ThermoFisher Scientific). Peptides were separated on an Acclaim PepMap RSLC C_18_ column (250 mm × 75 μm, ThermoFisher Scientific). The LC gradient was set from 9 to 40% mobile phase B in 60 min, mobile phases were A = water/acetonitrile/formic acid (98:2:0.15, v/v/v) and B = acetonitrile/water/formic acid (80:20:0.15, v/v/v), and the flow rate was 300 nl/min.

Each SEC fraction was injected in duplicate with two different data-dependent acquisition methods for MS analysis. Both methods used a top-speed method with 3 s cycle time and detection of precursors in the Orbitrap analyzer at 120000 resolution. Precursors were selected for fragmentation if they had a charge state between 3+ and 7+ and an m/z between 350 and 1500, and were fragmented in the linear ion trap at a normalized collision energy of 35%. The high-resolution method used detection of the fragment ions in the Orbitrap at 30000 resolution; the low-resolution method used detection in the linear ion trap at rapid scan speed. The two different methods were selected to benefit from either the higher mass accuracy of Orbitrap detection or the higher sensitivity of ion trap detection. xQuest (version 2.1.5, available from https://gitlab.ethz.ch/leitner_lab/xquest_xprophet (Walzthoeni *et al*, 2012; Leitner *et al*, 2013)) was used to identify cross-linked peptide pairs. MS/MS spectra were searched against custom databases containing the target protein sequences and contaminant proteins and their randomized entries. Important search parameters included: Enzyme specificity = trypsin (no cleavage before P) with maximum two missed cleavages, precursor mass tolerance = 15 ppm, fragment mass tolerance = 15 ppm for Orbitrap detection or 0.2/0.3 Da (common/cross-link ions) for ion trap detection. Oxidation of Met was selected as a variable modification, carbamidomethylation of Cys as a fixed modification. DSS was assumed to react with Lys or the protein N termini; PDH was assumed to react with Asp and Glu; DMTMM was assumed to react with Lys and Asp or Lys and Glu. Primary search results were filtered with a more stringent error tolerance (−5 to +1 ppm for the 5-subunit hGID complex, 0 to +5 ppm for the 6-subunit hGID complex), and were required to have xQuest deltaS scores ≤0.9 and TIC scores ≥0.1 (DSS) or ≥0.15 (DMTMM). The remaining spectra were manually evaluated to have at least four bond cleavages in total per peptide or three consecutive bond cleavages per peptide. Ambiguous identifications containing peptides that could be mapped to more than one protein (from tags) were removed. Finally, an xQuest score cut-off was selected so that the false positive rate was at 5% or less at the non-redundant peptide pair level. All cross-link identifications are provided in SI Tables 1, 2, 4, and 5. The mass spectrometry proteomics data have been deposited to the ProteomeXchange Consortium (http://proteomecentral.proteomexchange.org) via the PRIDE partner repository (Perez-Riverol *et al*, 2019) with the dataset identifier PXD024822. XL-MS data in Figures 3B and 4H were visualized with xiNET (Combe *et al*, 2015).

### Sample preparation and cryo electron microscopy analysis

In order to increase the stability of the 5-subunit and the 6-subunit hGID complexes, the gradient fixation (GraFix) protocol was applied (Stark, 2010). Briefly, samples were loaded on a glycerol gradient (10%-40% w/v) in the presence of the cross-linker glutaraldehyde (0.25% v/v), followed by ultracentrifugation (SW40Ti rotor) at 35,000 rpm for 18 h at 4 °C. Peak fractions containing the protein complexes were collected and buffer exchange for glycerol removal was performed by Zeba Spin columns in a buffer containing 50 mM HEPES pH 7.4, 200 mM NaCl, 1mM TCEP and 0.01% NP40 for the 5-subunit hGID complex, and 0.05% NP40 for the 6-subunit hGID complex. 4 μL sample (0.08-0.15 mg/ml) was then spotted on glow discharged Quantifoil holey grids (R2.2, Cu 300 mesh, Quantifoil Micro Tools GmbH, Grosslöbichau, Germany) after floating them with continuous 1 nm carbon film. Grids were incubated for 20-60 s at 4 °C and 100% humidity, blotted for 1 s with Whatman no.1 filter paper and vitrified by plunging into liquid ethane (Vitrobot, ThermoFischer).

#### Data collection

Three datasets of GID pentamer and one dataset of GID-ARMC8β complexes were collected with the Titan Krios cryo-electron microscope (Thermo Fisher Scientific Inc., Waltham MA) operated at 300kV, using the K2 and K3 direct electron detectors (Gatan Inc., Pleasanton CA), operated in counting or super-resolution mode. Data collection parameters are compiled in SI Table 3.

### Cryo-EM Data analysis of the 5-subunit hGID map

Data acquisition and preprocessing: All micrographs were drift corrected with MotionCor2 using a 5 by 5 patch (Li *et al*, 2013). In addition, micrographs recorded on the K3 detector in super-resolution mode were binned twofold with MotionCor2. Defocus of the drift corrected averages was determined by CTF fitting with Gctf (Zhang, 2016). For each dataset, particles from 10 micrographs representative of the defocus range of the entire dataset were manually selected. The manually selected particle positions were used to train a neural network in order to select particle of the entire dataset with crYOLO (Wagner *et al*, 2019). A total of 815538 particles were selected (88564 from dataset 1, 538734 from dataset 2, 188240 from dataset 3). Accuracy of automated particle selection was verified by manual inspection of particle positions.

2D Classification (GID pentamer): Image processing was carried out in Relion 3.1 (Scheres, 2016). Particles from datasets 1, 2, and 3 were extracted (box size 720, scaled to 96 pixels, resulting pixel size 6.3 Å/pixel) and combined into a single file with 815538 particles. Particles were subjected to two rounds of 2D classification into 100 classes. After the first round, 569845 particles (69%) were selected, rejecting obvious junk classes (ice blobs, edges). The selected particles were subjected to 2D classification in a second round with 429682 particles selected (75%) after removal of junk classes and obviously broken particles. The selected particles were re-extracted with a box size of 720 pixels, scaled to 180 pixels (resulting pixel size: 3.38 Å/pixel), and recentered by application of shifts applied during classification.

Initial model generation: An initial model of GID pentamer was generated in cryoSPARC (Punjani *et al*, 2017). Particles selected from dataset 2 and 3 of GID pentamer complex were extracted with a box size of 640 pixels and binned to 128 pixels (pixel size: 3.36 Å/pixel). After one round of 2D classification, junk classes (ice blobs, edges) were discarded and the remaining particles were used for initial model generation with 3 classes. The initial model generation without application of symmetry or with C2 symmetry resulted in ring shaped reconstruction with a strong density for one half of the ring with twofold symmetry. The application of D2 symmetry resulted in a ring-shaped reconstruction that matched the map for the initial models calculated with C1 and C2 symmetry and showed projections corresponding to the ring-shaped class averages. This model was used as an initial model for heterogeneous refinement into three classes resulting in three models that were very similar, one of which was chosen as initial model for further processing.

#### 3D structure refinement

Particles were 3D classified into 10 classes without application of symmetry, using a model generated with cryoSPARC as initial model. The reconstruction of class 10 showed the hallmark D2 symmetry of the intact GID pentamer complex. Class 10 contained 39255 particles, which corresponded to ∼9.1% of all particles that entered classification, and refined to a resolution of 19.5 Å when refined with application of D2 symmetry. After another step of re-centering and subsequent 2D classification (36134 particles selected, 92%), the particles were refined with application of D2 symmetry to a resolution of 17.3 Å. The refined particles were D2 symmetry expanded using the relion_particle_symmetry_expand function, resulting in 144536 asymmetric unit particles. One asymmetric unit with additional density at the edges (including the second WDR26 beta-propeller) was carved from the refined map using Chimera volume eraser to create a soft-edged mask. The mask, the map, and the expanded particles were all re-centered to the center of mass of the map. The symmetry expanded, recentered particles were subjected to 3D classification. The class that showed detailed structural features in agreement with the map calculated before symmetry expansion contained 33929 ASU particles, and was subjected to a 3D refinement to result in a 9 Å resolution map (FSC 0.143 criterion).

### Cryo-EM Data analysis of the 6-subunit hGID map (The 5-subunit GID and ARMC8β complex)

In order to localize ARMC8β in the GID complex, a difference map between the GID pentamer and the GID-ARMC8β complex was calculated. Drift correction of micrographs were performed with MotionCorr (Li *et al*, 2013) and defocus of the drift corrected averages was determined by CTF fitting with Gctf (Zhang, 2016), resulting in a data set of dataset 3048 micrographs. Particles from 10 representative micrographs were manually selected and used to train a neural network in order to pick particles of the remaining dataset with crYOLO (Wagner *et al*, 2019). A total of 73559 particles of GID-ARMC8β were selected and accuracy of automated particle selection was verified by manual inspection. A combined set of particles from datasets 2 and 3 of GID pentamer was used to calculate a map for comparison, undergoing identical processing steps as the GID-ARMC8β data. Particles were extracted and binned to the same pixel size of 8.4 Å/pixel (GID pentamer: 815538 particles, box size 640 pixels, scaled to box size of 64 pixels, GID-ARMC8β complex: 73559 particles, box size 504 pixels, scaled to a box size of 64 pixels). Both sets were subjected to three rounds of 2D classification into 100 classes where obvious junk classes showing ice contaminations or carbon edges were removed. The GID pentamer particle set was reduced to 197899 particles, GID-ARMC8β to 25753 particles. The GID pentamer dataset was randomly split and 25753 particles were selected. After re-extraction with a box size of 128 pixels and a pixel size of 4.2 Å/pixel, both particle sets were subjected to a 3D classification into five classes with application of D2 symmetry, using the map of GID filtered to 30 Å as initial model for both classifications. In both classifications only one class (GID pentamer: 14550 particles, GID-ARMC8β: 5419 particles) showed the known structural features of the GID complex. Refinement of particles from selected classes with application of D2 symmetry produced the final maps (GID: 20 Å resolution, GID-ARMC8β: 22 Å resolution). Maps were aligned and difference density was calculated in UCSF Chimera.

### Cryo-EM map interpretation

Models for RanBP9 (172-463), TWA1 (27-238), the WD40 domain of WDR26 (349-547) and ARMC8β (31-407) were obtained using homology modelling in Phyre2 (Mezulis *et al*, 2015) and the crystal structure of the SPRY domain of human RanBP9 (PDB 5JI7 (Hong *et al*, 2016)). The ring shaped WD40 domain of WDR26 was fitted into the cryo-EM map with the Chimera (Pettersen *et al*, 2004) fit command (highest correlation 0.95). For the RanBP9, TWA1 and ARMC8β subunits of the GID complex, a homology model was assembled by superimposing the homology structures on the yeast GID coordinates (PDB 6SWY; Qiao *et al*, 2019). The model was placed in the cryo-EM map based on the elongated shape of the ARMC8β difference density and ARMC8β was rigid body docked into the difference density. Based on the placement of ARMC8β, TWA1/RanBP9 was separately fitted as a rigid body into the ASU map. Subunit placements were cross-checked with cross-linking MS results. For visualization, surface representations of the domains were filtered to 10 Å. Images were created using PyMOL (PyMOL, version 2.4.0. New York: Schrodinger Inc., 2020).

### Data availability

Cryo-EM map of the 5-subunit and 6-subunit hGID complexes: Electron Microscopy Data Bank. Cross-linking Mass Spectroscopy: All cross-link identifications are provided in SI Tables 1, 2, 4, and 5. The mass spectrometry proteomics data have been deposited to the ProteomeXchange Consortium (http://proteomecentral.proteomexchange.org) via the PRIDE partner repository (Perez-Riverol *et al*, 2019) with the dataset identifier (PXD024822).

## Acknowledgement

We would like to thank Jason Greenwald for performing the MALS analysis of the 5-subunit GID complex, and Anton Khmelinskii for sharing unpublished results. We are grateful to Anna Maria Stier, Halil Bagci and members of the Peter lab for helpful discussions, and Alicia Smith for critical editing. We thank Eliane Züger for help with the ARMC8 isoforms, and Mahshid Gazorpak and the ETH block course students for initial overexpression experiments. We thank ScopeM and in particular Miroslav Peterek for microscopy training and support. We acknowledge Ruedi Aebersold and Paola Picotti for access to the Orbitrap Fusion Lumos mass spectrometer, which was funding by the ETH Scientific Equipment program and the European Union Grant ULTRA-DD (FP7-JTI 115766). Sophia Park profits from an ITN network grant from the European Research Commission. This work was supported by the Swiss National Science Foundation, the Swiss Cancer League, a donation from Dr. Walter Fischli and the ETH Zürich.

## Author contributions

W.I.W. and M.P. conceptualized the study. W.I.M performed the biochemical assays. W.I.M. and S.P. performed the cellular assays. W.I.M prepared the specimens for EM data collection, J.R., D.B. and W.I.M. imaged EM grids, and J.R. and D.B. processed the EM data. A.L. performed the XL-MS analysis. M.P. and W.I.M. wrote the manuscript, with critical input from all authors.

---

**Supplementary Figure 1:**
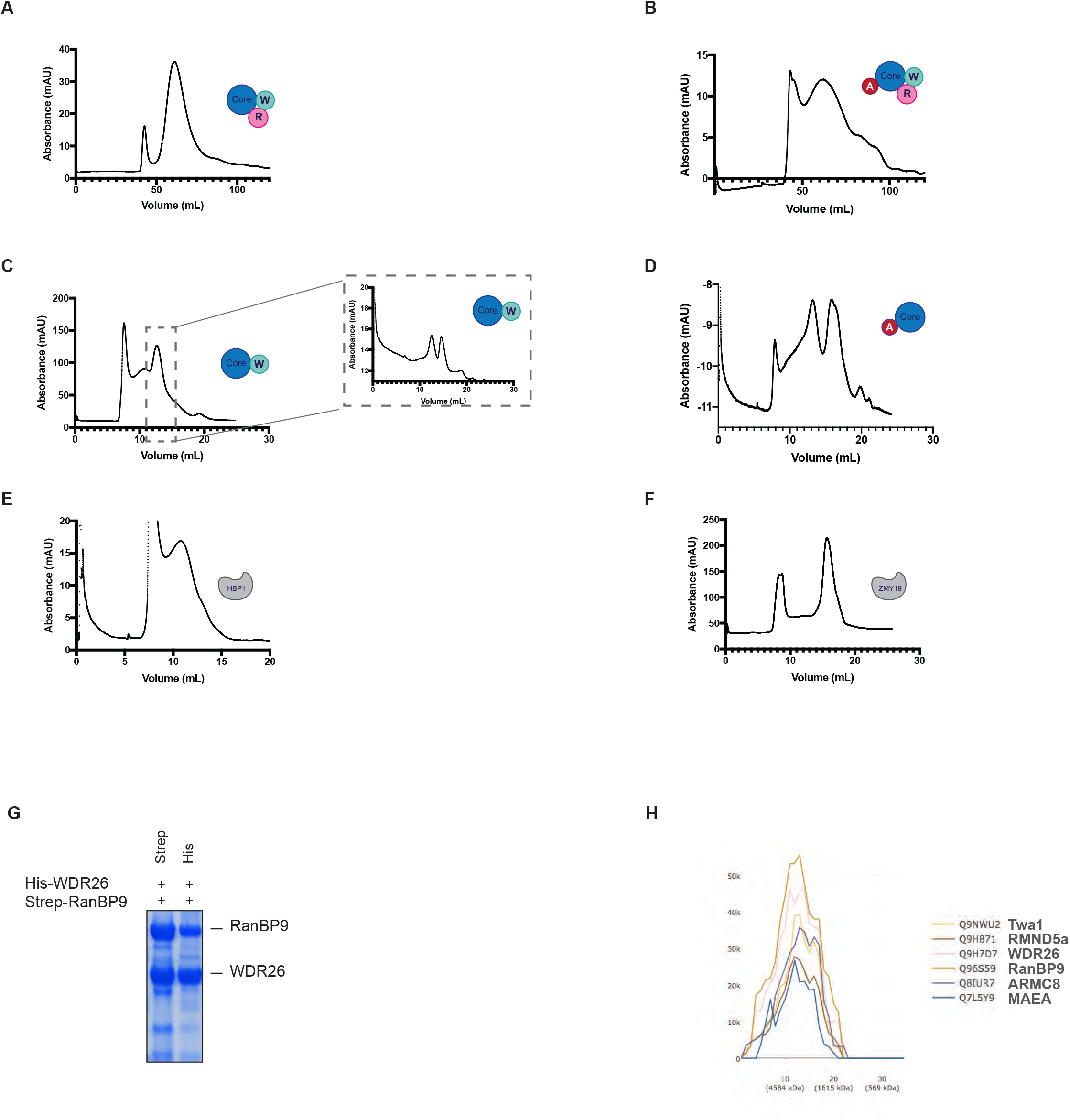
**A to F.** Size exclusion profiles of the indicated hGID sub-complexes. The catalytic core unit composed of MAEA, RMND5a and TWA1 is colored in blue, WDR26 in green, RanBP9 in light magenta, ARMC8 in dark red, and ZMYND19 and HBP1 in gray. **G.** *In vitro* pull-down assay of Strep-RanBP9 and His-WDR26 co-expressed in baculoviral *Sf9* cells, demonstrating the formation of a stable complex. **H.** Size-exclusion profiles of the different endogenous hGID subunits in HeLa cells analyzed by the SEC-Explorer web platform (Heusel *et al*, 2019).

**Supplementary Figure 2:**
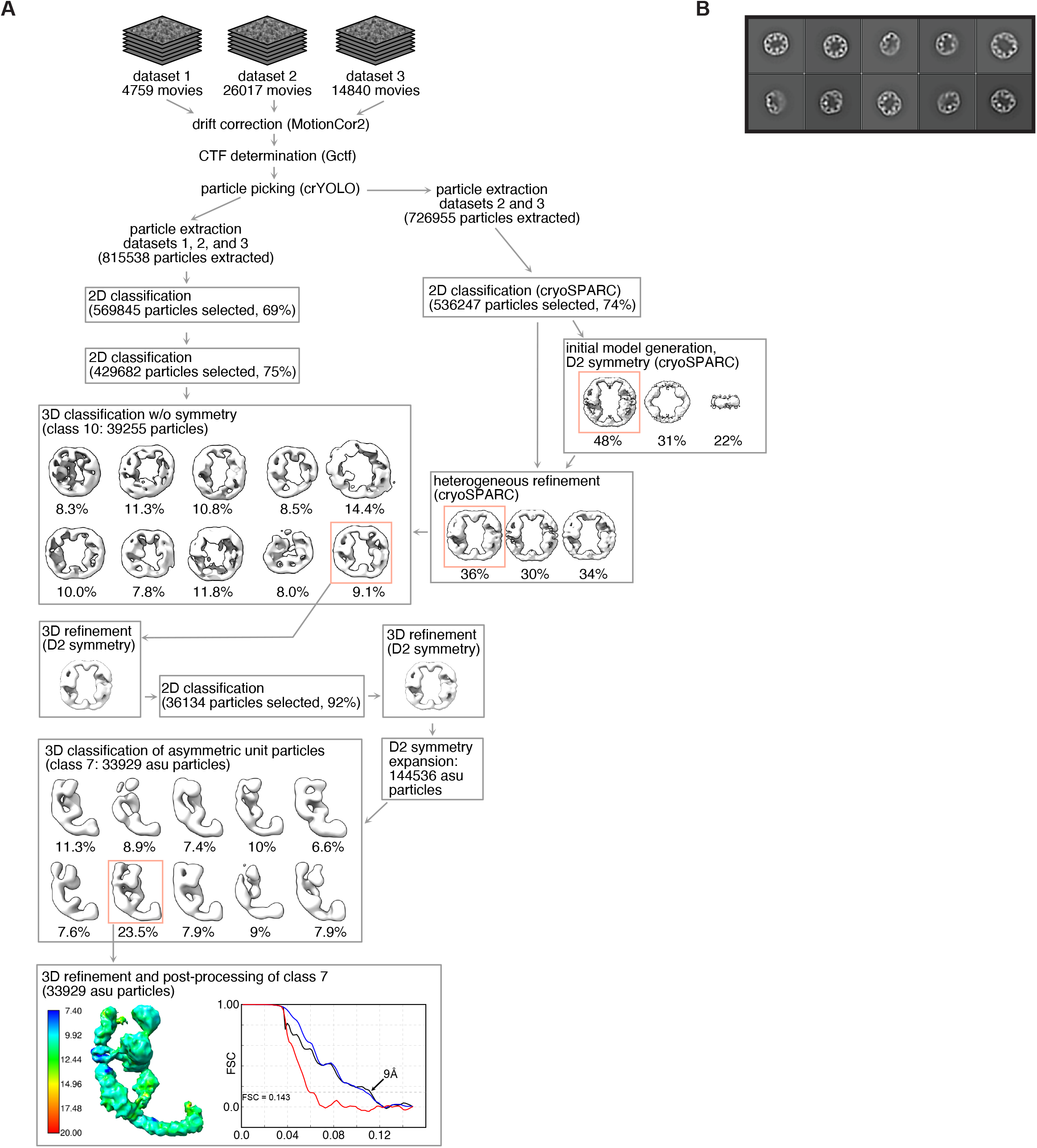
**A.** Single particle image processing of GID pentamers. Three data sets of five subunit GID complexes were combined and 816k particle images extracted. An initial model with D2 symmetry was obtained with CryoSPARC. Several cycles of 2D and 3D classification were required to obtain a homogeneous set of particles for symmetry expansion in Relion. 3D classification after symmetry expansion provided a set of particles that was refined to sub-nanometer resolution. The local resolution map shows that the resolution of the domains extending towards the center of the ring is lower probably due to higher flexibility. The FSC plot shows the masked (blue), masked corrected (black) and phase randomized mask FSC (red). **B.** 2D class averages of 5-subunit hGID complex. The 10 most populated classes (of 100) are shown, ordered by occupancy. The boxes shown are 604 Å across.

**Supplementary Figure 3:**
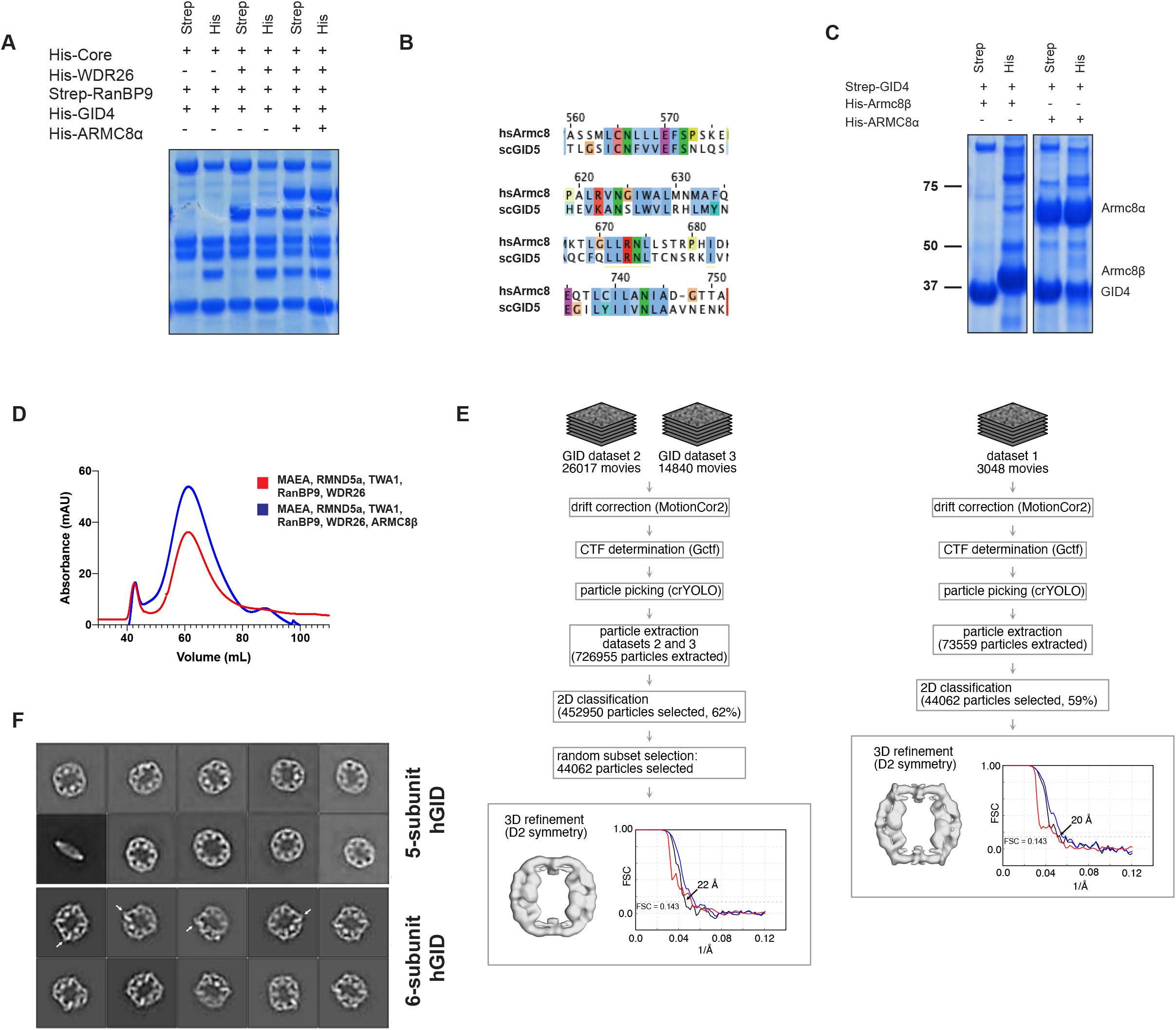
**A.** *In vitro* pull-down assay of His-GID4 and His-ARMC8α from baculoviral *Sf9* extracts co-expressing Strep-RanBP9, His-WDR26, FLAG-MAEA, His-RMND5a and His-TWA1. ARMC8α is required to recruit GID4 into the hGID complex. **B.** Conservation between human ARMC8α and yeast Gid5 at the region required for GID4 binding (Waterhouse *et al*, 2009). **C.** Baculoviral co-expression in *Sf9* cells of Strep-GID4 and His-ARMC8α or His-ARMC8β Note that ARMC8α but not ARMC8β forms a stable complex with GID4. **D.** Comparison of the size-exclusion chromatograms from Superose 6 column (Cytiva) of the 5-subunit (RanBP9, WDR26, RMND5a, MAEA and TWA1) and the 6-subunit (RanBP9, WDR26, RMND5a, MAEA, TWA1 and ARMC8β) hGID complexes. **E.** Single particle image processing scheme used to determine the difference map between 5-subunit and ARMC8β-containing 6-subunit GID complexes. The processing steps of the GID 5-subunit (left) and GID-ARMC8β 6-subunit complexes (right) were carried out with identical settings. The FSC plot shows the masked (blue), masked corrected (black) and phase randomized mask FSC (red). **F.** 2D class averages of the GID pentamer (top) and GID-ARMC8β complexes (bottom). The 10 most populated classes (of 100) are shown, ordered by occupancy. Boxes are 530 Å across. Additional density is visible in the 2D classes of the 6-subunit hGID corresponding to ARMC8β (white arrows).

**Supplementary Figure 4:**
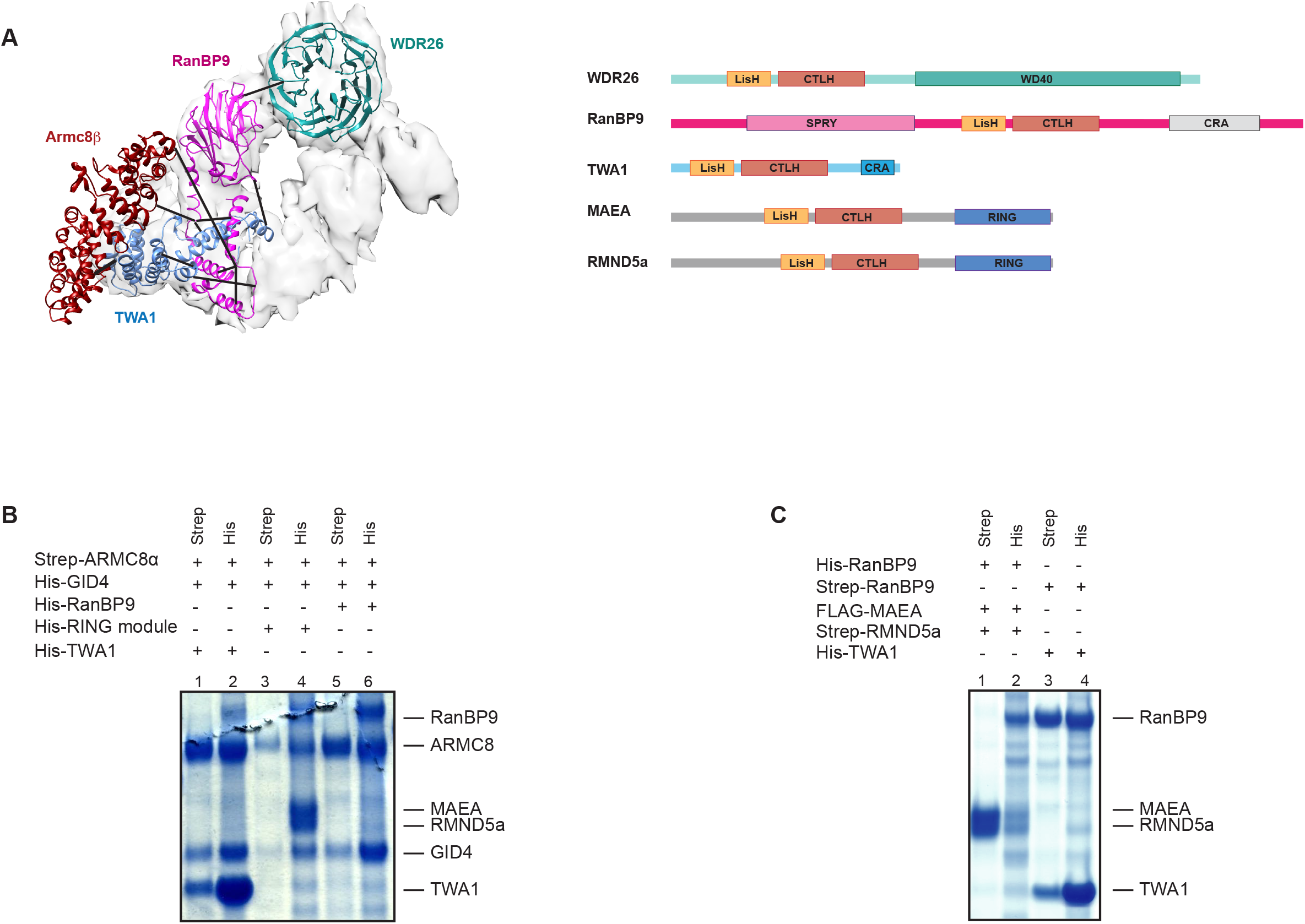
**A.** The fitted homology model of RanBP9 (SPRY and LisH domain) is in magenta, TWA1 is in blue, ARMC8β is in dark red, and WD40 is in dark cyan in the 9 Å map of the 5-subunit hGID complex (gray) (left panel). The observed cross-links between the different residues are indicated by black lines. Schematic architecture and domain representation of the 5 subunits of the hGID complex (RanBP9, WDR26, RMND5a, MAEA and TWA1) are shown to the right. **B.** SDS-PAGE shows *in vitro* pull-down assays by baculoviral co-expression in *Sf9* cells of Strep-ARMC8α which forms a complex with His-TWA1 and His-GID4 (lane 1 and 2) but not with FLAG-MAEA and His-RMND5a (lane 3 and 4). His-RanBP9 does not bind Strep-ARMC8α and His-GID4 (lane 5 and 6). **C.** His-RanBP9 does not interact with FLAG-MAEA and Strep-RMND5a (lane 1 and 2), but Strep-RanBP9 forms a complex with His-TWA1 (lane 3 and 4).

