## Supplementary material for "The human GID complex engages two independent modules for substrate recruitment": Supp. Table 1

**Supplementary table 1**

### High-res MS2

**Cross-linked peptide pairs in the 5mer complex identified using DMTMM as the cross-linking reagent and high-resolution MS/MS detection**

| Id | Protein1 | Protein2 | **XLType** | Spectrum | AbsPos1 | AbsPos2 | deltaAA | Mr | Mz | z | Error_rel[p | TIC | deltaS | **Id-Score** |
|---|---|---|---|---|---|---|---|---|---|---|---|---|---|---|
| HFSQAEGSQLDEVR-VGKAIDK-a6-b3 | pr\|PROT04\|MAEA | pr\|PROT05\|RMND5A | **inter-protein xl** | aleitner_D1 | 252 | 115 | | 2313.167 | 579.3 | 4 | -1.8 | 0.46 | 0 | **35.21** |
| LLNEVMVEHFFR-TKEVFHFSQAEK-a4-b2 | pr\|PROT05\|RMND5A | pr\|PROT04\|MAEA | **inter-protein xl** | aleitner_D1 | 144 | 405 | | 2964.492 | 593.906 | 5 | -1.1 | 0.23 | 0 | **34.51** |
| IQAQIDRFPIGDR-LAKLLK-a6-b3 | pr\|PROT01\|RanBP9 | pr\|PROT03\|Twa1 | **inter-protein xl** | aleitner_D1 | 378 | 218 | | 2194.291 | 439.866 | 5 | -2 | 0.24 | 0.9 | **33.85** |
| HFSQAEGSQLDEVR-TPKDAASVR-a12-b3 | pr\|PROT04\|MAEA | pr\|PROT01\|RanBP9 | **inter-protein xl** | aleitner_D1 | 258 | 221 | | 2527.235 | 632.817 | 4 | -2.5 | 0.19 | 0 | **33.58** |
| ELQAMSEQLR-AKAEWEGK-a7-b2 | pr\|PROT01\|RanBP9 | pr\|PROT02\|WDR26 | **inter-protein xl** | aleitner_D1 | 656 | 172 | | 2103.039 | 702.021 | 3 | -1.5 | 0.25 | 0 | **32.84** |
| HFSQAEGSQLDEVR-DALNKMFNGSK-a12-b5 | pr\|PROT04\|MAEA | pr\|PROT05\|RMND5A | **inter-protein xl** | aleitner_D1 | 258 | 389 | | 2807.331 | 702.841 | 4 | 0.7 | 0.27 | 0 | **32.64** |
| STDQTVLEELASIK-MTDLSKGVIEEPK-a9-b6 | pr\|PROT01\|RanBP9 | pr\|PROT03\|Twa1 | **inter-protein xl** | aleitner_D1 | 424 | 245 | | 2960.519 | 987.847 | 3 | -2.6 | 0.29 | 0 | **32.47** |
| HFSQAEGSQLDEVR-MTDLSKGVIEEPK-a11-b6 | pr\|PROT04\|MAEA | pr\|PROT03\|Twa1 | **inter-protein xl** | aleitner_D1 | 257 | 245 | | 3029.476 | 758.377 | 4 | 0 | 0.22 | 0 | **32.44** |
| HFSQAEGSQLDEVR-LASDHKDIHSSVSR-a11-b6 | pr\|PROT04\|MAEA | pr\|PROT05\|RMND5A | **inter-protein xl** | aleitner_D1 | 257 | 104 | | 3134.504 | 627.909 | 5 | -2.8 | 0.23 | 0 | **31.67** |
| MGEAIETTQQLYPSLLER-MTDLSKGVIEEPK-a17-b6 | pr\|PROT01\|RanBP9 | pr\|PROT03\|Twa1 | **inter-protein xl** | aleitner_D1 | 459 | 245 | | 3505.766 | 877.449 | 4 | -0.6 | 0.28 | 0 | **30.72** |
| QKVWSEVNQAVLDYENR-EGEWQTMIQK-a2-b1 | pr\|PROT03\|Twa1 | pr\|PROT01\|RanBP9 | **inter-protein xl** | aleitner_D1 | 195 | 386 | | 3307.588 | 827.905 | 4 | -1.7 | 0.34 | 0 | **29.71** |
| QKVWSEVNQAVLDYENR-EGEWQTXIQK-a2-b1 | pr\|PROT03\|Twa1 | pr\|PROT01\|RanBP9 | **inter-protein xl** | aleitner_D1 | 195 | 386 | | 3323.578 | 831.902 | 4 | -3 | 0.34 | 0 | **29.23** |
| MSYAEKPDEITKDEWMEK-RLYPAVDEQETPLPR-a6-b10 | pr\|PROT03\|Twa1 | pr\|PROT01\|RanBP9 | **inter-protein xl** | aleitner_D1 | 30 | 186 | | 3993.903 | 799.788 | 5 | -2.5 | 0.24 | 0 | **29.19** |
| ELQAMSEQLRR-AKAEWEGK-a7-b2 | pr\|PROT01\|RanBP9 | pr\|PROT02\|WDR26 | **inter-protein xl** | aleitner_D1 | 656 | 172 | | 2259.135 | 452.835 | 5 | -3.6 | 0.26 | 0 | **27.05** |
| AEWEGK-SKLLDK-a4-b2 | pr\|PROT02\|WDR26 | pr\|PROT02\|WDR26 | **intra-protein xl** | aleitner_D1 | 176 | 185 | 9 | 1402.742 | 468.589 | 3 | -2.5 | 0.48 | 0 | **35.52** |
| DEWMEK-LAKLLK-a1-b3 | pr\|PROT03\|Twa1 | pr\|PROT03\|Twa1 | **intra-protein xl** | aleitner_D1 | 37 | 218 | 181 | 1502.813 | 501.945 | 3 | -2.4 | 0.37 | 0 | **34.81** |
| GVIEEPK-LAKLLK-a4-b3 | pr\|PROT03\|Twa1 | pr\|PROT03\|Twa1 | **intra-protein xl** | aleitner_D1 | 249 | 218 | 31 | 1436.892 | 479.972 | 3 | -2.9 | 0.41 | 0 | **34.44** |
| VKYPKMTDLSK-GVIEEPK-a5-b4 | pr\|PROT03\|Twa1 | pr\|PROT03\|Twa1 | **intra-protein xl** | aleitner_D1 | 239 | 249 | 10 | 2061.112 | 516.286 | 4 | -2.9 | 0.26 | 0 | **34.25** |
| AEWEGKGTASR-YLEYLEDGK-a6-b7 | pr\|PROT02\|WDR26 | pr\|PROT02\|WDR26 | **intra-protein xl** | aleitner_D1 | 178 | 130 | 48 | 2301.086 | 768.036 | 3 | -2.5 | 0.39 | 0 | **34.24** |
| MSYAEKPDEITKDEWMEK-XSYAEKPDEITKDEWMEK-a6-b5 | pr\|PROT03\|Twa1 | pr\|PROT03\|Twa1 | **intra-protein xl** | aleitner_D1 | 30 | 29 | 1 | 4455.96 | 892.2 | 5 | -4 | 0.29 | 0 | **34** |
| RKAVESIQAEDESAK-EVEESLER-a2-b1 | pr\|PROT04\|MAEA | pr\|PROT04\|MAEA | **intra-protein xl** | aleitner_D1 | 107 | 188 | 81 | 2631.293 | 658.831 | 4 | -2.1 | 0.35 | 0 | **33.58** |
| DGYMGIGLSAQGVNMNR-TPKDAASVR-a1-b3 | pr\|PROT01\|RanBP9 | pr\|PROT01\|RanBP9 | **intra-protein xl** | aleitner_D1 | 251 | 221 | 30 | 2707.311 | 903.445 | 3 | -2.3 | 0.47 | 0 | **33.5** |
| VQCLWCLSDGKTVLASDTHQR-GYNFEDLTDR-a11-b5 | pr\|PROT02\|WDR26 | pr\|PROT02\|WDR26 | **intra-protein xl** | aleitner_D1 | 402 | 419 | 17 | 3683.707 | 921.935 | 4 | -0.7 | 0.24 | 0 | **32.57** |
| MSYAEKPDEITK-DEWMEK-a6-b1 | pr\|PROT03\|Twa1 | pr\|PROT03\|Twa1 | **intra-protein xl** | aleitner_D1 | 30 | 37 | 7 | 2228.991 | 744.005 | 3 | -2.6 | 0.26 | 0 | **32.42** |
| ESTPKLAK-DEWMEK-a5-b1 | pr\|PROT03\|Twa1 | pr\|PROT03\|Twa1 | **intra-protein xl** | aleitner_D1 | 215 | 37 | 178 | 1690.82 | 564.615 | 3 | -2.1 | 0.22 | 0 | **32.35** |
| CLYHNTKLDNNLDSVSLLIDHVCSR-SELPIAELTGHTR-a7-b7 | pr\|PROT02\|WDR26 | pr\|PROT02\|WDR26 | **intra-protein xl** | aleitner_D1 | 226 | 512 | 286 | 4390.177 | 879.043 | 5 | -0.6 | 0.23 | 0 | **32.26** |
| MTDLSKGVIEEPK-LIMNYLVTEGFK-a6-b9 | pr\|PROT03\|Twa1 | pr\|PROT03\|Twa1 | **intra-protein xl** | aleitner_D1 | 245 | 64 | 181 | 2854.483 | 714.628 | 4 | -1.2 | 0.21 | 0 | **32.04** |
| RKAVESIQAEDESAK-ETSHVTMVVAELEK-a2-b1 | pr\|PROT04\|MAEA | pr\|PROT04\|MAEA | **intra-protein xl** | aleitner_D1 | 107 | 66 | 41 | 3213.611 | 804.411 | 4 | -2.4 | 0.27 | 0 | **32** |
| AKAEWEGKGTASR-YLEYLEDGK-a8-b7 | pr\|PROT02\|WDR26 | pr\|PROT02\|WDR26 | **intra-protein xl** | aleitner_D1 | 178 | 130 | 48 | 2500.218 | 626.062 | 4 | -2.2 | 0.26 | 0 | **31.68** |
| YLEYLEDGK-AKAEWEGK-a7-b2 | pr\|PROT02\|WDR26 | pr\|PROT02\|WDR26 | **intra-protein xl** | aleitner_D1 | 130 | 172 | 42 | 2027.978 | 677 | 3 | -3 | 0.26 | 0 | **31.58** |
| AAQKNIDRETSHVTMVVAELEK-AVESIQAEDESAK-a4-b3 | pr\|PROT04\|MAEA | pr\|PROT04\|MAEA | **intra-protein xl** | aleitner_D1 | 61 | 110 | 49 | 3825.894 | 766.187 | 5 | -3 | 0.27 | 0 | **30.9** |
| NPNLLFTLKVR-ELQAMSEQLR-a9-b1 | pr\|PROT01\|RanBP9 | pr\|PROT01\|RanBP9 | **intra-protein xl** | aleitner_D1 | 469 | 650 | 181 | 2499.356 | 625.847 | 4 | -2.7 | 0.3 | 0 | **30.43** |
| CLYHNTKLDNNLDSVSLLIDHVCSR-RSELPIAELTGHTR-a7-b8 | pr\|PROT02\|WDR26 | pr\|PROT02\|WDR26 | **intra-protein xl** | aleitner_D1 | 226 | 512 | 286 | 4546.285 | 758.722 | 6 | 1 | 0.33 | 0 | **29.5** |
| ETSHVTMVVAELEK-KAVESIQAEDESAK-a1-b1 | pr\|PROT04\|MAEA | pr\|PROT04\|MAEA | **intra-protein xl** | aleitner_D1 | 66 | 107 | 41 | 3057.516 | 765.387 | 4 | -0.7 | 0.36 | 0 | **29.29** |
| XSYAEKPDEITK-DEWXEK-a6-b1 | pr\|PROT03\|Twa1 | pr\|PROT03\|Twa1 | **intra-protein xl** | aleitner_D1 | 30 | 37 | 7 | 2260.978 | 754.667 | 3 | -3.9 | 0.18 | 0 | **28.83** |
| MSYAEKPDEITK-DEWXEK-a6-b1 | pr\|PROT03\|Twa1 | pr\|PROT03\|Twa1 | **intra-protein xl** | aleitner_D1 | 30 | 37 | 7 | 2244.984 | 749.336 | 3 | -3.5 | 0.21 | 0 | **28.46** |
| STDQTVLEELASIKNR-EGEWQTMIQK-a14-b3 | pr\|PROT01\|RanBP9 | pr\|PROT01\|RanBP9 | **intra-protein xl** | aleitner_D1 | 429 | 388 | 41 | 3033.508 | 759.385 | 4 | 0.2 | 0.23 | 0 | **27.94** |

### Low-res MS2

**Cross-linked peptide pairs in the 5mer complex identified using DMTMM as the cross-linking reagent and low-resolution MS/MS detection**

| Id | Protein1 | Protein2 | **XLType** | Spectrum | AbsPos1 | AbsPos2 | deltaAA | Mr | Mz | z | Error_rel[p | TIC | deltaS | **Id-Score** |
|---|---|---|---|---|---|---|---|---|---|---|---|---|---|---|
| IQAQIDR-LAKLLK-a6-b3 | pr\|PROT01\|RanBP9 | pr\|PROT03\|Twa1 | **inter-protein xl** | aleitner_D1 | 378 | 218 | | 1508.936 | 378.242 | 4 | -3.1 | 0.5 | 0.88 | **28.1** |
| HFSQAEGSQLDEVR-TPKDAASVR-a12-b3 | pr\|PROT04\|MAEA | pr\|PROT01\|RanBP9 | **inter-protein xl** | aleitner_D1 | 258 | 221 | | 2527.235 | 632.817 | 4 | -2.5 | 0.31 | 0.34 | **27.11** |
| IQAQIDRFPIGDR-LAKLLK-a6-b3 | pr\|PROT01\|RanBP9 | pr\|PROT03\|Twa1 | **inter-protein xl** | aleitner_D1 | 378 | 218 | | 2194.288 | 439.865 | 5 | -3.1 | 0.29 | 0.62 | **24.28** |
| QKVWSEVNQAVLDYENR-EGEWQTMIQK-a2-b1 | pr\|PROT03\|Twa1 | pr\|PROT01\|RanBP9 | **inter-protein xl** | aleitner_D1 | 195 | 386 | | 3307.586 | 1103.537 | 3 | -2 | 0.31 | 0.44 | **23.77** |
| QKVWSEVNQAVLDYENR-EGEWQTXIQK-a2-b3 | pr\|PROT03\|Twa1 | pr\|PROT01\|RanBP9 | **inter-protein xl** | aleitner_D1 | 195 | 388 | | 3323.577 | 831.902 | 4 | -3.2 | 0.34 | 0 | **22.05** |
| AEWEGK-SKLLDK-a4-b2 | pr\|PROT02\|WDR26 | pr\|PROT02\|WDR26 | **intra-protein xl** | aleitner_D1 | 176 | 185 | 9 | 1402.741 | 468.588 | 3 | -3.1 | 0.39 | 0 | **31.34** |
| GVIEEPK-LAKLLK-a5-b3 | pr\|PROT03\|Twa1 | pr\|PROT03\|Twa1 | **intra-protein xl** | aleitner_D1 | 250 | 218 | 32 | 1436.893 | 360.231 | 4 | -2.3 | 0.57 | 0 | **30.63** |
| DEWMEK-LAKLLK-a2-b3 | pr\|PROT03\|Twa1 | pr\|PROT03\|Twa1 | **intra-protein xl** | aleitner_D1 | 38 | 218 | 180 | 1502.812 | 501.945 | 3 | -2.8 | 0.23 | 0 | **29.4** |
| YLEYLEDGK-AKAEWEGK-a6-b2 | pr\|PROT02\|WDR26 | pr\|PROT02\|WDR26 | **intra-protein xl** | aleitner_D1 | 129 | 172 | 43 | 2027.975 | 508.002 | 4 | -4.5 | 0.2 | 0 | **27.13** |
| DGYMGIGLSAQGVNMNR-TPKDAASVR-a1-b3 | pr\|PROT01\|RanBP9 | pr\|PROT01\|RanBP9 | **intra-protein xl** | aleitner_D1 | 251 | 221 | 30 | 2707.309 | 903.444 | 3 | -2.9 | 0.41 | 0.36 | **26.85** |
| MSYAEKPDEITKDEWMEK-XSYAEKPDEITKDEWMEK-a5-b6 | pr\|PROT03\|Twa1 | pr\|PROT03\|Twa1 | **intra-protein xl** | aleitner_D1 | 29 | 30 | 1 | 4455.962 | 892.2 | 5 | -3.6 | 0.24 | 0 | **26.64** |
| XSYAEKPDEITK-DEWMEK-a6-b1 | pr\|PROT03\|Twa1 | pr\|PROT03\|Twa1 | **intra-protein xl** | aleitner_D1 | 30 | 37 | 7 | 2244.987 | 749.337 | 3 | -2.1 | 0.17 | 0 | **24.92** |
| DEWMEK-LAKLLK-a1-b3 | pr\|PROT03\|Twa1 | pr\|PROT03\|Twa1 | **intra-protein xl** | aleitner_D1 | 37 | 218 | 181 | 1502.813 | 501.946 | 3 | -2.4 | 0.19 | 0 | **24.46** |
| GVIEEPK-KVKYPK-a5-b3 | pr\|PROT03\|Twa1 | pr\|PROT03\|Twa1 | **intra-protein xl** | aleitner_D1 | 250 | 236 | 14 | 1513.882 | 505.635 | 3 | -3.2 | 0.33 | 0 | **24.39** |
| RKAVESIQAEDESAK-EVEESLER-a2-b1 | pr\|PROT04\|MAEA | pr\|PROT04\|MAEA | **intra-protein xl** | aleitner_D1 | 107 | 188 | 81 | 2631.29 | 658.83 | 4 | -3.2 | 0.37 | 0.22 | **23.51** |
| VKYPKMTDLSK-GVIEEPK-a5-b4 | pr\|PROT03\|Twa1 | pr\|PROT03\|Twa1 | **intra-protein xl** | aleitner_D1 | 239 | 249 | 10 | 2061.11 | 516.285 | 4 | -4.1 | 0.21 | 0.75 | **23.08** |
| AEWEGKGTASR-YLEYLEDGK-a6-b7 | pr\|PROT02\|WDR26 | pr\|PROT02\|WDR26 | **intra-protein xl** | aleitner_D1 | 178 | 130 | 48 | 2301.086 | 768.036 | 3 | -2.5 | 0.18 | 0 | **22.85** |
| AKAEWEGKGTASR-YLEYLEDGK-a8-b7 | pr\|PROT02\|WDR26 | pr\|PROT02\|WDR26 | **intra-protein xl** | aleitner_D1 | 178 | 130 | 48 | 2500.213 | 501.05 | 5 | -4.2 | 0.19 | 0.29 | **20.14** |
| MSYAEKPDEITK-DEWMEK-a6-b1 | pr\|PROT03\|Twa1 | pr\|PROT03\|Twa1 | **intra-protein xl** | aleitner_D1 | 30 | 37 | 7 | 2228.992 | 744.005 | 3 | -2 | 0.15 | 0 | **19.72** |
| AVESIQAEDESAK-LSVLKR-a3-b5 | pr\|PROT04\|MAEA | pr\|PROT04\|MAEA | **intra-protein xl** | aleitner_D1 | 110 | 105 | 5 | 2072.106 | 519.034 | 4 | -2.7 | 0.21 | 0.75 | **19.08** |
| MTDLSKGVIEEPK-LIMNYLVTEGFK-a6-b9 | pr\|PROT03\|Twa1 | pr\|PROT03\|Twa1 | **intra-protein xl** | aleitner_D1 | 245 | 64 | 181 | 2854.479 | 714.628 | 4 | -2.4 | 0.34 | 0.48 | **18.99** |
| VQCLWCLSDGKTVLASDTHQR-GYNFEDLTDR-a11-b5 | pr\|PROT02\|WDR26 | pr\|PROT02\|WDR26 | **intra-protein xl** | aleitner_D1 | 402 | 419 | 17 | 3683.703 | 737.748 | 5 | -1.9 | 0.21 | 0.48 | **18.95** |
| HFSQAEGSQLDEVR-TKEVFHFSQAEK-a12-b2 | pr\|PROT04\|MAEA | pr\|PROT04\|MAEA | **intra-protein xl** | aleitner_D1 | 258 | 405 | 147 | 3033.452 | 607.698 | 5 | -2 | 0.23 | 0.65 | **18.51** |
| LLNEVMVEHFFR-DALNKMFNGSK-a4-b5 | pr\|PROT05\|RMND5A | pr\|PROT05\|RMND5A | **intra-protein xl** | aleitner_D1 | 144 | 389 | 245 | 2738.365 | 685.599 | 4 | -1 | 0.2 | 0.57 | **18.29** |
| EHSSDQPAAASVWKR-VQEYPTLK-a14-b3 | pr\|PROT04\|MAEA | pr\|PROT04\|MAEA | **intra-protein xl** | aleitner_D1 | 144 | 41 | 103 | 2626.309 | 657.585 | 4 | -1.9 | 0.33 | 0.47 | **18.18** |

**Description of the column headers**

| | |
|---|---|
| *Id* | Assigned peptides and cross-linking sites within the *peptide* sequences. The longer peptide is designated as (a)lpha, the shorter as (b)eta. |
| *Protein1* | SwissProt/UniProt accession number and identifier of the protein 1 (containing peptide desginated as alpha). |
| *Protein2* | SwissProt/UniProt accession number and identifier of the protein 2 (containing peptide desginated as beta). |
| *XLType* | Intra- or inter-protein, sometimes also ambiguous. |
| *Spectrum* | References the actual MS/MS spectra that were used for the assignment. |
| *AbsPos1* | Position in the *protein* sequence of protein 1. |
| *AbsPos2* | Position in the *protein* sequence of protein 2. |
| *deltaAA* | Distance in the primary sequence (makes only sense for intra-proteins cross-links). |
| *Mr* | Molecular mass calculated from experimental m/z and z (neutral mass). |
| *Mz* | Experimentally observed mass-to-charge ratio of the precursor ion in Da. |
| *z* | Experimentally observed precursor charge. |
| *Error [ppm]* | Deviation between experimental and theoretical mass in ppm. |
| *TIC* | Fraction of the total ion current covered by assigned fragment ions. |
| *delta score* | Relative score for the top hit in relation to the second best hit for this spectrum, calculated as Id-score of 2nd best hit divided by Id-score of best hit. "0" means no second best hit exists. |
| *Id-score* | Identification score as assigned by xQuest. The higher, the better. |
| *X* | in the amino acid sequence denotes an oxidized Met residue |
