## Supplementary material for "The human GID complex engages two independent modules for substrate recruitment": Supp. Table 2

**Supplementary table 2**

### High-res MS2

**Cross-linked peptide pairs in the 5mer complex identified using DSS as the cross-linking reagent and high-resolution MS/MS detection**

| Id | Protein1 | Protein2 | **XLType** | Spectrum | AbsPos1 | AbsPos2 | deltaAA | Mr | Mz | z | Error_rel[p | TIC | deltaS | **Id-Score** |
|---|---|---|---|---|---|---|---|---|---|---|---|---|---|---|
| IQKLVLAGR-EAAEKFR-a3-b5 | pr\|PROT01\|RanBP9 | pr\|PROT03\|Twa1 | **inter-protein xl** | aleitner_D1 | 436 | 72 | | 1984.142 | 662.389 | 3 | -2.4 | 0.3 | 0 | **38.58** |
| DCGKNTANK-AKAEWEGK-a4-b2 | pr\|PROT01\|RanBP9 | pr\|PROT02\|WDR26 | **inter-protein xl** | aleitner_D1 | 664 | 172 | | 2061.974 | 688.332 | 3 | -2.5 | 0.37 | 0 | **38.11** |
| CGYYNTAVKLAR-ILEALKVR-a9-b6 | pr\|PROT04\|MAEA | pr\|PROT05\|RMND5A | **inter-protein xl** | aleitner_D1 | 167 | 189 | | 2493.373 | 624.351 | 4 | -1.9 | 0.24 | 0 | **36.91** |
| KHFSQAEGSQLDEVR-LASDHKDIHSSVSR-a1-b6 | pr\|PROT04\|MAEA | pr\|PROT05\|RMND5A | **inter-protein xl** | aleitner_D1 | 246 | 104 | | 3418.679 | 855.678 | 4 | -2.2 | 0.33 | 0 | **36.71** |
| LLLWAQNELDQKK-TKIQAQIDR-a12-b2 | pr\|PROT03\|Twa1 | pr\|PROT01\|RanBP9 | **inter-protein xl** | aleitner_D1 | 233 | 372 | | 2807.551 | 702.896 | 4 | -1.2 | 0.29 | 0 | **36.18** |
| EHSSDQPAAASVWKR-ILEALKVR-a14-b6 | pr\|PROT04\|MAEA | pr\|PROT05\|RMND5A | **inter-protein xl** | aleitner_D1 | 144 | 189 | | 2746.469 | 687.625 | 4 | -2.8 | 0.2 | 0 | **35.95** |
| TKIQAQIDRFPIGDR-LLLWAQNELDQKK-a2-b12 | pr\|PROT01\|RanBP9 | pr\|PROT03\|Twa1 | **inter-protein xl** | aleitner_D1 | 372 | 233 | | 3492.894 | 1165.306 | 3 | -4.3 | 0.24 | 0 | **34.64** |
| LATGSKDTTVIIWQVDPDTHLLK-TPKDAASVR-a6-b3 | pr\|PROT02\|WDR26 | pr\|PROT01\|RanBP9 | **inter-protein xl** | aleitner_D1 | 279 | 221 | | 3631.941 | 908.993 | 4 | -1.4 | 0.36 | 0 | **34.08** |
| AAQKNIDRETSHVTMVVAELEK-LASDHKDIHSSVSR-a4-b6 | pr\|PROT04\|MAEA | pr\|PROT05\|RMND5A | **inter-protein xl** | aleitner_D1 | 61 | 104 | | 4157.117 | 832.431 | 5 | -0.1 | 0.22 | 0 | **32.36** |
| EHSSDQPAAASVWKR-KCWYHSIFACPILR-a14-b1 | pr\|PROT04\|MAEA | pr\|PROT05\|RMND5A | **inter-protein xl** | aleitner_D1 | 144 | 351 | | 3655.783 | 914.954 | 4 | 0.3 | 0.17 | 0 | **31.68** |
| EGEWQTMIQKMVSSYLVHHGYCATAEAFAR-XTDLSKGVIEEPK-a10-b6 | pr\|PROT01\|RanBP9 | pr\|PROT03\|Twa1 | **inter-protein xl** | aleitner_D1 | 395 | 245 | | 5099.422 | 1020.892 | 5 | 0.8 | 0.25 | 0 | **28.00** |
| AEWEGKGTASR-SKLLDK-a6-b2 | pr\|PROT02\|WDR26 | pr\|PROT02\|WDR26 | **intra-protein xl** | aleitner_D1 | 178 | 185 | 7 | 2031.058 | 678.027 | 3 | -2.8 | 0.46 | 0.9 | **38.72** |
| TPKDAASVR-SWSPKDK-a3-b5 | pr\|PROT01\|RanBP9 | pr\|PROT01\|RanBP9 | **intra-protein xl** | aleitner_D1 | 221 | 196 | 25 | 1927.996 | 643.673 | 3 | -2.3 | 0.39 | 0 | **38.47** |
| KHFSQAEGSQLDEVR-AAQKNIDR-a1-b4 | pr\|PROT04\|MAEA | pr\|PROT04\|MAEA | **intra-protein xl** | aleitner_D1 | 246 | 61 | 185 | 2782.396 | 696.607 | 4 | -1.6 | 0.34 | 0 | **38.02** |
| KAVESIQAEDESAK-AAQKNIDR-a1-b4 | pr\|PROT04\|MAEA | pr\|PROT04\|MAEA | **intra-protein xl** | aleitner_D1 | 107 | 61 | 46 | 2556.301 | 853.108 | 3 | -0.8 | 0.47 | 0 | **38.02** |
| LLDKLQTYLPPSVMLPPR-CELTPLKYNTER-a4-b7 | pr\|PROT02\|WDR26 | pr\|PROT02\|WDR26 | **intra-protein xl** | aleitner_D1 | 189 | 148 | 41 | 3740.98 | 936.253 | 4 | -2.2 | 0.3 | 0 | **37.87** |
| TKEVFHFSQAEK-QDDKVVCPR-a2-b4 | pr\|PROT04\|MAEA | pr\|PROT04\|MAEA | **intra-protein xl** | aleitner_D1 | 405 | 398 | 7 | 2703.326 | 902.117 | 3 | -2.3 | 0.25 | 0.81 | **37.82** |
| TPKDAASVR-IVSKGR-a3-b4 | pr\|PROT01\|RanBP9 | pr\|PROT01\|RanBP9 | **intra-protein xl** | aleitner_D1 | 221 | 248 | 27 | 1739.985 | 436.004 | 4 | -2.3 | 0.41 | 0 | **37.71** |
| DEWMEKLNNLHVQR-MSYAEKPDEITK-a6-b6 | pr\|PROT03\|Twa1 | pr\|PROT03\|Twa1 | **intra-protein xl** | aleitner_D1 | 42 | 30 | 12 | 3359.606 | 672.929 | 5 | -3.1 | 0.32 | 0.66 | **37.49** |
| CELTPLKYNTER-AKAEWEGK-a7-b2 | pr\|PROT02\|WDR26 | pr\|PROT02\|WDR26 | **intra-protein xl** | aleitner_D1 | 148 | 172 | 24 | 2578.27 | 645.575 | 4 | -1.5 | 0.37 | 0 | **37.16** |
| RETATCLAWCHDNKSR-AAQKNIDR-a14-b4 | pr\|PROT04\|MAEA | pr\|PROT04\|MAEA | **intra-protein xl** | aleitner_D1 | 209 | 61 | 148 | 3056.468 | 612.301 | 5 | 0.3 | 0.19 | 0 | **37.04** |
| RKAVESIQAEDESAK-AAQKNIDR-a2-b4 | pr\|PROT04\|MAEA | pr\|PROT04\|MAEA | **intra-protein xl** | aleitner_D1 | 107 | 61 | 46 | 2712.398 | 679.107 | 4 | -2.2 | 0.35 | 0 | **37.02** |
| VPYETLNKR-AAQKNIDR-a8-b4 | pr\|PROT04\|MAEA | pr\|PROT04\|MAEA | **intra-protein xl** | aleitner_D1 | 54 | 61 | 7 | 2171.165 | 543.799 | 4 | -2.4 | 0.3 | 0 | **37.01** |
| AKAEWEGK-SKLLDK-a2-b2 | pr\|PROT02\|WDR26 | pr\|PROT02\|WDR26 | **intra-protein xl** | aleitner_D1 | 172 | 185 | 13 | 1757.954 | 586.992 | 3 | -1.4 | 0.39 | 0 | **36.94** |
| SLNKLAQPLPMAHCANSR-EDGSSKSPDCPVCSR-a4-b6 | pr\|PROT04\|MAEA | pr\|PROT04\|MAEA | **intra-protein xl** | aleitner_D1 | 346 | 333 | 13 | 3824.761 | 957.198 | 4 | -2.5 | 0.38 | 0.9 | **36.81** |
| MSYAEKPDEITKDEWMEK-MTDLSKGVIEEPK-a12-b6 | pr\|PROT03\|Twa1 | pr\|PROT03\|Twa1 | **intra-protein xl** | aleitner_D1 | 36 | 245 | 209 | 3812.8 | 763.568 | 5 | -2.3 | 0.24 | 0 | **36.66** |
| KAVESIQAEDESAK-VPYETLNKR-a1-b8 | pr\|PROT04\|MAEA | pr\|PROT04\|MAEA | **intra-protein xl** | aleitner_D1 | 107 | 54 | 53 | 2760.411 | 921.145 | 3 | -2.7 | 0.27 | 0 | **36.62** |
| YPKMTDLSK-LAKLLK-a3-b3 | pr\|PROT03\|Twa1 | pr\|PROT03\|Twa1 | **intra-protein xl** | aleitner_D1 | 239 | 218 | 21 | 1904.101 | 635.708 | 3 | -2.2 | 0.41 | 0 | **36.58** |
| RETATCLAWCHDNKSR-KHFSQAEGSQLDEVR-a14-b1 | pr\|PROT04\|MAEA | pr\|PROT04\|MAEA | **intra-protein xl** | aleitner_D1 | 209 | 246 | 37 | 3871.804 | 775.369 | 5 | -2 | 0.32 | 0 | **36.47** |
| TPQCYKEDGSSKSPDCPVCSR-SLNKLAQPLPMAHCANSR-a12-b4 | pr\|PROT04\|MAEA | pr\|PROT04\|MAEA | **intra-protein xl** | aleitner_D1 | 333 | 346 | 13 | 4602.109 | 921.43 | 5 | -2.1 | 0.26 | 0.67 | **36.24** |
| CELTPLKYNTER-MKFLLLQQK-a7-b2 | pr\|PROT02\|WDR26 | pr\|PROT02\|WDR26 | **intra-protein xl** | aleitner_D1 | 148 | 116 | 32 | 2808.487 | 937.17 | 3 | -1.8 | 0.36 | 0 | **36.03** |
| ETATCLAWCHDNKSR-KAVESIQAEDESAK-a13-b1 | pr\|PROT04\|MAEA | pr\|PROT04\|MAEA | **intra-protein xl** | aleitner_D1 | 209 | 107 | 102 | 3489.604 | 1164.209 | 3 | -2.9 | 0.28 | 0 | **35.95** |
| TKMSQSHEDSLTSVAWNPDGK-FSNDGTKLATGSK-a2-b7 | pr\|PROT02\|WDR26 | pr\|PROT02\|WDR26 | **intra-protein xl** | aleitner_D1 | 344 | 273 | 71 | 3779.794 | 945.956 | 4 | -0.3 | 0.3 | 0 | **35.76** |
| LASDHKDIHSSVSR-VGKAIDK-a6-b3 | pr\|PROT05\|RMND5A | pr\|PROT05\|RMND5A | **intra-protein xl** | aleitner_D1 | 104 | 115 | 11 | 2418.28 | 484.664 | 5 | -3 | 0.32 | 0.66 | **35.59** |
| ETATCLAWCHDNKSR-KHFSQAEGSQLDEVR-a13-b1 | pr\|PROT04\|MAEA | pr\|PROT04\|MAEA | **intra-protein xl** | aleitner_D1 | 209 | 246 | 37 | 3715.705 | 929.934 | 4 | -1.5 | 0.35 | 0 | **35.42** |
| STDQTVLEELASIKNR-IQKLVLAGR-a14-b3 | pr\|PROT01\|RanBP9 | pr\|PROT01\|RanBP9 | **intra-protein xl** | aleitner_D1 | 429 | 436 | 7 | 2937.644 | 980.223 | 3 | -1.9 | 0.22 | 0 | **35.42** |
| ELEKVLHK-VGKAIDK-a4-b3 | pr\|PROT05\|RMND5A | pr\|PROT05\|RMND5A | **intra-protein xl** | aleitner_D1 | 37 | 115 | 78 | 1862.083 | 466.529 | 4 | -2.6 | 0.21 | 0 | **35.34** |
| MSYAEKPDEITKDEWMEK-LAKLLK-a12-b3 | pr\|PROT03\|Twa1 | pr\|PROT03\|Twa1 | **intra-protein xl** | aleitner_D1 | 36 | 218 | 182 | 3051.55 | 611.318 | 5 | -1.7 | 0.23 | 0 | **34.60** |
| SLNKLAQPLPMAHCANSR-EHSSDQPAAASVWKR-a4-b14 | pr\|PROT04\|MAEA | pr\|PROT04\|MAEA | **intra-protein xl** | aleitner_D1 | 346 | 144 | 202 | 3812.872 | 763.582 | 5 | -3.1 | 0.24 | 0 | **34.09** |
| MTDLSKGVIEEPK-ESTPKLAK-a6-b5 | pr\|PROT03\|Twa1 | pr\|PROT03\|Twa1 | **intra-protein xl** | aleitner_D1 | 245 | 215 | 30 | 2456.299 | 615.083 | 4 | -3.9 | 0.2 | 0 | **34.06** |
| DEWXEKLNNLHVQR-XSYAEKPDEITK-a6-b6 | pr\|PROT03\|Twa1 | pr\|PROT03\|Twa1 | **intra-protein xl** | aleitner_D1 | 42 | 30 | 12 | 3391.605 | 848.909 | 4 | -0.5 | 0.15 | 0 | **34.03** |
| MSYAEKPDEITKDEWMEK-LLLWAQNELDQKK-a6-b12 | pr\|PROT03\|Twa1 | pr\|PROT03\|Twa1 | **intra-protein xl** | aleitner_D1 | 30 | 233 | 203 | 3964.941 | 793.996 | 5 | -1.7 | 0.16 | 0 | **33.68** |
| VQCLWCLSDGKTVLASDTHQR-MSQSHEDSLTSVAWNPDGKR-a11-b19 | pr\|PROT02\|WDR26 | pr\|PROT02\|WDR26 | **intra-protein xl** | aleitner_D1 | 402 | 363 | 39 | 4855.266 | 972.061 | 5 | -1.8 | 0.15 | 0 | **33.63** |
| ETATCLAWCHDNKSR-VPYETLNKR-a13-b8 | pr\|PROT04\|MAEA | pr\|PROT04\|MAEA | **intra-protein xl** | aleitner_D1 | 209 | 54 | 155 | 3104.474 | 777.126 | 4 | -2.1 | 0.12 | 0 | **33.25** |
| KHFSQAEGSQLDEVR-KAVESIQAEDESAK-a1-b1 | pr\|PROT04\|MAEA | pr\|PROT04\|MAEA | **intra-protein xl** | aleitner_D1 | 246 | 107 | 139 | 3371.642 | 843.918 | 4 | -1.7 | 0.33 | 0 | **31.39** |
| MSYAEKPDEITKDEWMEK-ESTPKLAK-a12-b5 | pr\|PROT03\|Twa1 | pr\|PROT03\|Twa1 | **intra-protein xl** | aleitner_D1 | 36 | 215 | 179 | 3239.551 | 648.918 | 5 | -3.2 | 0.12 | 0 | **31.37** |
| MSYAEKPDEITKDEWMEK-XTDLSKGVIEEPK-a12-b6 | pr\|PROT03\|Twa1 | pr\|PROT03\|Twa1 | **intra-protein xl** | aleitner_D1 | 36 | 245 | 209 | 3828.799 | 958.208 | 4 | -1.1 | 0.19 | 0 | **31.19** |
| VQCLWCLSDGKTVLASDTHQR-XSQSHEDSLTSVAWNPDGKR-a11-b19 | pr\|PROT02\|WDR26 | pr\|PROT02\|WDR26 | **intra-protein xl** | aleitner_D1 | 402 | 363 | 39 | 4871.251 | 975.258 | 5 | -4 | 0.18 | 0 | **31.09** |
| QKVWSEVNQAVLDYENR-MTDLSKGVIEEPK-a2-b6 | pr\|PROT03\|Twa1 | pr\|PROT03\|Twa1 | **intra-protein xl** | aleitner_D1 | 195 | 245 | 50 | 3660.832 | 916.216 | 4 | -0.6 | 0.37 | 0 | **30.46** |
| SLNKLAQPLPXAHCANSR-EDGSSKSPDCPVCSR-a4-b6 | pr\|PROT04\|MAEA | pr\|PROT04\|MAEA | **intra-protein xl** | aleitner_D1 | 346 | 333 | 13 | 3840.76 | 769.16 | 5 | -1.4 | 0.34 | 0.77 | **30.24** |

### Low-res MS2

**Cross-linked peptide pairs in the 5mer complex identified using DSS as the cross-linking reagent and low-resolution MS/MS detection**

| Id | Protein1 | Protein2 | XLType | Spectrum | AbsPos1 | AbsPos2 | deltaAA | Mr | Mz | z | Error_rel[p | TIC | deltaS | **Id-Score** |
|---|---|---|---|---|---|---|---|---|---|---|---|---|---|---|
| IQKLVLAGR-EAAEKFR-a3-b5 | pr\|PROT01\|RanBP9 | pr\|PROT03\|Twa1 | inter-protein xl | aleitner_D1 | 436 | 72 | | 1984.143 | 497.044 | 4 | -1.8 | 0.3 | 0.69 | **28.74** |
| AAQKNIDR-VGKAIDK-a4-b3 | pr\|PROT04\|MAEA | pr\|PROT05\|RMND5A | inter-protein xl | aleitner_D1 | 61 | 115 | | 1781.996 | 595.006 | 3 | -2.4 | 0.25 | 0 | **28.62** |
| DCGKNTANK-AKAEWEGK-a4-b2 | pr\|PROT01\|RanBP9 | pr\|PROT02\|WDR26 | inter-protein xl | aleitner_D1 | 664 | 172 | | 2061.974 | 688.332 | 3 | -2.5 | 0.22 | 0.66 | **28.41** |
| LLLWAQNELDQKK-TKIQAQIDR-a12-b2 | pr\|PROT03\|Twa1 | pr\|PROT01\|RanBP9 | inter-protein xl | aleitner_D1 | 233 | 372 | | 2807.552 | 702.896 | 4 | -1 | 0.2 | 0 | **26.99** |
| TKMSQSHEDSLTSVAWNPDGK-TPKDAASVR-a2-b3 | pr\|PROT02\|WDR26 | pr\|PROT01\|RanBP9 | inter-protein xl | aleitner_D1 | 344 | 221 | | 3398.635 | 850.667 | 4 | -1.8 | 0.24 | 0.36 | **25.37** |
| CGYYNTAVKLAR-ILEALKVR-a9-b6 | pr\|PROT04\|MAEA | pr\|PROT05\|RMND5A | inter-protein xl | aleitner_D1 | 167 | 189 | | 2493.372 | 624.351 | 4 | -2.3 | 0.32 | 0 | **24.01** |
| RETATCLAWCHDNKSR-AAQKNIDR-a14-b4 | pr\|PROT04\|MAEA | pr\|PROT04\|MAEA | intra-protein xl | aleitner_D1 | 209 | 61 | 148 | 3056.458 | 612.299 | 5 | -2.8 | 0.24 | 0 | **31.35** |
| LASDHKDIHSSVSR-VGKAIDK-a6-b3 | pr\|PROT05\|RMND5A | pr\|PROT05\|RMND5A | intra-protein xl | aleitner_D1 | 104 | 115 | 11 | 2418.283 | 605.578 | 4 | -1.7 | 0.42 | 0.68 | **31.17** |
| STDQTVLEELASIKNR-IQKLVLAGR-a14-b3 | pr\|PROT01\|RanBP9 | pr\|PROT01\|RanBP9 | intra-protein xl | aleitner_D1 | 429 | 436 | 7 | 2937.645 | 735.419 | 4 | -1.6 | 0.32 | 0.52 | **29.05** |
| KHFSQAEGSQLDEVR-AAQKNIDR-a1-b4 | pr\|PROT04\|MAEA | pr\|PROT04\|MAEA | intra-protein xl | aleitner_D1 | 246 | 61 | 185 | 2782.394 | 557.487 | 5 | -2.1 | 0.34 | 0 | **28.31** |
| MSYAEKPDEITKDEWMEK-LAKLLK-a12-b3 | pr\|PROT03\|Twa1 | pr\|PROT03\|Twa1 | intra-protein xl | aleitner_D1 | 36 | 218 | 182 | 3051.55 | 611.318 | 5 | -1.5 | 0.32 | 0.58 | **27.81** |
| RKAVESIQAEDESAK-AAQKNIDR-a2-b4 | pr\|PROT04\|MAEA | pr\|PROT04\|MAEA | intra-protein xl | aleitner_D1 | 107 | 61 | 46 | 2712.398 | 543.487 | 5 | -2.3 | 0.35 | 0 | **27.21** |
| DDDDKSAVDENLYFQGGGR-AAQKNIDR-a5-b4 | pr\|PROT04\|MAEA | pr\|PROT04\|MAEA | intra-protein xl | aleitner_D1 | 9 | 61 | 52 | 3152.46 | 631.5 | 5 | -1.5 | 0.59 | 0.43 | **27.15** |
| KAVESIQAEDESAK-VPYETLNKR-a1-b8 | pr\|PROT04\|MAEA | pr\|PROT04\|MAEA | intra-protein xl | aleitner_D1 | 107 | 54 | 53 | 2760.413 | 921.145 | 3 | -2 | 0.29 | 0 | **26.69** |
| KAVESIQAEDESAK-AAQKNIDR-a1-b4 | pr\|PROT04\|MAEA | pr\|PROT04\|MAEA | intra-protein xl | aleitner_D1 | 107 | 61 | 46 | 2556.299 | 853.107 | 3 | -1.6 | 0.37 | 0 | **25.86** |
| LEHPSATKFR-MKFLLLQQK-a8-b2 | pr\|PROT02\|WDR26 | pr\|PROT02\|WDR26 | intra-protein xl | aleitner_D1 | 66 | 116 | 50 | 2470.367 | 495.081 | 5 | -3.8 | 0.31 | 0 | **25.48** |
| VPYETLNKR-AAQKNIDR-a8-b4 | pr\|PROT04\|MAEA | pr\|PROT04\|MAEA | intra-protein xl | aleitner_D1 | 54 | 61 | 7 | 2171.165 | 724.73 | 3 | -2.2 | 0.18 | 0 | **25.08** |
| TPKDAASVR-IVSKGR-a3-b4 | pr\|PROT01\|RanBP9 | pr\|PROT01\|RanBP9 | intra-protein xl | aleitner_D1 | 221 | 248 | 27 | 1739.985 | 436.004 | 4 | -2.5 | 0.17 | 0 | **25.02** |
| ETATCLAWCHDNKSR-AAQKNIDR-a13-b4 | pr\|PROT04\|MAEA | pr\|PROT04\|MAEA | intra-protein xl | aleitner_D1 | 209 | 61 | 148 | 2900.358 | 726.097 | 4 | -2.6 | 0.14 | 0 | **24.95** |
| CELTPLKYNTER-AKAEWEGK-a7-b2 | pr\|PROT02\|WDR26 | pr\|PROT02\|WDR26 | intra-protein xl | aleitner_D1 | 148 | 172 | 24 | 2578.266 | 645.574 | 4 | -3.2 | 0.23 | 0 | **24.93** |
| VLHKFSGYGQLCER-DTVQKLASDHK-a4-b5 | pr\|PROT05\|RMND5A | pr\|PROT05\|RMND5A | intra-protein xl | aleitner_D1 | 41 | 98 | 57 | 3071.544 | 512.932 | 6 | -2 | 0.21 | 0 | **24.17** |
| TKEVFHFSQAEK-QDDKVVCPR-a2-b4 | pr\|PROT04\|MAEA | pr\|PROT04\|MAEA | intra-protein xl | aleitner_D1 | 405 | 398 | 7 | 2703.328 | 676.84 | 4 | -1.9 | 0.19 | 0.61 | **24.13** |
| RETATCLAWCHDNKSR-KHFSQAEGSQLDEVR-a14-b1 | pr\|PROT04\|MAEA | pr\|PROT04\|MAEA | intra-protein xl | aleitner_D1 | 209 | 246 | 37 | 3871.806 | 775.369 | 5 | -1.6 | 0.28 | 0 | **23.76** |
| LASDHKDIHSSVSR-IKDTVQK-a6-b2 | pr\|PROT05\|RMND5A | pr\|PROT05\|RMND5A | intra-protein xl | aleitner_D1 | 104 | 93 | 11 | 2519.33 | 504.874 | 5 | -1.9 | 0.21 | 0.75 | **23.53** |
| ELEKVLHK-VGKAIDK-a4-b3 | pr\|PROT05\|RMND5A | pr\|PROT05\|RMND5A | intra-protein xl | aleitner_D1 | 37 | 115 | 78 | 1862.082 | 466.528 | 4 | -3.2 | 0.19 | 0 | **23.5** |
| MTDLSKGVIEEPK-ESTPKLAK-a6-b5 | pr\|PROT03\|Twa1 | pr\|PROT03\|Twa1 | intra-protein xl | aleitner_D1 | 245 | 215 | 30 | 2456.303 | 615.084 | 4 | -2.2 | 0.19 | 0.76 | **23.41** |
| SLNKLAQPLPMAHCANSR-EDGSSKSPDCPVCSR-a4-b6 | pr\|PROT04\|MAEA | pr\|PROT04\|MAEA | intra-protein xl | aleitner_D1 | 346 | 333 | 13 | 3824.762 | 957.198 | 4 | -2.2 | 0.26 | 0.78 | **23.01** |
| ETATCLAWCHDNKSR-KHFSQAEGSQLDEVR-a13-b1 | pr\|PROT04\|MAEA | pr\|PROT04\|MAEA | intra-protein xl | aleitner_D1 | 209 | 246 | 37 | 3715.703 | 929.934 | 4 | -2.1 | 0.26 | 0 | **22.58** |
| MSYAEKPDEITKDEWMEK-MTDLSKGVIEEPK-a12-b6 | pr\|PROT03\|Twa1 | pr\|PROT03\|Twa1 | intra-protein xl | aleitner_D1 | 36 | 245 | 209 | 3812.806 | 954.209 | 4 | -0.8 | 0.17 | 0.7 | **22.33** |
| YLEYLEDGKVLEALQVLR-AEWEGKGTASR-a9-b6 | pr\|PROT02\|WDR26 | pr\|PROT02\|WDR26 | intra-protein xl | aleitner_D1 | 132 | 178 | 46 | 3478.793 | 870.706 | 4 | -1.6 | 0.23 | 0 | **22.21** |
| CELTPLKYNTER-MKFLLLQQK-a7-b2 | pr\|PROT02\|WDR26 | pr\|PROT02\|WDR26 | intra-protein xl | aleitner_D1 | 148 | 116 | 32 | 2808.482 | 937.168 | 3 | -3.5 | 0.11 | 0.7 | **21.95** |
| LLDKLQTYLPPSVMLPPR-CELTPLKYNTER-a4-b7 | pr\|PROT02\|WDR26 | pr\|PROT02\|WDR26 | intra-protein xl | aleitner_D1 | 189 | 148 | 41 | 3740.99 | 936.255 | 4 | 0.4 | 0.21 | 0 | **21.34** |
| LKCPYCPMEQSPGDAK-DALNKMFNGSK-a2-b5 | pr\|PROT05\|RMND5A | pr\|PROT05\|RMND5A | intra-protein xl | aleitner_D1 | 397 | 389 | 8 | 3241.483 | 811.379 | 4 | -2.5 | 0.22 | 0.79 | **21.2** |
| DKFSYIGLSQNNLR-VHYKGHGK-a2-b4 | pr\|PROT01\|RanBP9 | pr\|PROT01\|RanBP9 | intra-protein xl | aleitner_D1 | 198 | 214 | 16 | 2716.401 | 906.475 | 3 | -2.9 | 0.22 | 0.46 | **20.62** |
| TPQCYKEDGSSKSPDCPVCSR-SLNKLAQPLPMAHCANSR-a12-b4 | pr\|PROT04\|MAEA | pr\|PROT04\|MAEA | intra-protein xl | aleitner_D1 | 333 | 346 | 13 | 4602.106 | 921.429 | 5 | -2.6 | 0.17 | 0.84 | **19.83** |
| VLHKFSGYGQLCER-IKDTVQKLASDHK-a4-b7 | pr\|PROT05\|RMND5A | pr\|PROT05\|RMND5A | intra-protein xl | aleitner_D1 | 41 | 98 | 57 | 3312.725 | 663.553 | 5 | -1.1 | 0.16 | 0 | **19.64** |
| RKAVESIQAEDESAK-VPYETLNKR-a2-b8 | pr\|PROT04\|MAEA | pr\|PROT04\|MAEA | intra-protein xl | aleitner_D1 | 107 | 54 | 53 | 2916.514 | 730.136 | 4 | -1.8 | 0.17 | 0 | **19.38** |
| LLLWAQNELDQKK-MTDLSKGVIEEPK-a12-b6 | pr\|PROT03\|Twa1 | pr\|PROT03\|Twa1 | intra-protein xl | aleitner_D1 | 233 | 245 | 12 | 3181.689 | 796.43 | 4 | -1.6 | 0.14 | 0 | **19.15** |
| MSYAEKPDEITKDEWMEK-XTDLSKGVIEEPK-a6-b6 | pr\|PROT03\|Twa1 | pr\|PROT03\|Twa1 | intra-protein xl | aleitner_D1 | 30 | 245 | 215 | 3828.788 | 958.205 | 4 | -4 | 0.18 | 0.79 | **18.88** |
| VQCLWCLSDGKTVLASDTHQR-MSQSHEDSLTSVAWNPDGKR-a11-b19 | pr\|PROT02\|WDR26 | pr\|PROT02\|WDR26 | intra-protein xl | aleitner_D1 | 402 | 363 | 39 | 4855.27 | 1214.825 | 4 | -1 | 0.17 | 0 | **18.58** |
| XSYAEKPDEITKDEWMEK-MTDLSKGVIEEPK-a12-b6 | pr\|PROT03\|Twa1 | pr\|PROT03\|Twa1 | intra-protein xl | aleitner_D1 | 36 | 245 | 209 | 3828.792 | 766.766 | 5 | -3.1 | 0.12 | 0.89 | **17.8** |
| ETATCLAWCHDNKSR-KAVESIQAEDESAK-a13-b1 | pr\|PROT04\|MAEA | pr\|PROT04\|MAEA | intra-protein xl | aleitner_D1 | 209 | 107 | 102 | 3489.611 | 873.411 | 4 | -0.9 | 0.25 | 0 | **17.79** |
| MSYAEKPDEITKDEWMEK-MTDLSKGVIEEPK-a6-b6 | pr\|PROT03\|Twa1 | pr\|PROT03\|Twa1 | intra-protein xl | aleitner_D1 | 30 | 245 | 215 | 3812.798 | 763.567 | 5 | -2.8 | 0.14 | 0 | **17.66** |
| YPKMTDLSK-ESTPKLAK-a3-b5 | pr\|PROT03\|Twa1 | pr\|PROT03\|Twa1 | intra-protein xl | aleitner_D1 | 239 | 215 | 24 | 2092.107 | 524.035 | 4 | -2.5 | 0.14 | 0 | **16.31** |

**Description of the column headers**

| | |
|---|---|
| *Id* | Assigned peptides and cross-linking sites within the *peptide* sequences. The longer peptide is designated as (a)lpha, the shorter as (b)eta. |
| *Protein1* | SwissProt/UniProt accession number and identifier of the protein 1 (containing peptide desginated as alpha). |
| *Protein2* | SwissProt/UniProt accession number and identifier of the protein 2 (containing peptide desginated as beta). |
| *XLType* | Intra- or inter-protein, sometimes also ambiguous. |
| *Spectrum* | References the actual MS/MS spectra that were used for the assignment. |
| *AbsPos1* | Position in the *protein* sequence of protein 1. |
| *AbsPos2* | Position in the *protein* sequence of protein 2. |
| *deltaAA* | Distance in the primary sequence (makes only sense for intra-proteins cross-links). |
| *Mr* | Molecular mass calculated from experimental m/z and z (neutral mass). |
| *Mz* | Experimentally observed mass-to-charge ratio of the precursor ion in Da. |
| *z* | Experimentally observed precursor charge. |
| *Error [ppm]* | Deviation between experimental and theoretical mass in ppm. |
| *TIC* | Fraction of the total ion current covered by assigned fragment ions. |
| *delta score* | Relative score for the top hit in relation to the second best hit for this spectrum, calculated as Id-score of 2nd best hit divided by Id-score of best hit. "0" means no second best hit exists. |
| *Id-score* | Identification score as assigned by xQuest. The higher, the better. |
| *X* | in the amino acid sequence denotes an oxidized Met residue |
