## Supplementary material for "The human GID complex engages two independent modules for substrate recruitment": Supp. Table 3

**Supplementary table 3**

| | GID dataset 1 | GID dataset 2 | GID dataset 3 | GID-ARMC8β |
|---|---|---|---|---|
| **Movies** | 4759 | 26017 | 14840 | 3048 |
| **Detector** | K2 | K3 | K2 | K2 |
| **Mode** | counting | superresolution | counting | counting |
| **Magnification** | 165'000 | 105'000 | 165'000 | 130'000 |
| **Pixel size** | 0.84 | 0.42 | 0.84 | 1.07 |
| **Frames** | 40 | 29 | 40 | 40 |
| **Exposure time** | 8.5 | 1.56 | 8.5 | 8.5 |
| **Dose** | 78 | 78 | 78 | 80 |
| **Defocus** | -1.0 - -3.0 | -1.2 - -2.8 | -1.5 - -3.7 | -1.5 - -3.4 |
| **Particles No.** | 88564 | 538734 | 188240 | 73559 |
