## Supplementary material for "The human GID complex engages two independent modules for substrate recruitment": Supp. Table 4

**Supplementary table 4**

### High-res MS2

**Cross-linked peptide pairs in the 6mer complex identified using DMTMM as the cross-linking reagent and high-resolution MS/MS detection**

| Id | Protein1 | Protein2 | XLType | Spectrum | AbsPos1 | AbsPos2 | deltaAA | Mr | Mz | z | Error_rel[p | TIC | deltaS | Id-Score |
|---|---|---|---|---|---|---|---|---|---|---|---|---|---|---|
| IQAQIDRFPIGDR-LAKLLK-a6-b3 | pr\|PROT01\|RanBP9 | pr\|PROT03\|Twa1 | **inter-protein xl** | 210307_LU | 378 | 218 | | 2194.298 | 439.867 | 5 | 1.5 | 0.27 | 0 | **33.98** |
| IQAQIDR-LAKLLK-a6-b3 | pr\|PROT01\|RanBP9 | pr\|PROT03\|Twa1 | **inter-protein xl** | 210307_LU | 378 | 218 | | 1508.943 | 378.244 | 4 | 2.1 | 0.33 | 0 | **33.24** |
| QRETEAALEFAQTQLAEQGEESR-NAVIGNNKQK-a21-b8 | pr\|PROT03\|Twa1 | pr\|PROT06\|ARMC8b | **inter-protein xl** | 210307_LU | 159 | 62 | | 3686.834 | 738.375 | 5 | 2.4 | 0.27 | 0 | **33.22** |
| TKEVFHFSQAEK-LLEER-a2-b3 | pr\|PROT04\|MAEA | pr\|PROT06\|ARMC8b | **inter-protein xl** | 210307_LU | 405 | 305 | | 2090.083 | 419.024 | 5 | 1.7 | 0.22 | 0 | **32.8** |
| QRETEAALEFAQTQLAEQGEESR-NAVIGNNKQK-a17-b8 | pr\|PROT03\|Twa1 | pr\|PROT06\|ARMC8b | **inter-protein xl** | 210307_LU | 155 | 62 | | 3686.831 | 738.374 | 5 | 1.8 | 0.23 | 0 | **32.72** |
| LASDHKDIHSSVSR-AVESIQAEDESAK-a6-b3 | pr\|PROT05\|RMND5A | pr\|PROT04\|MAEA | **inter-protein xl** | 210307_LU | 104 | 110 | | 2908.42 | 728.113 | 4 | 1.4 | 0.26 | 0 | **32.19** |
| ELQAXSEQLR-AKAEWEGK-a7-b2 | pr\|PROT01\|RanBP9 | pr\|PROT02\|WDR26 | **inter-protein xl** | 210307_LU | 656 | 172 | | 2119.041 | 530.768 | 4 | 1.7 | 0.29 | 0 | **31.8** |
| MSYAEKPDEITKDEWXEK-RLYPAVDEQETPLPR-a6-b10 | pr\|PROT03\|Twa1 | pr\|PROT01\|RanBP9 | **inter-protein xl** | 210307_LU | 30 | 186 | | 4009.919 | 802.992 | 5 | 2.8 | 0.29 | 0 | **29.84** |
| KVSLGEGRPPVLTASR-AVESIQAEDESAK-a1-b9 | pr\|PROT06\|ARMC8b | pr\|PROT04\|MAEA | **inter-protein xl** | 210307_LU | 388 | 116 | | 3023.596 | 605.727 | 5 | 2.4 | 0.22 | 0 | **29.68** |
| ELQAMSEQLR-AKAEWEGK-a7-b2 | pr\|PROT01\|RanBP9 | pr\|PROT02\|WDR26 | **inter-protein xl** | 210307_LU | 656 | 172 | | 2103.045 | 702.023 | 3 | 1.4 | 0.29 | 0 | **29.66** |
| QKVWSEVNQAVLDYENR-EGEWQTMIQK-a2-b1 | pr\|PROT03\|Twa1 | pr\|PROT01\|RanBP9 | **inter-protein xl** | 210307_LU | 195 | 386 | | 3307.599 | 827.908 | 4 | 1.7 | 0.35 | 0 | **29.08** |
| KAVESIQAEDESAK-EVEESLER-a1-b1 | pr\|PROT04\|MAEA | pr\|PROT04\|MAEA | **intra-protein xl** | 210307_LU | 107 | 188 | 81 | 2475.204 | 826.076 | 3 | 2.7 | 0.27 | 0 | **35.39** |
| DELPIEVDLGKK-VGKAIDK-a2-b3 | pr\|PROT05\|RMND5A | pr\|PROT05\|RMND5A | **intra-protein xl** | 210307_LU | 341 | 115 | 226 | 2066.165 | 517.549 | 4 | 1.4 | 0.42 | 0 | **35.34** |
| LLNEVMVEHFFR-VGKAIDK-a4-b3 | pr\|PROT05\|RMND5A | pr\|PROT05\|RMND5A | **intra-protein xl** | 210307_LU | 144 | 115 | 29 | 2244.216 | 562.062 | 4 | 3.3 | 0.28 | 0 | **34.83** |
| GVIEEPK-LAKLLK-a4-b3 | pr\|PROT03\|Twa1 | pr\|PROT03\|Twa1 | **intra-protein xl** | 210307_LU | 249 | 218 | 31 | 1436.899 | 479.974 | 3 | 1.7 | 0.46 | 0 | **34.62** |
| LAKLLK-MTDLSK-a3-b3 | pr\|PROT03\|Twa1 | pr\|PROT03\|Twa1 | **intra-protein xl** | 210307_LU | 218 | 242 | 24 | 1359.817 | 454.28 | 3 | 1.1 | 0.35 | 0 | **34.45** |
| GVIEEPK-LAKLLK-a5-b3 | pr\|PROT03\|Twa1 | pr\|PROT03\|Twa1 | **intra-protein xl** | 210307_LU | 250 | 218 | 32 | 1436.898 | 360.232 | 4 | 1.2 | 0.4 | 0 | **34.44** |
| TKEVFHFSQAEK-EVFHFSQAEK-a2-b1 | pr\|PROT04\|MAEA | pr\|PROT04\|MAEA | **intra-protein xl** | 210307_LU | 405 | 406 | 1 | 2652.302 | 664.083 | 4 | 2 | 0.3 | 0 | **34.25** |
| RKAVESIQAEDESAK-EVEESLER-a2-b1 | pr\|PROT04\|MAEA | pr\|PROT04\|MAEA | **intra-protein xl** | 210307_LU | 107 | 188 | 81 | 2631.304 | 658.834 | 4 | 1.9 | 0.36 | 0 | **34.22** |
| MAVQESAAQLSMTLK-AAQKNIDR-a5-b4 | pr\|PROT04\|MAEA | pr\|PROT04\|MAEA | **intra-protein xl** | 210307_LU | 28 | 61 | 33 | 2503.292 | 835.438 | 3 | 1.2 | 0.37 | 0 | **34.21** |
| KHFSQAEGSQLDEVR-EVEESLER-a1-b1 | pr\|PROT04\|MAEA | pr\|PROT04\|MAEA | **intra-protein xl** | 210307_LU | 246 | 188 | 58 | 2701.302 | 676.333 | 4 | 2.7 | 0.34 | 0 | **34.17** |
| DKPIEMQLTSAK-DKPIEMQLTSAK-a1-b2 | pr\|PROT06\|ARMC8b | pr\|PROT06\|ARMC8b | **intra-protein xl** | 210307_LU | 257 | 258 | 1 | 2701.412 | 676.361 | 4 | 3.5 | 0.32 | 0 | **34.14** |
| NPNLLFTLKVR-ELQAMSEQLR-a9-b1 | pr\|PROT01\|RanBP9 | pr\|PROT01\|RanBP9 | **intra-protein xl** | 210307_LU | 469 | 650 | 181 | 2499.368 | 625.85 | 4 | 1.8 | 0.34 | 0 | **34.13** |
| XTDLSKGVIEEPK-LIXNYLVTEGFK-a6-b9 | pr\|PROT03\|Twa1 | pr\|PROT03\|Twa1 | **intra-protein xl** | 210307_LU | 245 | 64 | 181 | 2886.484 | 722.629 | 4 | 2.8 | 0.34 | 0 | **33.84** |
| VKYPKMTDLSK-GVIEEPK-a5-b4 | pr\|PROT03\|Twa1 | pr\|PROT03\|Twa1 | **intra-protein xl** | 210307_LU | 239 | 249 | 10 | 2061.122 | 516.288 | 4 | 1.7 | 0.3 | 0 | **33.68** |
| DGYMGIGLSAQGVNMNR-TPKDAASVR-a1-b3 | pr\|PROT01\|RanBP9 | pr\|PROT01\|RanBP9 | **intra-protein xl** | 210307_LU | 251 | 221 | 30 | 2707.326 | 903.45 | 3 | 3.2 | 0.48 | 0 | **33.55** |
| RKAVESIQAEDESAK-AVESIQAEDESAK-a2-b3 | pr\|PROT04\|MAEA | pr\|PROT04\|MAEA | **intra-protein xl** | 210307_LU | 107 | 110 | 3 | 3017.482 | 755.378 | 4 | 1.1 | 0.23 | 0 | **33.44** |
| KVSLGEGRPPVLTASR-LYASLGANDEDIR-a1-b11 | pr\|PROT06\|ARMC8b | pr\|PROT06\|ARMC8b | **intra-protein xl** | 210307_LU | 388 | 384 | 4 | 3083.647 | 617.737 | 5 | 3.5 | 0.3 | 0 | **33.44** |
| KVSLGEGRPPVLTASR-LYASLGANDEDIR-a1-b9 | pr\|PROT06\|ARMC8b | pr\|PROT06\|ARMC8b | **intra-protein xl** | 210307_LU | 388 | 382 | 6 | 3083.644 | 771.919 | 4 | 2.6 | 0.38 | 0 | **33.35** |
| MTDLSKGVIEEPK-LIXNYLVTEGFK-a6-b9 | pr\|PROT03\|Twa1 | pr\|PROT03\|Twa1 | **intra-protein xl** | 210307_LU | 245 | 64 | 181 | 2870.491 | 718.631 | 4 | 3.5 | 0.33 | 0 | **33.29** |
| XESGIEPSVDLETLDER-XTDLSKGVIEEPK-a6-b6 | pr\|PROT03\|Twa1 | pr\|PROT03\|Twa1 | **intra-protein xl** | 210307_LU | 80 | 245 | 165 | 3378.615 | 845.662 | 4 | 2.9 | 0.24 | 0 | **33.25** |
| AEWEGKGTASR-YLEYLEDGK-a6-b7 | pr\|PROT02\|WDR26 | pr\|PROT02\|WDR26 | **intra-protein xl** | 210307_LU | 178 | 130 | 48 | 2301.097 | 576.282 | 4 | 2.3 | 0.21 | 0 | **33.22** |
| VEGAETLAYLIEPDVELQR-NAVIGNNKQK-a5-b8 | pr\|PROT06\|ARMC8b | pr\|PROT06\|ARMC8b | **intra-protein xl** | 210307_LU | 312 | 62 | 250 | 3210.699 | 803.683 | 4 | 3.3 | 0.36 | 0 | **33.2** |
| RKAVESIQAEDESAK-ETSHVTXVVAELEK-a2-b1 | pr\|PROT04\|MAEA | pr\|PROT04\|MAEA | **intra-protein xl** | 210307_LU | 107 | 66 | 41 | 3229.621 | 808.413 | 4 | 2.2 | 0.25 | 0 | **33.06** |
| TECAVVLGSLAXGTENNVK-VLQGVIDMKNAVIGNNK-a2-b9 | pr\|PROT06\|ARMC8b | pr\|PROT06\|ARMC8b | **intra-protein xl** | 210307_LU | 92 | 54 | 38 | 3801.952 | 1268.325 | 3 | 2.4 | 0.31 | 0 | **33.02** |
| MTDLSKGVIEEPK-LLLWAQNELDQK-a6-b8 | pr\|PROT03\|Twa1 | pr\|PROT03\|Twa1 | **intra-protein xl** | 210307_LU | 245 | 229 | 16 | 2897.527 | 725.39 | 4 | 2.2 | 0.26 | 0 | **33.01** |
| MSYAEKPDEITK-DEWMEK-a6-b1 | pr\|PROT03\|Twa1 | pr\|PROT03\|Twa1 | **intra-protein xl** | 210307_LU | 30 | 37 | 7 | 2229.001 | 744.008 | 3 | 1.7 | 0.29 | 0 | **32.96** |
| LLLWAQNELDQKK-LLLWAQNELDQK-a12-b10 | pr\|PROT03\|Twa1 | pr\|PROT03\|Twa1 | **intra-protein xl** | 210307_LU | 233 | 231 | 2 | 3049.669 | 1017.564 | 3 | 3 | 0.31 | 0 | **32.82** |
| MSYAEKPDEITK-LAKLLK-a5-b3 | pr\|PROT03\|Twa1 | pr\|PROT03\|Twa1 | **intra-protein xl** | 210307_LU | 29 | 218 | 189 | 2077.152 | 416.438 | 5 | 1.3 | 0.26 | 0 | **32.68** |
| XTDLSKGVIEEPK-LLLWAQNELDQK-a6-b8 | pr\|PROT03\|Twa1 | pr\|PROT03\|Twa1 | **intra-protein xl** | 210307_LU | 245 | 229 | 16 | 2913.52 | 729.388 | 4 | 1.5 | 0.21 | 0 | **32.57** |
| IEHLKEHSSDQPAAASVWK-AVESIQAEDESAK-a5-b3 | pr\|PROT04\|MAEA | pr\|PROT04\|MAEA | **intra-protein xl** | 210307_LU | 130 | 110 | 20 | 3489.707 | 873.435 | 4 | 1.7 | 0.2 | 0 | **32.54** |
| RKAVESIQAEDESAK-EHSSDQPAAASVWK-a2-b5 | pr\|PROT04\|MAEA | pr\|PROT04\|MAEA | **intra-protein xl** | 210307_LU | 107 | 135 | 28 | 3153.54 | 631.716 | 5 | 2.2 | 0.19 | 0 | **32.53** |
| MTDLSKGVIEEPK-LIMNYLVTEGFK-a6-b9 | pr\|PROT03\|Twa1 | pr\|PROT03\|Twa1 | **intra-protein xl** | 210307_LU | 245 | 64 | 181 | 2854.49 | 714.63 | 4 | 1.2 | 0.22 | 0 | **32.34** |
| LLLWAQNELDQKK-LIXNYLVTEGFK-a12-b9 | pr\|PROT03\|Twa1 | pr\|PROT03\|Twa1 | **intra-protein xl** | 210307_LU | 233 | 64 | 169 | 3022.627 | 756.665 | 4 | 2.3 | 0.19 | 0 | **32.18** |
| KKVSLGEGRPPVLTASR-LYASLGANDEDIR-a2-b11 | pr\|PROT06\|ARMC8b | pr\|PROT06\|ARMC8b | **intra-protein xl** | 210307_LU | 388 | 384 | 4 | 3211.741 | 643.356 | 5 | 3 | 0.28 | 0.86 | **32.11** |
| LLNEVMVEHFFR-IKDTVQK-a8-b2 | pr\|PROT05\|RMND5A | pr\|PROT05\|RMND5A | **intra-protein xl** | 210307_LU | 148 | 93 | 55 | 2345.26 | 470.06 | 5 | 1.6 | 0.16 | 0 | **31.97** |
| LLLWAQNELDQKK-LLLWAQNELDQKK-a10-b12 | pr\|PROT03\|Twa1 | pr\|PROT03\|Twa1 | **intra-protein xl** | 210307_LU | 231 | 233 | 2 | 3177.763 | 795.448 | 4 | 2.4 | 0.25 | 0 | **31.83** |
| NAVIGNNKQK-MEVTASSR-a8-b2 | pr\|PROT06\|ARMC8b | pr\|PROT06\|ARMC8b | **intra-protein xl** | 210307_LU | 62 | 26 | 36 | 1946.004 | 649.676 | 3 | 1.7 | 0.34 | 0 | **31.6** |
| AKAEWEGKGTASR-YLEYLEDGK-a8-b7 | pr\|PROT02\|WDR26 | pr\|PROT02\|WDR26 | **intra-protein xl** | 210307_LU | 178 | 130 | 48 | 2500.231 | 626.066 | 4 | 3.2 | 0.31 | 0 | **31.54** |
| MSYAEKPDEITKDEWMEK-XSYAEKPDEITKDEWMEK-a6-b5 | pr\|PROT03\|Twa1 | pr\|PROT03\|Twa1 | **intra-protein xl** | 210307_LU | 30 | 29 | 1 | 4455.984 | 892.205 | 5 | 1.4 | 0.21 | 0 | **31.52** |
| AAQKNIDRETSHVTMVVAELEK-AVESIQAEDESAK-a4-b3 | pr\|PROT04\|MAEA | pr\|PROT04\|MAEA | **intra-protein xl** | 210307_LU | 61 | 110 | 49 | 3825.909 | 766.19 | 5 | 1 | 0.21 | 0 | **31.37** |
| XTDLSKGVIEEPK-LIMNYLVTEGFK-a6-b9 | pr\|PROT03\|Twa1 | pr\|PROT03\|Twa1 | **intra-protein xl** | 210307_LU | 245 | 64 | 181 | 2870.487 | 957.837 | 3 | 2.2 | 0.25 | 0 | **30.99** |
| KVSLGEGRPPVLTASR-LYASLGANDEDIRK-a1-b10 | pr\|PROT06\|ARMC8b | pr\|PROT06\|ARMC8b | **intra-protein xl** | 210307_LU | 388 | 383 | 5 | 3211.737 | 1071.587 | 3 | 1.7 | 0.19 | 0 | **30.89** |
| ETEAALEFAQTQLAEQGEESR-MTDLSKGVIEEPK-a7-b6 | pr\|PROT03\|Twa1 | pr\|PROT03\|Twa1 | **intra-protein xl** | 210307_LU | 147 | 245 | 98 | 3763.817 | 941.962 | 4 | 1.8 | 0.35 | 0 | **30.65** |
| EHSSDQPAAASVWK-KAVESIQAEDESAK-a5-b1 | pr\|PROT04\|MAEA | pr\|PROT04\|MAEA | **intra-protein xl** | 210307_LU | 135 | 107 | 28 | 2997.433 | 600.495 | 5 | 0.6 | 0.21 | 0 | **30.38** |
| GQIQEAIALINSLHPELLDTNR-XTDLSKGVIEEPK-a5-b6 | pr\|PROT03\|Twa1 | pr\|PROT03\|Twa1 | **intra-protein xl** | 210307_LU | 105 | 245 | 140 | 3888.042 | 973.018 | 4 | 3.1 | 0.27 | 0 | **29.7** |
| ETSHVTMVVAELEK-AAQKNIDR-a11-b4 | pr\|PROT04\|MAEA | pr\|PROT04\|MAEA | **intra-protein xl** | 210307_LU | 76 | 61 | 15 | 2468.273 | 618.076 | 4 | 1.7 | 0.16 | 0 | **29.69** |
| RKAVESIQAEDESAK-EVFHFSQAEK-a2-b1 | pr\|PROT04\|MAEA | pr\|PROT04\|MAEA | **intra-protein xl** | 210307_LU | 107 | 406 | 299 | 2862.42 | 716.613 | 4 | 1.9 | 0.19 | 0 | **28.88** |
| TECAVVLGSLAXGTENNVK-VLQGVIDXKNAVIGNNK-a2-b9 | pr\|PROT06\|ARMC8b | pr\|PROT06\|ARMC8b | **intra-protein xl** | 210307_LU | 92 | 54 | 38 | 3817.949 | 1273.657 | 3 | 2.9 | 0.28 | 0 | **28.77** |
| XESGIEPSVDLETLDER-LLLWAQNELDQKK-a12-b12 | pr\|PROT03\|Twa1 | pr\|PROT03\|Twa1 | **intra-protein xl** | 210307_LU | 86 | 233 | 147 | 3514.758 | 879.697 | 4 | 2.3 | 0.19 | 0.83 | **28.24** |
| VWSEVNQAVLDYENR-LLLWAQNELDQKK-a11-b12 | pr\|PROT03\|Twa1 | pr\|PROT03\|Twa1 | **intra-protein xl** | 210307_LU | 206 | 233 | 27 | 3400.746 | 1134.59 | 3 | 1.3 | 0.34 | 0 | **28.03** |
| MESGIEPSVDLETLDER-LLLWAQNELDQKK-a10-b12 | pr\|PROT03\|Twa1 | pr\|PROT03\|Twa1 | **intra-protein xl** | 210307_LU | 84 | 233 | 149 | 3498.769 | 875.7 | 4 | 3.9 | 0.19 | 0 | **27.92** |
| QKVWSEVNQAVLDYENR-XESGIEPSVDLETLDER-a2-b6 | pr\|PROT03\|Twa1 | pr\|PROT03\|Twa1 | **intra-protein xl** | 210307_LU | 195 | 80 | 115 | 3993.899 | 999.483 | 4 | 2.3 | 0.32 | 0 | **27.11** |
| ETEAALEFAQTQLAEQGEESR-LLLWAQNELDQKK-a7-b12 | pr\|PROT03\|Twa1 | pr\|PROT03\|Twa1 | **intra-protein xl** | 210307_LU | 147 | 233 | 86 | 3915.962 | 979.998 | 4 | 3.3 | 0.16 | 0 | **26.97** |
| VEGAETLAYLIEPDVELQR-DKPIEMQLTSAK-a5-b2 | pr\|PROT06\|ARMC8b | pr\|PROT06\|ARMC8b | **intra-protein xl** | 210307_LU | 312 | 258 | 54 | 3485.805 | 872.459 | 4 | 2.5 | 0.25 | 0 | **26.88** |
| MSYAEKPDEITK-ESTPKLAK-a8-b5 | pr\|PROT03\|Twa1 | pr\|PROT03\|Twa1 | **intra-protein xl** | 210307_LU | 32 | 215 | 183 | 2265.162 | 756.062 | 3 | 2.6 | 0.21 | 0 | **25.72** |
| ETEAALEFAQTQLAEQGEESR-MTDLSKGVIEEPK-a15-b6 | pr\|PROT03\|Twa1 | pr\|PROT03\|Twa1 | **intra-protein xl** | 210307_LU | 155 | 245 | 90 | 3763.825 | 941.964 | 4 | 4.1 | 0.21 | 0 | **24.88** |
| XESGIEPSVDLETLDER-XTDLSKGVIEEPK-a15-b6 | pr\|PROT03\|Twa1 | pr\|PROT03\|Twa1 | **intra-protein xl** | 210307_LU | 89 | 245 | 156 | 3378.616 | 845.662 | 4 | 3 | 0.18 | 0 | **23.63** |
| MSYAEKPDEITKDEWMEK-XSYAEKPDEITKDEWMEK-a5-b6 | pr\|PROT03\|Twa1 | pr\|PROT03\|Twa1 | **intra-protein xl** | 210307_LU | 29 | 30 | 1 | 4455.982 | 892.204 | 5 | 0.8 | 0.21 | 0 | **23.04** |
| XSYAEKPDEITKDEWMEK-ESTPKLAK-a5-b5 | pr\|PROT03\|Twa1 | pr\|PROT03\|Twa1 | **intra-protein xl** | 210307_LU | 29 | 215 | 186 | 3099.485 | 775.879 | 4 | 2.3 | 0.17 | 0 | **22.33** |
| VEGAETLAYLIEPDVELQR-QAAFKLYASLGANDEDIR-a5-b5 | pr\|PROT06\|ARMC8b | pr\|PROT06\|ARMC8b | **intra-protein xl** | 210307_LU | 312 | 373 | 61 | 4107.099 | 1027.782 | 4 | 4.5 | 0.4 | 0 | **22.2** |

### Low-res MS2

**Cross-linked peptide pairs in the 6mer complex identified using DMTMM as the cross-linking reagent and low-resolution MS/MS detection**

| Id | Protein1 | Protein2 | XLType | Spectrum | AbsPos1 | AbsPos2 | deltaAA | Mr | Mz | z | Error_rel[p | TIC | deltaS | Id-Score |
|---|---|---|---|---|---|---|---|---|---|---|---|---|---|---|
| VSLGEGRPPVLTASR-LAKLLK-a5-b3 | pr\|PROT06\|ARMC8b | pr\|PROT03\|Twa1 | **inter-protein xl** | 210307_LU | 393 | 218 | | 2204.341 | 552.093 | 4 | 2 | 0.37 | 0 | **29.59** |
| ELQAMSEQLR-AKAEWEGK-a7-b2 | pr\|PROT01\|RanBP9 | pr\|PROT02\|WDR26 | **inter-protein xl** | 210307_LU | 656 | 172 | | 2103.044 | 702.022 | 3 | 1 | 0.42 | 0.66 | **29.11** |
| IQAQIDR-LAKLLK-a6-b3 | pr\|PROT01\|RanBP9 | pr\|PROT03\|Twa1 | **inter-protein xl** | 210307_LU | 378 | 218 | | 1508.941 | 378.243 | 4 | 0.3 | 0.28 | 0 | **28.77** |
| EVFHFSQAEK-VGKAIDK-a1-b3 | pr\|PROT04\|MAEA | pr\|PROT05\|RMND5A | **inter-protein xl** | 210307_LU | 406 | 115 | | 1932.014 | 484.011 | 4 | 1.6 | 0.31 | 0.41 | **28.7** |
| KVSLGEGRPPVLTASR-GVIEEPK-a1-b4 | pr\|PROT06\|ARMC8b | pr\|PROT03\|Twa1 | **inter-protein xl** | 210307_LU | 388 | 249 | | 2418.365 | 605.599 | 4 | 2.3 | 0.38 | 0.39 | **26.98** |
| KVSLGEGRPPVLTASR-EVEESLER-a1-b1 | pr\|PROT06\|ARMC8b | pr\|PROT04\|MAEA | **inter-protein xl** | 210307_LU | 388 | 188 | | 2637.413 | 528.49 | 5 | 1.6 | 0.34 | 0.33 | **26.39** |
| HFSQAEGSQLDEVR-VGKAIDK-a12-b3 | pr\|PROT04\|MAEA | pr\|PROT05\|RMND5A | **inter-protein xl** | 210307_LU | 258 | 115 | | 2313.178 | 772.067 | 3 | 2.8 | 0.39 | 0 | **26.32** |
| QRETEAALEFAQTQLAEQGEESR-NAVIGNNKQK-a17-b8 | pr\|PROT03\|Twa1 | pr\|PROT06\|ARMC8b | **inter-protein xl** | 210307_LU | 155 | 62 | | 3686.833 | 922.716 | 4 | 2.2 | 0.34 | 0.46 | **26.26** |
| QKVWSEVNQAVLDYENR-EGEWQTMIQK-a2-b1 | pr\|PROT03\|Twa1 | pr\|PROT01\|RanBP9 | **inter-protein xl** | 210307_LU | 195 | 386 | | 3307.602 | 827.908 | 4 | 2.8 | 0.42 | 0.51 | **25.87** |
| QRETEAALEFAQTQLAEQGEESR-NAVIGNNKQK-a20-b8 | pr\|PROT03\|Twa1 | pr\|PROT06\|ARMC8b | **inter-protein xl** | 210307_LU | 158 | 62 | | 3686.835 | 922.717 | 4 | 2.9 | 0.3 | 0.37 | **24.86** |
| VSLGEGRPPVLTASR-ESTPKLAK-a5-b5 | pr\|PROT06\|ARMC8b | pr\|PROT03\|Twa1 | **inter-protein xl** | 210307_LU | 393 | 215 | | 2392.348 | 599.095 | 4 | 1.9 | 0.28 | 0.42 | **24.61** |
| ETEAALEFAQTQLAEQGEESR-TPKDAASVR-a7-b3 | pr\|PROT03\|Twa1 | pr\|PROT01\|RanBP9 | **inter-protein xl** | 210307_LU | 147 | 221 | | 3261.586 | 816.404 | 4 | 3.4 | 0.39 | 0.48 | **24.24** |
| QRETEAALEFAQTQLAEQGEESR-NAVIGNNKQK-a21-b8 | pr\|PROT03\|Twa1 | pr\|PROT06\|ARMC8b | **inter-protein xl** | 210307_LU | 159 | 62 | | 3686.835 | 738.375 | 5 | 2.7 | 0.35 | 0.45 | **23.66** |
| HFSQAEGSQLDEVR-VGKAIDK-a6-b3 | pr\|PROT04\|MAEA | pr\|PROT05\|RMND5A | **inter-protein xl** | 210307_LU | 252 | 115 | | 2313.175 | 579.302 | 4 | 1.5 | 0.22 | 0.47 | **23.6** |
| TKEVFHFSQAEK-AVIEQR-a2-b4 | pr\|PROT04\|MAEA | pr\|PROT05\|RMND5A | **inter-protein xl** | 210307_LU | 405 | 328 | | 2146.119 | 430.232 | 5 | 0.8 | 0.29 | 0.74 | **23.37** |
| ESTPKLAK-FIEACLR-a5-b3 | pr\|PROT03\|Twa1 | pr\|PROT06\|ARMC8b | **inter-protein xl** | 210307_LU | 215 | 133 | | 1761.946 | 441.494 | 4 | 0.8 | 0.22 | 0.53 | **22.9** |
| KVSLGEGRPPVLTASR-AVESIQAEDESAK-a1-b9 | pr\|PROT06\|ARMC8b | pr\|PROT04\|MAEA | **inter-protein xl** | 210307_LU | 388 | 116 | | 3023.599 | 605.728 | 5 | 3.6 | 0.31 | 0.82 | **22.29** |
| KVSLGEGRPPVLTASR-LIXNYLVTEGFK-a1-b9 | pr\|PROT06\|ARMC8b | pr\|PROT03\|Twa1 | **inter-protein xl** | 210307_LU | 388 | 64 | | 3090.699 | 619.148 | 5 | 2.9 | 0.35 | 0.43 | **22.14** |
| EVEESLER-VGKAIDK-a3-b3 | pr\|PROT04\|MAEA | pr\|PROT05\|RMND5A | **inter-protein xl** | 210307_LU | 190 | 115 | | 1700.897 | 426.232 | 4 | 1.5 | 0.19 | 0 | **22.03** |
| ETSHVTMVVAELEK-IKDTVQK-a11-b2 | pr\|PROT04\|MAEA | pr\|PROT05\|RMND5A | **inter-protein xl** | 210307_LU | 76 | 93 | | 2384.266 | 597.074 | 4 | 1.7 | 0.28 | 0.46 | **21.38** |
| ETEAALEFAQTQLAEQGEESR-NAVIGNNKQK-a18-b8 | pr\|PROT03\|Twa1 | pr\|PROT06\|ARMC8b | **inter-protein xl** | 210307_LU | 158 | 62 | | 3402.675 | 851.676 | 4 | 2.8 | 0.2 | 0.52 | **21.35** |
| HFSQAEGSQLDEVR-LASDHKDIHSSVSR-a12-b6 | pr\|PROT04\|MAEA | pr\|PROT05\|RMND5A | **inter-protein xl** | 210307_LU | 258 | 104 | | 3134.519 | 627.912 | 5 | 2 | 0.35 | 0.69 | **21.22** |
| RKAVESIQAEDESAK-EPFVELNR-a2-b1 | pr\|PROT04\|MAEA | pr\|PROT05\|RMND5A | **inter-protein xl** | 210307_LU | 107 | 176 | | 2644.355 | 662.096 | 4 | 3.4 | 0.27 | 0.53 | **20.96** |
| ELQAXSEQLR-AKAEWEGK-a7-b2 | pr\|PROT01\|RanBP9 | pr\|PROT02\|WDR26 | **inter-protein xl** | 210307_LU | 656 | 172 | | 2119.039 | 530.768 | 4 | 1.3 | 0.39 | 0 | **20.5** |
| AEWEGKGTASR-YLEYLEDGK-a6-b7 | pr\|PROT02\|WDR26 | pr\|PROT02\|WDR26 | **intra-protein xl** | 210307_LU | 178 | 130 | 48 | 2301.098 | 768.041 | 3 | 3 | 0.51 | 0 | **33.34** |
| XESGIEPSVDLETLDER-LAKLLK-a10-b3 | pr\|PROT03\|Twa1 | pr\|PROT03\|Twa1 | **intra-protein xl** | 210307_LU | 84 | 218 | 134 | 2601.364 | 651.349 | 4 | 2.5 | 0.43 | 0.55 | **32.6** |
| XSYAEKPDEITK-DEWMEK-a6-b1 | pr\|PROT03\|Twa1 | pr\|PROT03\|Twa1 | **intra-protein xl** | 210307_LU | 30 | 37 | 7 | 2244.996 | 749.34 | 3 | 1.9 | 0.29 | 0 | **32.04** |
| DELPIEVDLGKK-VGKAIDK-a2-b3 | pr\|PROT05\|RMND5A | pr\|PROT05\|RMND5A | **intra-protein xl** | 210307_LU | 341 | 115 | 226 | 2066.166 | 517.549 | 4 | 2 | 0.5 | 0.53 | **31.71** |
| TKEVFHFSQAEK-EVFHFSQAEK-a2-b1 | pr\|PROT04\|MAEA | pr\|PROT04\|MAEA | **intra-protein xl** | 210307_LU | 405 | 406 | 1 | 2652.302 | 885.108 | 3 | 1.6 | 0.21 | 0 | **31.63** |
| LAKLLK-MTDLSK-a3-b3 | pr\|PROT03\|Twa1 | pr\|PROT03\|Twa1 | **intra-protein xl** | 210307_LU | 218 | 242 | 24 | 1359.817 | 454.28 | 3 | 0.8 | 0.37 | 0 | **31.59** |
| YPKMTDLSK-GVIEEPK-a3-b5 | pr\|PROT03\|Twa1 | pr\|PROT03\|Twa1 | **intra-protein xl** | 210307_LU | 239 | 250 | 11 | 1833.959 | 612.328 | 3 | 2.6 | 0.26 | 0.68 | **31.36** |
| GVIEEPK-LAKLLK-a5-b3 | pr\|PROT03\|Twa1 | pr\|PROT03\|Twa1 | **intra-protein xl** | 210307_LU | 250 | 218 | 32 | 1436.898 | 360.232 | 4 | 1 | 0.62 | 0 | **31.34** |
| LLLWAQNELDQKK-LLLWAQNELDQKK-a10-b12 | pr\|PROT03\|Twa1 | pr\|PROT03\|Twa1 | **intra-protein xl** | 210307_LU | 231 | 233 | 2 | 3177.763 | 795.448 | 4 | 2.4 | 0.43 | 0.25 | **31.14** |
| DKPIEMQLTSAK-DKPIEMQLTSAK-a2-b5 | pr\|PROT06\|ARMC8b | pr\|PROT06\|ARMC8b | **intra-protein xl** | 210307_LU | 258 | 261 | 3 | 2701.405 | 676.359 | 4 | 0.9 | 0.42 | 0.29 | **30.38** |
| TKEVFHFSQAEK-VQEYPTLK-a2-b3 | pr\|PROT04\|MAEA | pr\|PROT04\|MAEA | **intra-protein xl** | 210307_LU | 405 | 41 | 364 | 2408.239 | 482.656 | 5 | 0.7 | 0.28 | 0.63 | **30.17** |
| LLLWAQNELDQKK-LLLWAQNELDQK-a12-b10 | pr\|PROT03\|Twa1 | pr\|PROT03\|Twa1 | **intra-protein xl** | 210307_LU | 233 | 231 | 2 | 3049.666 | 763.424 | 4 | 1.9 | 0.22 | 0.23 | **30.13** |
| MTDLSKGVIEEPK-GVIEEPK-a6-b4 | pr\|PROT03\|Twa1 | pr\|PROT03\|Twa1 | **intra-protein xl** | 210307_LU | 245 | 249 | 4 | 2198.153 | 733.726 | 3 | 1.4 | 0.39 | 0 | **30.12** |
| MAVQESAAQLSMTLK-AAQKNIDR-a5-b4 | pr\|PROT04\|MAEA | pr\|PROT04\|MAEA | **intra-protein xl** | 210307_LU | 28 | 61 | 33 | 2503.293 | 835.439 | 3 | 1.8 | 0.38 | 0 | **29.48** |
| GVIEEPK-LAKLLK-a4-b3 | pr\|PROT03\|Twa1 | pr\|PROT03\|Twa1 | **intra-protein xl** | 210307_LU | 249 | 218 | 31 | 1436.899 | 479.974 | 3 | 1.3 | 0.4 | 0 | **29.46** |
| XESGIEPSVDLETLDER-LAKLLK-a12-b3 | pr\|PROT03\|Twa1 | pr\|PROT03\|Twa1 | **intra-protein xl** | 210307_LU | 86 | 218 | 132 | 2601.363 | 651.349 | 4 | 2.4 | 0.44 | 0.54 | **28.99** |
| YPSSVSAITDIKR-LLEER-a12-b3 | pr\|PROT06\|ARMC8b | pr\|PROT06\|ARMC8b | **intra-protein xl** | 210307_LU | 355 | 305 | 50 | 2076.123 | 520.039 | 4 | 0.7 | 0.33 | 0 | **28.83** |
| KAVESIQAEDESAK-EVEESLER-a1-b1 | pr\|PROT04\|MAEA | pr\|PROT04\|MAEA | **intra-protein xl** | 210307_LU | 107 | 188 | 81 | 2475.201 | 619.808 | 4 | 1.2 | 0.36 | 0 | **28.77** |
| DGYMGIGLSAQGVNMNR-TPKDAASVR-a1-b3 | pr\|PROT01\|RanBP9 | pr\|PROT01\|RanBP9 | **intra-protein xl** | 210307_LU | 251 | 221 | 30 | 2707.324 | 903.449 | 3 | 2.7 | 0.44 | 0.43 | **28.56** |
| MSYAEKPDEITK-DEWMEK-a6-b1 | pr\|PROT03\|Twa1 | pr\|PROT03\|Twa1 | **intra-protein xl** | 210307_LU | 30 | 37 | 7 | 2229 | 744.008 | 3 | 1.4 | 0.38 | 0 | **28.41** |
| RKAVESIQAEDESAK-EVEESLER-a2-b1 | pr\|PROT04\|MAEA | pr\|PROT04\|MAEA | **intra-protein xl** | 210307_LU | 107 | 188 | 81 | 2631.304 | 658.834 | 4 | 2 | 0.5 | 0.31 | **28.23** |
| DKPIEMQLTSAK-LLEER-a2-b4 | pr\|PROT06\|ARMC8b | pr\|PROT06\|ARMC8b | **intra-protein xl** | 210307_LU | 258 | 306 | 48 | 2000.066 | 501.024 | 4 | 2.3 | 0.51 | 0.46 | **28.05** |
| XSYAEKPDEITK-DEWXEK-a6-b1 | pr\|PROT03\|Twa1 | pr\|PROT03\|Twa1 | **intra-protein xl** | 210307_LU | 30 | 37 | 7 | 2260.992 | 754.672 | 3 | 2.5 | 0.23 | 0 | **27.76** |
| KVSLGEGRPPVLTASR-LLEER-a1-b3 | pr\|PROT06\|ARMC8b | pr\|PROT06\|ARMC8b | **intra-protein xl** | 210307_LU | 388 | 305 | 83 | 2306.311 | 577.586 | 4 | 1.6 | 0.39 | 0.72 | **27.57** |
| LASDHKDIHSSVSR-DELPIEVDLGKK-a6-b2 | pr\|PROT05\|RMND5A | pr\|PROT05\|RMND5A | **intra-protein xl** | 210307_LU | 104 | 341 | 237 | 2887.507 | 482.259 | 6 | 1.2 | 0.44 | 0.68 | **27.44** |
| MSYAEKPDEITK-ESTPKLAK-a8-b5 | pr\|PROT03\|Twa1 | pr\|PROT03\|Twa1 | **intra-protein xl** | 210307_LU | 32 | 215 | 183 | 2265.162 | 756.062 | 3 | 2.7 | 0.34 | 0 | **27.35** |
| AKAEWEGKGTASR-YLEYLEDGK-a8-b7 | pr\|PROT02\|WDR26 | pr\|PROT02\|WDR26 | **intra-protein xl** | 210307_LU | 178 | 130 | 48 | 2500.229 | 834.418 | 3 | 2.4 | 0.42 | 0 | **27.35** |
| KHFSQAEGSQLDEVR-EVEESLER-a1-b1 | pr\|PROT04\|MAEA | pr\|PROT04\|MAEA | **intra-protein xl** | 210307_LU | 246 | 188 | 58 | 2701.301 | 676.333 | 4 | 2.6 | 0.32 | 0.28 | **27.34** |
| KVSLGEGRPPVLTASR-LYASLGANDEDIR-a1-b11 | pr\|PROT06\|ARMC8b | pr\|PROT06\|ARMC8b | **intra-protein xl** | 210307_LU | 388 | 384 | 4 | 3083.646 | 617.737 | 5 | 3 | 0.42 | 0.79 | **27.18** |
| ESTPKLAK-GVIEEPK-a5-b5 | pr\|PROT03\|Twa1 | pr\|PROT03\|Twa1 | **intra-protein xl** | 210307_LU | 215 | 250 | 35 | 1624.905 | 542.643 | 3 | 1 | 0.23 | 0 | **27.08** |
| XTDLSKGVIEEPK-LIXNYLVTEGFK-a6-b9 | pr\|PROT03\|Twa1 | pr\|PROT03\|Twa1 | **intra-protein xl** | 210307_LU | 245 | 64 | 181 | 2886.488 | 722.63 | 4 | 4.4 | 0.36 | 0.37 | **26.92** |
| VKYPKMTDLSK-GVIEEPK-a5-b5 | pr\|PROT03\|Twa1 | pr\|PROT03\|Twa1 | **intra-protein xl** | 210307_LU | 239 | 250 | 11 | 2061.121 | 413.232 | 5 | 1.4 | 0.37 | 0.75 | **26.74** |
| AEWEGKGTASR-YLEYLEDGK-a6-b6 | pr\|PROT02\|WDR26 | pr\|PROT02\|WDR26 | **intra-protein xl** | 210307_LU | 178 | 129 | 49 | 2301.095 | 576.281 | 4 | 1.5 | 0.27 | 0 | **26.72** |
| MSYAEKPDEITKDEWMEK-ESTPKLAK-a5-b5 | pr\|PROT03\|Twa1 | pr\|PROT03\|Twa1 | **intra-protein xl** | 210307_LU | 29 | 215 | 186 | 3083.491 | 771.881 | 4 | 2.5 | 0.33 | 0.21 | **26.71** |
| VWSEVNQAVLDYENR-MTDLSKGVIEEPK-a11-b6 | pr\|PROT03\|Twa1 | pr\|PROT03\|Twa1 | **intra-protein xl** | 210307_LU | 206 | 245 | 39 | 3248.606 | 813.159 | 4 | 1.2 | 0.39 | 0.74 | **26.52** |
| KVSLGEGRPPVLTASR-LYASLGANDEDIR-a1-b10 | pr\|PROT06\|ARMC8b | pr\|PROT06\|ARMC8b | **intra-protein xl** | 210307_LU | 388 | 383 | 5 | 3083.645 | 617.737 | 5 | 2.8 | 0.3 | 0.74 | **26.33** |
| LIMNYLVTEGFK-ESTPKLAK-a9-b5 | pr\|PROT03\|Twa1 | pr\|PROT03\|Twa1 | **intra-protein xl** | 210307_LU | 64 | 215 | 151 | 2281.246 | 761.423 | 3 | 3.2 | 0.32 | 0.6 | **26.29** |
| MESGIEPSVDLETLDER-VKYPK-a12-b2 | pr\|PROT03\|Twa1 | pr\|PROT03\|Twa1 | **intra-protein xl** | 210307_LU | 86 | 236 | 150 | 2534.263 | 634.574 | 4 | 2.3 | 0.31 | 0.51 | **25.93** |
| XTDLSKGVIEEPK-LLLWAQNELDQK-a6-b8 | pr\|PROT03\|Twa1 | pr\|PROT03\|Twa1 | **intra-protein xl** | 210307_LU | 245 | 229 | 16 | 2913.527 | 729.39 | 4 | 3.8 | 0.23 | 0.53 | **25.82** |
| LLLWAQNELDQKK-LIXNYLVTEGFK-a12-b9 | pr\|PROT03\|Twa1 | pr\|PROT03\|Twa1 | **intra-protein xl** | 210307_LU | 233 | 64 | 169 | 3022.627 | 756.665 | 4 | 2.3 | 0.31 | 0.43 | **25.78** |
| VKYPKMTDLSK-GVIEEPK-a5-b4 | pr\|PROT03\|Twa1 | pr\|PROT03\|Twa1 | **intra-protein xl** | 210307_LU | 239 | 249 | 10 | 2061.122 | 516.288 | 4 | 2 | 0.2 | 0.63 | **25.75** |
| ETSHVTMVVAELEK-KAVESIQAEDESAK-a1-b1 | pr\|PROT04\|MAEA | pr\|PROT04\|MAEA | **intra-protein xl** | 210307_LU | 66 | 107 | 41 | 3057.528 | 765.39 | 4 | 3.5 | 0.45 | 0.39 | **25.71** |
| EHSSDQPAAASVWKR-VQEYPTLK-a14-b3 | pr\|PROT04\|MAEA | pr\|PROT04\|MAEA | **intra-protein xl** | 210307_LU | 144 | 41 | 103 | 2626.317 | 526.271 | 5 | 1.2 | 0.26 | 0.59 | **25.42** |
| MTDLSKGVIEEPK-LIXNYLVTEGFK-a6-b9 | pr\|PROT03\|Twa1 | pr\|PROT03\|Twa1 | **intra-protein xl** | 210307_LU | 245 | 64 | 181 | 2870.49 | 718.63 | 4 | 3.3 | 0.38 | 0.62 | **25.38** |
| DKPIEMQLTSAK-LLEER-a2-b3 | pr\|PROT06\|ARMC8b | pr\|PROT06\|ARMC8b | **intra-protein xl** | 210307_LU | 258 | 305 | 47 | 2000.064 | 501.024 | 4 | 1.5 | 0.45 | 0.31 | **25.27** |
| MSYAEKPDEITK-VKYPK-a9-b2 | pr\|PROT03\|Twa1 | pr\|PROT03\|Twa1 | **intra-protein xl** | 210307_LU | 33 | 236 | 203 | 2026.05 | 676.358 | 3 | 2.9 | 0.32 | 0 | **25.08** |

| | | | | | | | | | | | | | | |
|---|---|---|---|---|---|---|---|---|---|---|---|---|---|---|
| NPNLLFTLKVR-ELQAMSEQLR-a9-b1 | pr\|PROT01\|RanBP9 | pr\|PROT01\|RanBP9 | **intra-protein xl** | 210307_LU | 469 | 650 | 181 | 2499.37 | 834.131 | 3 | 2.8 | 0.31 | 0 | **24.75** |
| VLHKFSGYGQLCER-DIHSSVSR-a4-b1 | pr\|PROT05\|RMND5A | pr\|PROT05\|RMND5A | **intra-protein xl** | 210307_LU | 41 | 105 | 64 | 2574.28 | 644.578 | 4 | 1.3 | 0.36 | 0.31 | **24.66** |
| VGKAIDK-ILEALK-a3-b3 | pr\|PROT05\|RMND5A | pr\|PROT05\|RMND5A | **intra-protein xl** | 210307_LU | 115 | 186 | 71 | 1396.867 | 466.63 | 3 | 0.9 | 0.26 | 0 | **24.64** |
| MTDLSKGVIEEPK-LIMNYLVTEGFK-a6-b9 | pr\|PROT03\|Twa1 | pr\|PROT03\|Twa1 | **intra-protein xl** | 210307_LU | 245 | 64 | 181 | 2854.493 | 714.631 | 4 | 2.6 | 0.37 | 0.56 | **24.61** |
| KVSLGEGRPPVLTASR-LLEER-a1-b4 | pr\|PROT06\|ARMC8b | pr\|PROT06\|ARMC8b | **intra-protein xl** | 210307_LU | 388 | 306 | 82 | 2306.311 | 577.586 | 4 | 1.9 | 0.27 | 0.24 | **24.58** |
| MTDLSKGVIEEPK-LLLWAQNELDQK-a6-b10 | pr\|PROT03\|Twa1 | pr\|PROT03\|Twa1 | **intra-protein xl** | 210307_LU | 245 | 231 | 14 | 2897.524 | 725.389 | 4 | 1.1 | 0.27 | 0.6 | **24.14** |
| NPNLLFTLKVR-ELQAXSEQLR-a9-b1 | pr\|PROT01\|RanBP9 | pr\|PROT01\|RanBP9 | **intra-protein xl** | 210307_LU | 469 | 650 | 181 | 2515.362 | 629.848 | 4 | 1.7 | 0.24 | 0 | **23.84** |
| RKAVESIQAEDESAK-EHSSDQPAAASVWK-a2-b1 | pr\|PROT04\|MAEA | pr\|PROT04\|MAEA | **intra-protein xl** | 210307_LU | 107 | 131 | 24 | 3153.538 | 789.392 | 4 | 1.6 | 0.29 | 0.65 | **23.68** |
| LIMNYLVTEGFK-KVKYPK-a9-b1 | pr\|PROT03\|Twa1 | pr\|PROT03\|Twa1 | **intra-protein xl** | 210307_LU | 64 | 234 | 170 | 2170.225 | 435.053 | 5 | 1.3 | 0.61 | 0.61 | **23.32** |
| LLNEVMVEHFFR-VGKAIDK-a4-b3 | pr\|PROT05\|RMND5A | pr\|PROT05\|RMND5A | **intra-protein xl** | 210307_LU | 144 | 115 | 29 | 2244.211 | 449.85 | 5 | 0.8 | 0.36 | 0.74 | **23.29** |
| QKVWSEVNQAVLDYENR-LLLWAQNELDQK-a2-b8 | pr\|PROT03\|Twa1 | pr\|PROT03\|Twa1 | **intra-protein xl** | 210307_LU | 195 | 229 | 34 | 3528.809 | 883.21 | 4 | 2.4 | 0.35 | 0.43 | **23.28** |
| ETSHVTMVVAELEK-AAQKNIDR-a11-b4 | pr\|PROT04\|MAEA | pr\|PROT04\|MAEA | **intra-protein xl** | 210307_LU | 76 | 61 | 15 | 2468.275 | 618.077 | 4 | 2.4 | 0.2 | 0 | **23.24** |
| MSYAEKPDEITK-LAKLLK-a8-b3 | pr\|PROT03\|Twa1 | pr\|PROT03\|Twa1 | **intra-protein xl** | 210307_LU | 32 | 218 | 186 | 2077.15 | 416.438 | 5 | 0.4 | 0.39 | 0.44 | **23.13** |
| VEGAETLAYLIEPDVELQR-QKANLIVLGAVPR-a5-b2 | pr\|PROT06\|ARMC8b | pr\|PROT06\|ARMC8b | **intra-protein xl** | 210307_LU | 312 | 64 | 248 | 3503.944 | 876.994 | 4 | 2.4 | 0.36 | 0.52 | **23.13** |
| AVESIQAEDESAK-TKEVFHFSQAEK-a8-b2 | pr\|PROT04\|MAEA | pr\|PROT04\|MAEA | **intra-protein xl** | 210307_LU | 115 | 405 | 290 | 2807.368 | 702.85 | 4 | 2.3 | 0.24 | 0.85 | **22.98** |
| EHSSDQPAAASVWKR-AVESIQAEDESAK-a14-b10 | pr\|PROT04\|MAEA | pr\|PROT04\|MAEA | **intra-protein xl** | 210307_LU | 144 | 117 | 27 | 3025.444 | 606.097 | 5 | 2.1 | 0.33 | 0.89 | **22.79** |
| RKAVESIQAEDESAK-ETSHVTXVVAELEK-a2-b1 | pr\|PROT04\|MAEA | pr\|PROT04\|MAEA | **intra-protein xl** | 210307_LU | 107 | 66 | 41 | 3229.626 | 646.933 | 5 | 3.9 | 0.4 | 0.56 | **22.77** |
| GQIQEAIALINSLHPELLDTNR-XTDLSKGVIEEPK-a5-b6 | pr\|PROT03\|Twa1 | pr\|PROT03\|Twa1 | **intra-protein xl** | 210307_LU | 105 | 245 | 140 | 3888.037 | 778.615 | 5 | 1.8 | 0.49 | 0.42 | **22.72** |
| AVESIQAEDESAK-LSVLKR-a8-b5 | pr\|PROT04\|MAEA | pr\|PROT04\|MAEA | **intra-protein xl** | 210307_LU | 115 | 105 | 10 | 2072.114 | 519.036 | 4 | 1.2 | 0.29 | 0.6 | **22.72** |
| RKAVESIQAEDESAK-HFSQAEGSQLDEVR-a2-b12 | pr\|PROT04\|MAEA | pr\|PROT04\|MAEA | **intra-protein xl** | 210307_LU | 107 | 258 | 151 | 3243.581 | 649.724 | 5 | 1.6 | 0.26 | 0.52 | **22.19** |
| XAVQESAAQLSMTLK-AAQKNIDR-a5-b4 | pr\|PROT04\|MAEA | pr\|PROT04\|MAEA | **intra-protein xl** | 210307_LU | 28 | 61 | 33 | 2519.288 | 630.83 | 4 | 1.7 | 0.21 | 0.65 | **22.18** |
| KVSLGEGRPPVLTASR-MEVTASSR-a1-b2 | pr\|PROT06\|ARMC8b | pr\|PROT06\|ARMC8b | **intra-protein xl** | 210307_LU | 388 | 26 | 362 | 2527.359 | 843.461 | 3 | 2 | 0.18 | 0.52 | **22.17** |
| MESGIEPSVDLETLDER-LLLWAQNELDQKK-a10-b12 | pr\|PROT03\|Twa1 | pr\|PROT03\|Twa1 | **intra-protein xl** | 210307_LU | 84 | 233 | 149 | 3498.769 | 875.7 | 4 | 4 | 0.3 | 0.77 | **22.1** |
| KKVSLGEGRPPVLTASR-LYASLGANDEDIR-a2-b11 | pr\|PROT06\|ARMC8b | pr\|PROT06\|ARMC8b | **intra-protein xl** | 210307_LU | 388 | 384 | 4 | 3211.733 | 643.354 | 5 | 0.5 | 0.44 | 0.84 | **22.05** |
| QKVWSEVNQAVLDYENR-XESGIEPSVDLETLDER-a2-b6 | pr\|PROT03\|Twa1 | pr\|PROT03\|Twa1 | **intra-protein xl** | 210307_LU | 195 | 80 | 115 | 3993.905 | 999.484 | 4 | 3.7 | 0.35 | 0.46 | **21.85** |
| VQCLWCLSDGKTVLASDTHQR-GYNFEDLTDR-a11-b5 | pr\|PROT02\|WDR26 | pr\|PROT02\|WDR26 | **intra-protein xl** | 210307_LU | 402 | 419 | 17 | 3683.719 | 921.938 | 4 | 2.5 | 0.21 | 0.82 | **21.81** |
| ETEAALEFAQTQLAEQGEESR-VKYPKXTDLSK-a7-b5 | pr\|PROT03\|Twa1 | pr\|PROT03\|Twa1 | **intra-protein xl** | 210307_LU | 147 | 239 | 92 | 3642.782 | 729.564 | 5 | 2.6 | 0.35 | 0.76 | **21.69** |
| DEWMEK-LAKLLK-a2-b3 | pr\|PROT03\|Twa1 | pr\|PROT03\|Twa1 | **intra-protein xl** | 210307_LU | 38 | 218 | 180 | 1502.821 | 501.948 | 3 | 2.8 | 0.19 | 0 | **21.62** |
| EHSSDQPAAASVWKR-AVESIQAEDESAK-a14-b8 | pr\|PROT04\|MAEA | pr\|PROT04\|MAEA | **intra-protein xl** | 210307_LU | 144 | 115 | 29 | 3025.443 | 757.369 | 4 | 1.9 | 0.4 | 0.36 | **21.55** |
| KVSLGEGRPPVLTASR-LYASLGANDEDIR-a1-b9 | pr\|PROT06\|ARMC8b | pr\|PROT06\|ARMC8b | **intra-protein xl** | 210307_LU | 388 | 382 | 6 | 3083.645 | 617.737 | 5 | 2.8 | 0.29 | 0.81 | **21.55** |
| LIMNYLVTEGFKEAAEK-MESGIEPSVDLETLDER-a12-b12 | pr\|PROT03\|Twa1 | pr\|PROT03\|Twa1 | **intra-protein xl** | 210307_LU | 67 | 86 | 19 | 3855.892 | 964.981 | 4 | 3 | 0.29 | 0.71 | **21.45** |
| MESGIEPSVDLETLDER-ESTPKLAK-a10-b5 | pr\|PROT03\|Twa1 | pr\|PROT03\|Twa1 | **intra-protein xl** | 210307_LU | 84 | 215 | 131 | 2773.374 | 694.351 | 4 | 1.9 | 0.28 | 0.5 | **21.4** |
| YPKXTDLSK-GVIEEPK-a3-b4 | pr\|PROT03\|Twa1 | pr\|PROT03\|Twa1 | **intra-protein xl** | 210307_LU | 239 | 249 | 10 | 1849.951 | 463.496 | 4 | 1 | 0.18 | 0 | **21.34** |
| LLLWAQNELDQKK-LAKLLK-a8-b3 | pr\|PROT03\|Twa1 | pr\|PROT03\|Twa1 | **intra-protein xl** | 210307_LU | 229 | 218 | 11 | 2264.367 | 453.881 | 5 | 2.2 | 0.16 | 0.69 | **21.27** |
| MSYAEKPDEITK-DEWXEK-a6-b1 | pr\|PROT03\|Twa1 | pr\|PROT03\|Twa1 | **intra-protein xl** | 210307_LU | 30 | 37 | 7 | 2244.995 | 562.257 | 4 | 1.4 | 0.54 | 0.5 | **20.9** |
| LIMNYLVTEGFKEAAEK-XESGIEPSVDLETLDER-a12-b12 | pr\|PROT03\|Twa1 | pr\|PROT03\|Twa1 | **intra-protein xl** | 210307_LU | 67 | 86 | 19 | 3871.889 | 968.98 | 4 | 3.6 | 0.19 | 0.3 | **20.82** |
| VEGAETLAYLIEPDVELQR-DKPIEMQLTSAK-a12-b2 | pr\|PROT06\|ARMC8b | pr\|PROT06\|ARMC8b | **intra-protein xl** | 210307_LU | 319 | 258 | 61 | 3485.812 | 872.461 | 4 | 4.6 | 0.25 | 0.51 | **20.65** |
| TECAVVLGSLAXGTENNVK-VLQGVIDMKNAVIGNNK-a2-b9 | pr\|PROT06\|ARMC8b | pr\|PROT06\|ARMC8b | **intra-protein xl** | 210307_LU | 92 | 54 | 38 | 3801.952 | 951.496 | 4 | 2.4 | 0.31 | 0.63 | **20.49** |
| RKAVESIQAEDESAK-VQEYPTLK-a2-b3 | pr\|PROT04\|MAEA | pr\|PROT04\|MAEA | **intra-protein xl** | 210307_LU | 107 | 41 | 66 | 2618.36 | 655.598 | 4 | 1.7 | 0.32 | 0.58 | **20.49** |
| LLYLLQQETSSTELK-NAVIGNNKQK-a8-b8 | pr\|PROT06\|ARMC8b | pr\|PROT06\|ARMC8b | **intra-protein xl** | 210307_LU | 83 | 62 | 21 | 2831.549 | 708.895 | 4 | 3.5 | 0.19 | 0.38 | **20.32** |
| XESGIEPSVDLETLDER-LLLWAQNELDQKK-a12-b12 | pr\|PROT03\|Twa1 | pr\|PROT03\|Twa1 | **intra-protein xl** | 210307_LU | 86 | 233 | 147 | 3514.764 | 879.699 | 4 | 3.8 | 0.26 | 0.55 | **20.24** |
| QRETEAALEFAQTQLAEQGEESR-MTDLSKGVIEEPK-a9-b6 | pr\|PROT03\|Twa1 | pr\|PROT03\|Twa1 | **intra-protein xl** | 210307_LU | 147 | 245 | 98 | 4047.979 | 810.604 | 5 | 2.4 | 0.33 | 0.57 | **20.19** |
| VWSEVNQAVLDYENR-LLLWAQNELDQKK-a4-b12 | pr\|PROT03\|Twa1 | pr\|PROT03\|Twa1 | **intra-protein xl** | 210307_LU | 199 | 233 | 34 | 3400.749 | 851.195 | 4 | 2.3 | 0.23 | 0.42 | **20.09** |
| KVSLGEGRPPVLTASR-LYASLGANDEDIRK-a1-b9 | pr\|PROT06\|ARMC8b | pr\|PROT06\|ARMC8b | **intra-protein xl** | 210307_LU | 388 | 382 | 6 | 3211.74 | 643.356 | 5 | 2.7 | 0.46 | 0.89 | **20.09** |
| TECAVVLGSLAXGTENNVK-VLQGVIDXKNAVIGNNK-a2-b9 | pr\|PROT06\|ARMC8b | pr\|PROT06\|ARMC8b | **intra-protein xl** | 210307_LU | 92 | 54 | 38 | 3817.956 | 955.497 | 4 | 4.9 | 0.32 | 0.33 | **19.99** |
| LIMNYLVTEGFK-KVKYPK-a9-b3 | pr\|PROT03\|Twa1 | pr\|PROT03\|Twa1 | **intra-protein xl** | 210307_LU | 64 | 236 | 172 | 2170.224 | 543.564 | 4 | 0.7 | 0.61 | 0.81 | **19.89** |
| KVSLGEGRPPVLTASR-VSLGEGRPPVLTASR-a1-b5 | pr\|PROT06\|ARMC8b | pr\|PROT06\|ARMC8b | **intra-protein xl** | 210307_LU | 388 | 393 | 5 | 3185.81 | 638.17 | 5 | 3.1 | 0.35 | 0 | **19.87** |
| MSYAEKPDEITK-DEWMEK-a6-b2 | pr\|PROT03\|Twa1 | pr\|PROT03\|Twa1 | **intra-protein xl** | 210307_LU | 30 | 38 | 8 | 2229 | 744.008 | 3 | 1.5 | 0.21 | 0 | **19.71** |
| DKPIEXQLTSAK-FIEACLR-a2-b3 | pr\|PROT06\|ARMC8b | pr\|PROT06\|ARMC8b | **intra-protein xl** | 210307_LU | 258 | 133 | 125 | 2265.158 | 454.039 | 5 | 3.7 | 0.16 | 0.65 | **19.61** |
| MSYAEKPDEITKDEWMEK-XSYAEKPDEITKDEWMEK-a5-b12 | pr\|PROT03\|Twa1 | pr\|PROT03\|Twa1 | **intra-protein xl** | 210307_LU | 29 | 36 | 7 | 4455.991 | 637.578 | 7 | 2.9 | 0.27 | 0.83 | **19.51** |
| VWSEVNQAVLDYENR-ESTPKLAK-a4-b5 | pr\|PROT03\|Twa1 | pr\|PROT03\|Twa1 | **intra-protein xl** | 210307_LU | 199 | 215 | 16 | 2675.36 | 669.848 | 4 | 1.7 | 0.29 | 0.35 | **19.45** |
| LLLWAQNELDQK-YPKMTDLSK-a8-b3 | pr\|PROT03\|Twa1 | pr\|PROT03\|Twa1 | **intra-protein xl** | 210307_LU | 229 | 239 | 10 | 2533.331 | 845.451 | 3 | 2.2 | 0.34 | 0.38 | **19.44** |
| ETEAALEFAQTQLAEQGEESR-XTDLSKGVIEEPK-a7-b6 | pr\|PROT03\|Twa1 | pr\|PROT03\|Twa1 | **intra-protein xl** | 210307_LU | 147 | 245 | 98 | 3779.817 | 945.962 | 4 | 3.3 | 0.27 | 0.53 | **19.34** |
| MSYAEKPDEITKDEWMEK-LLLWAQNELDQKK-a9-b12 | pr\|PROT03\|Twa1 | pr\|PROT03\|Twa1 | **intra-protein xl** | 210307_LU | 33 | 233 | 200 | 3808.874 | 762.783 | 5 | 1.1 | 0.25 | 0.52 | **19.3** |
| LLLWAQNELDQKK-ESTPKLAK-a8-b5 | pr\|PROT03\|Twa1 | pr\|PROT03\|Twa1 | **intra-protein xl** | 210307_LU | 229 | 215 | 14 | 2452.371 | 491.482 | 5 | 0.8 | 0.21 | 0.71 | **19.23** |
| VSLGEGRPPVLTASR-NAVIGNNKQK-a5-b8 | pr\|PROT06\|ARMC8b | pr\|PROT06\|ARMC8b | **intra-protein xl** | 210307_LU | 393 | 62 | 331 | 2604.452 | 521.898 | 5 | 2.1 | 0.27 | 0.61 | **19.22** |
| ETSHVTXVVAELEK-AAQKNIDR-a1-b4 | pr\|PROT04\|MAEA | pr\|PROT04\|MAEA | **intra-protein xl** | 210307_LU | 66 | 61 | 5 | 2484.268 | 497.861 | 5 | 1.6 | 0.3 | 0.56 | **19.02** |
| XSYAEKPDEITKDEWMEK-LAKLLK-a8-b3 | pr\|PROT03\|Twa1 | pr\|PROT03\|Twa1 | **intra-protein xl** | 210307_LU | 32 | 218 | 186 | 2911.479 | 486.254 | 6 | 2.7 | 0.21 | 0.88 | **18.76** |
| FRXESGIEPSVDLETLDER-ESTPKLAK-a12-b5 | pr\|PROT03\|Twa1 | pr\|PROT03\|Twa1 | **intra-protein xl** | 210307_LU | 84 | 215 | 131 | 3092.541 | 619.516 | 5 | 2.5 | 0.33 | 0.64 | **18.74** |
| VEGAETLAYLIEPDVELQR-KVSLGEGRPPVLTASR-a5-b1 | pr\|PROT06\|ARMC8b | pr\|PROT06\|ARMC8b | **intra-protein xl** | 210307_LU | 312 | 388 | 76 | 3792.052 | 759.418 | 5 | 2.6 | 0.31 | 0.58 | **18.56** |
| LLLWAQNELDQK-ESTPKLAK-a8-b5 | pr\|PROT03\|Twa1 | pr\|PROT03\|Twa1 | **intra-protein xl** | 210307_LU | 229 | 215 | 14 | 2324.278 | 582.077 | 4 | 1.7 | 0.16 | 0.42 | **18.45** |
| RKAVESIQAEDESAK-EVFHFSQAEK-a2-b9 | pr\|PROT04\|MAEA | pr\|PROT04\|MAEA | **intra-protein xl** | 210307_LU | 107 | 414 | 307 | 2862.423 | 716.614 | 4 | 3.1 | 0.28 | 0.3 | **18.32** |
| FRMESGIEPSVDLETLDER-ESTPKLAK-a12-b5 | pr\|PROT03\|Twa1 | pr\|PROT03\|Twa1 | **intra-protein xl** | 210307_LU | 84 | 215 | 131 | 3076.546 | 770.144 | 4 | 2.4 | 0.28 | 0.41 | **18.32** |
| VEGAETLAYLIEPDVELQR-YPSSVSAITDIKR-a12-b12 | pr\|PROT06\|ARMC8b | pr\|PROT06\|ARMC8b | **intra-protein xl** | 210307_LU | 319 | 355 | 36 | 3561.874 | 1188.299 | 3 | 4.8 | 0.37 | 0 | **18.31** |
| KGPAAIQK-IQDWYDK-a1-b6 | sp\|P35527\|K1C9_HUMAN | sp\|P35527\|K1C9_HUMAN | **intra-protein xl** | 210307_LU | 192 | 190 | 2 | 1759.928 | 440.99 | 4 | 1.3 | 0.23 | 0.5 | **18.13** |
| ETEAALEFAQTQLAEQGEESR-VKYPKMTDLSK-a3-b5 | pr\|PROT03\|Twa1 | pr\|PROT03\|Twa1 | **intra-protein xl** | 210307_LU | 143 | 239 | 96 | 3626.79 | 726.366 | 5 | 3.4 | 0.33 | 0.53 | **18.02** |
| AVESIQAEDESAK-AAQKNIDR-a10-b4 | pr\|PROT04\|MAEA | pr\|PROT04\|MAEA | **intra-protein xl** | 210307_LU | 117 | 61 | 56 | 2272.138 | 758.387 | 3 | 3.6 | 0.17 | 0 | **18.02** |
| VQEYPTLKVPYETLNK-AVESIQAEDESAK-a8-b8 | pr\|PROT04\|MAEA | pr\|PROT04\|MAEA | **intra-protein xl** | 210307_LU | 46 | 115 | 69 | 3278.662 | 820.673 | 4 | 1.9 | 0.32 | 0.67 | **17.93** |
| EHSSDQPAAASVWK-KAVESIQAEDESAK-a1-b1 | pr\|PROT04\|MAEA | pr\|PROT04\|MAEA | **intra-protein xl** | 210307_LU | 131 | 107 | 24 | 2997.438 | 1000.154 | 3 | 2.1 | 0.17 | 0.59 | **17.87** |
| MSYAEKPDEITKDEWMEK-LLLWAQNELDQK-a12-b8 | pr\|PROT03\|Twa1 | pr\|PROT03\|Twa1 | **intra-protein xl** | 210307_LU | 36 | 229 | 193 | 3680.786 | 921.204 | 4 | 3.3 | 0.15 | 0.41 | **17.85** |
| VPYETLNKR-EVEESLER-a8-b1 | pr\|PROT04\|MAEA | pr\|PROT04\|MAEA | **intra-protein xl** | 210307_LU | 54 | 188 | 134 | 2090.074 | 697.699 | 3 | 4.4 | 0.2 | 0 | **17.77** |
| QKANLIVLGAVPR-MEVTASSR-a2-b2 | pr\|PROT06\|ARMC8b | pr\|PROT06\|ARMC8b | **intra-protein xl** | 210307_LU | 64 | 26 | 38 | 2239.252 | 560.821 | 4 | 2.3 | 0.2 | 0 | **17.55** |
| LLNEVMVEHFFR-ILEALKVR-a8-b6 | pr\|PROT05\|RMND5A | pr\|PROT05\|RMND5A | **intra-protein xl** | 210307_LU | 148 | 189 | 41 | 2455.381 | 614.853 | 4 | 1.5 | 0.24 | 0.56 | **17.48** |
| XESGIEPSVDLETLDER-XTDLSKGVIEEPK-a6-b6 | pr\|PROT03\|Twa1 | pr\|PROT03\|Twa1 | **intra-protein xl** | 210307_LU | 80 | 245 | 165 | 3378.61 | 1127.211 | 3 | 1.2 | 0.17 | 0 | **17.46** |
| ETEAALEFAQTQLAEQGEESR-MTDLSKGVIEEPK-a7-b6 | pr\|PROT03\|Twa1 | pr\|PROT03\|Twa1 | **intra-protein xl** | 210307_LU | 147 | 245 | 98 | 3763.825 | 941.964 | 4 | 3.9 | 0.23 | 0.59 | **17.34** |
| MSYAEKPDEITKDEWMEK-LLLWAQNELDQK-a12-b10 | pr\|PROT03\|Twa1 | pr\|PROT03\|Twa1 | **intra-protein xl** | 210307_LU | 36 | 231 | 195 | 3680.787 | 737.165 | 5 | 3.4 | 0.22 | 0.61 | **17.24** |
| QAAFKLYASLGANDEDIRK-VEGAETLAYLIEPDVELQR-a5-b5 | pr\|PROT06\|ARMC8b | pr\|PROT06\|ARMC8b | **intra-protein xl** | 210307_LU | 373 | 312 | 61 | 4235.189 | 1059.805 | 4 | 3.3 | 0.32 | 0.45 | **17.08** |
| EHSSDQPAAASVWKR-MAVQESAAQLSXTLK-a14-b5 | pr\|PROT04\|MAEA | pr\|PROT04\|MAEA | **intra-protein xl** | 210307_LU | 144 | 28 | 116 | 3272.602 | 819.158 | 4 | 3.1 | 0.15 | 0.42 | **17** |
| DEWMEKLNNLHVQR-XSYAEKPDEITK-a6-b8 | pr\|PROT03\|Twa1 | pr\|PROT03\|Twa1 | **intra-protein xl** | 210307_LU | 42 | 32 | 10 | 3219.539 | 644.916 | 5 | 1.8 | 0.16 | 0.6 | **16.98** |
| AAQKNIDRETSHVTMVVAELEK-AVESIQAEDESAK-a4-b3 | pr\|PROT04\|MAEA | pr\|PROT04\|MAEA | **intra-protein xl** | 210307_LU | 61 | 110 | 49 | 3825.917 | 766.191 | 5 | 3 | 0.33 | 0.82 | **16.96** |
| MSYAEKPDEITKDEWMEK-GVIEEPK-a6-b4 | pr\|PROT03\|Twa1 | pr\|PROT03\|Twa1 | **intra-protein xl** | 210307_LU | 30 | 249 | 219 | 2981.412 | 597.29 | 5 | 2.7 | 0.18 | 0.58 | **16.82** |
| QRETEAALEFAQTQLAEQGEESR-XTDLSKGVIEEPK-a5-b6 | pr\|PROT03\|Twa1 | pr\|PROT03\|Twa1 | **intra-protein xl** | 210307_LU | 143 | 245 | 102 | 4063.98 | 813.804 | 5 | 3.8 | 0.33 | 0.63 | **16.71** |
| GQIQEAIALINSLHPELLDTNR-XTDLSKGVIEEPK-a16-b6 | pr\|PROT03\|Twa1 | pr\|PROT03\|Twa1 | **intra-protein xl** | 210307_LU | 116 | 245 | 129 | 3888.041 | 778.616 | 5 | 2.8 | 0.24 | 0.42 | **16.56** |
| XESGIEPSVDLETLDER-XTDLSKGVIEEPK-a10-b6 | pr\|PROT03\|Twa1 | pr\|PROT03\|Twa1 | **intra-protein xl** | 210307_LU | 84 | 245 | 161 | 3378.616 | 845.662 | 4 | 3.1 | 0.22 | 0.52 | **16.55** |
| LIXNYLVTEGFK-ESTPKLAK-a9-b5 | pr\|PROT03\|Twa1 | pr\|PROT03\|Twa1 | **intra-protein xl** | 210307_LU | 64 | 215 | 151 | 2297.24 | 575.318 | 4 | 2.3 | 0.18 | 0.85 | **16.5** |
| LLLWAQNELDQKK-LIMNYLVTEGFK-a12-b9 | pr\|PROT03\|Twa1 | pr\|PROT03\|Twa1 | **intra-protein xl** | 210307_LU | 233 | 64 | 169 | 3006.635 | 752.667 | 4 | 3.2 | 0.16 | 0.38 | **16.48** |
| TLALLAFDSPEESPFGDLLHTXQR-ESTPKLAK-a11-b5 | pr\|PROT03\|Twa1 | pr\|PROT03\|Twa1 | **intra-protein xl** | 210307_LU | 180 | 215 | 35 | 3557.819 | 712.572 | 5 | 3.2 | 0.26 | 0.39 | **16.32** |
| MSYAEKPDEITK-ESTPKLAK-a5-b5 | pr\|PROT03\|Twa1 | pr\|PROT03\|Twa1 | **intra-protein xl** | 210307_LU | 29 | 215 | 186 | 2265.161 | 567.298 | 4 | 2.1 | 0.15 | 0.86 | **16.2** |

**Description of the column headers**

| | |
|---|---|
| *Id* | Assigned peptides and cross-linking sites within the *peptide* sequences. The longer peptide is designated as (a)lpha, the shorter as (b)eta. |
| *Protein1* | SwissProt/UniProt accession number and identifier of the protein 1 (containing peptide desginated as alpha). |
| *Protein2* | SwissProt/UniProt accession number and identifier of the protein 2 (containing peptide desginated as beta). |
| *XLType* | Intra- or inter-protein, sometimes also ambiguous. |
| *Spectrum* | References the actual MS/MS spectra that were used for the assignment. |
| *AbsPos1* | Position in the *protein* sequence of protein 1. |
| *AbsPos2* | Position in the *protein* sequence of protein 2. |
| *deltaAA* | Distance in the primary sequence (makes only sense for intra-proteins cross-links). |
| *Mr* | Molecular mass calculated from experimental m/z and z (neutral mass). |
| *Mz* | Experimentally observed mass-to-charge ratio of the precursor ion in Da. |
| *z* | Experimentally observed precursor charge. |
| *Error [ppm]* | Deviation between experimental and theoretical mass in ppm. |
| *TIC* | Fraction of the total ion current covered by assigned fragment ions. |
| *delta score* | Relative score for the top hit in relation to the second best hit for this spectrum, calculated as Id-score of 2nd best hit divided by Id-score of best hit. "0" means no second best hit exists. |
| *Id-score* | Identification score as assigned by xQuest. The higher, the better. |
| *X* | in the amino acid sequence denotes an oxidized Met residue |
