## Supplementary material for "The human GID complex engages two independent modules for substrate recruitment": Supp. Table 5

**Supplementary table 5**

### High-Res MS2

**Cross-linked peptide pairs in the 6mer complex identified using DSS as the cross-linking reagent and high-resolution MS/MS detection**

| Id | Protein1 | Protein2 | XLType | Spectrum | AbsPos1 | AbsPos2 | deltaAA | Mr | Mz | z | Error_rel[p | TIC | deltaS | Id-Score |
|---|---|---|---|---|---|---|---|---|---|---|---|---|---|---|
| AAQKNIDR-ILEALKVR-a4-b6 | pr\|PROT04\|MAEA | pr\|PROT05\|RMND5A | inter-protein xl | 210307_LU | 61 | 189 | | 1993.17 | 499.3 | 4 | 0.7 | 0.35 | 0 | **39.27** |
| TKIQAQIDR-MCSKER-a2-b4 | pr\|PROT01\|RanBP9 | pr\|PROT06\|ARMC8b | inter-protein xl | 210307_LU | 372 | 300 | | 2019.028 | 505.765 | 4 | 2 | 0.44 | 0 | **39** |
| LLLWAQNELDQKK-TKIQAQIDR-a12-b2 | pr\|PROT03\|Twa1 | pr\|PROT01\|RanBP9 | inter-protein xl | 210307_LU | 233 | 372 | | 2807.564 | 936.862 | 3 | 3.4 | 0.41 | 0 | **38.68** |
| QDDKVVCPR-ILEALKVR-a4-b6 | pr\|PROT04\|MAEA | pr\|PROT05\|RMND5A | inter-protein xl | 210307_LU | 398 | 189 | | 2194.218 | 732.414 | 3 | 1.9 | 0.21 | 0 | **38.12** |
| KHFSQAEGSQLDEVR-LASDHKDIHSSVSR-a1-b6 | pr\|PROT04\|MAEA | pr\|PROT05\|RMND5A | inter-protein xl | 210307_LU | 246 | 104 | | 3418.696 | 855.682 | 4 | 2.8 | 0.36 | 0 | **37.39** |
| LASDHKDIHSSVSR-AAQKNIDR-a6-b4 | pr\|PROT05\|RMND5A | pr\|PROT04\|MAEA | inter-protein xl | 210307_LU | 104 | 61 | | 2603.348 | 651.845 | 4 | 2.3 | 0.27 | 0 | **37.38** |
| CGYYNTAVKLAR-ILEALKVR-a9-b6 | pr\|PROT04\|MAEA | pr\|PROT05\|RMND5A | inter-protein xl | 210307_LU | 167 | 189 | | 2493.387 | 624.354 | 4 | 3.6 | 0.23 | 0 | **36.67** |
| TKIQAQIDR-XCSKER-a2-b4 | pr\|PROT01\|RanBP9 | pr\|PROT06\|ARMC8b | inter-protein xl | 210307_LU | 372 | 300 | | 2035.024 | 679.349 | 3 | 2.5 | 0.41 | 0 | **36.66** |
| KVSLGEGRPPVLTASR-MTDLSKGVIEEPK-a1-b6 | pr\|PROT06\|ARMC8b | pr\|PROT03\|Twa1 | inter-protein xl | 210307_LU | 388 | 245 | | 3249.774 | 650.963 | 5 | 3.1 | 0.4 | 0 | **36.26** |
| AAQKNIDR-VGKAIDK-a4-b3 | pr\|PROT04\|MAEA | pr\|PROT05\|RMND5A | inter-protein xl | 210307_LU | 61 | 115 | | 1782.003 | 446.509 | 4 | 1.8 | 0.37 | 0 | **36.14** |
| EHSSDQPAAASVWKR-ILEALKVR-a14-b6 | pr\|PROT04\|MAEA | pr\|PROT05\|RMND5A | inter-protein xl | 210307_LU | 144 | 189 | | 2746.483 | 687.629 | 4 | 2.3 | 0.23 | 0 | **35.74** |
| TKIQAQIDRFPIGDR-XCSKER-a2-b4 | pr\|PROT01\|RanBP9 | pr\|PROT06\|ARMC8b | inter-protein xl | 210307_LU | 372 | 300 | | 2720.38 | 681.103 | 4 | 2.4 | 0.25 | 0 | **35.7** |
| LASDHKDIHSSVSR-QDDKVVCPR-a6-b4 | pr\|PROT05\|RMND5A | pr\|PROT04\|MAEA | inter-protein xl | 210307_LU | 104 | 398 | | 2804.394 | 561.887 | 5 | 2.3 | 0.25 | 0 | **35.64** |
| LASDHKDIHSSVSR-VPYETLNKR-a6-b8 | pr\|PROT05\|RMND5A | pr\|PROT04\|MAEA | inter-protein xl | 210307_LU | 104 | 54 | | 2807.463 | 468.918 | 6 | 2.2 | 0.18 | 0 | **35.4** |
| TKIQAQIDRFPIGDR-MCSKER-a2-b4 | pr\|PROT01\|RanBP9 | pr\|PROT06\|ARMC8b | inter-protein xl | 210307_LU | 372 | 300 | | 2704.386 | 541.885 | 5 | 2.5 | 0.32 | 0 | **35.35** |
| AAQKNIDRETSHVTMVVAELEK-LASDHKDIHSSVSR-a4-b6 | pr\|PROT04\|MAEA | pr\|PROT05\|RMND5A | inter-protein xl | 210307_LU | 61 | 104 | | 4157.136 | 1040.292 | 4 | 4.4 | 0.28 | 0 | **35.32** |
| ETATCLAWCHDNKSR-LASDHKDIHSSVSR-a13-b6 | pr\|PROT04\|MAEA | pr\|PROT05\|RMND5A | inter-protein xl | 210307_LU | 209 | 104 | | 3536.663 | 885.174 | 4 | 2.9 | 0.26 | 0 | **35.14** |
| KVSLGEGRPPVLTASR-EAAEKFR-a1-b5 | pr\|PROT06\|ARMC8b | pr\|PROT03\|Twa1 | inter-protein xl | 210307_LU | 388 | 72 | | 2653.459 | 531.7 | 5 | 1.3 | 0.22 | 0 | **35.04** |
| AAQKNIDRETSHVTXVVAELEK-LASDHKDIHSSVSR-a4-b6 | pr\|PROT04\|MAEA | pr\|PROT05\|RMND5A | inter-protein xl | 210307_LU | 61 | 104 | | 4173.124 | 835.633 | 5 | 2.7 | 0.22 | 0 | **34.46** |
| CGYYNTAVKLAR-EAAEKFR-a9-b5 | pr\|PROT04\|MAEA | pr\|PROT03\|Twa1 | inter-protein xl | 210307_LU | 167 | 72 | | 2402.213 | 601.561 | 4 | 3.1 | 0.1 | 0 | **33.07** |
| EGEWQTMIQKMVSSYLVHHGYCATAEAFAR-MTDLSKGVIEEPK-a10-b6 | pr\|PROT01\|RanBP9 | pr\|PROT03\|Twa1 | inter-protein xl | 210307_LU | 395 | 245 | | 5083.441 | 1017.696 | 5 | 3.5 | 0.23 | 0 | **31.67** |
| QGMLDVAEELCQESGLSVDPSQKEPFVELNR-SLNKLAQPLPXAHCANSR-a23-b4 | pr\|PROT05\|RMND5A | pr\|PROT04\|MAEA | inter-protein xl | 210307_LU | 175 | 346 | | 5664.747 | 1133.957 | 5 | 2.7 | 0.18 | 0 | **29.56** |
| YPSSVSAITDIKR-ESTPKLAK-a12-b5 | pr\|PROT06\|ARMC8b | pr\|PROT03\|Twa1 | inter-protein xl | 210307_LU | 355 | 215 | | 2446.338 | 612.592 | 4 | 2.5 | 0.11 | 0 | **28.94** |
| LASDHKDIHSSVSR-VGKAIDK-a6-b3 | pr\|PROT05\|RMND5A | pr\|PROT05\|RMND5A | intra-protein xl | 210307_LU | 104 | 115 | 11 | 2418.294 | 807.106 | 3 | 2.9 | 0.42 | 0 | **40.55** |
| QDDKVVCPR-AAQKNIDR-a4-b4 | pr\|PROT04\|MAEA | pr\|PROT04\|MAEA | intra-protein xl | 210307_LU | 398 | 61 | 337 | 2168.104 | 543.034 | 4 | 1.7 | 0.32 | 0 | **39.79** |
| KHFSQAEGSQLDEVR-AAQKNIDR-a1-b4 | pr\|PROT04\|MAEA | pr\|PROT04\|MAEA | intra-protein xl | 210307_LU | 246 | 61 | 185 | 2782.408 | 696.61 | 4 | 3.1 | 0.42 | 0 | **39.61** |
| KAVESIQAEDESAK-AAQKNIDR-a1-b4 | pr\|PROT04\|MAEA | pr\|PROT04\|MAEA | intra-protein xl | 210307_LU | 107 | 61 | 46 | 2556.31 | 853.111 | 3 | 2.7 | 0.57 | 0 | **38.89** |
| ILEALKVR-VGKAIDK-a6-b3 | pr\|PROT05\|RMND5A | pr\|PROT05\|RMND5A | intra-protein xl | 210307_LU | 189 | 115 | 74 | 1808.117 | 453.037 | 4 | 2.1 | 0.24 | 0 | **38.64** |
| TPKDAASVR-IVSKGR-a3-b4 | pr\|PROT01\|RanBP9 | pr\|PROT01\|RanBP9 | intra-protein xl | 210307_LU | 221 | 248 | 27 | 1739.992 | 436.006 | 4 | 1.6 | 0.43 | 0 | **38.22** |
| AKAEWEGK-SKLLDK-a2-b2 | pr\|PROT02\|WDR26 | pr\|PROT02\|WDR26 | intra-protein xl | 210307_LU | 172 | 185 | 13 | 1757.961 | 586.995 | 3 | 2.8 | 0.47 | 0 | **38.2** |
| KAVESIQAEDESAK-VPYETLNKR-a1-b8 | pr\|PROT04\|MAEA | pr\|PROT04\|MAEA | intra-protein xl | 210307_LU | 107 | 54 | 53 | 2760.426 | 921.15 | 3 | 3 | 0.45 | 0 | **37.98** |
| RETATCLAWCHDNKSR-KHFSQAEGSQLDEVR-a14-b1 | pr\|PROT04\|MAEA | pr\|PROT04\|MAEA | intra-protein xl | 210307_LU | 209 | 246 | 37 | 3871.827 | 775.373 | 5 | 3.8 | 0.33 | 0 | **37.79** |
| LKCPYCPMEQSPGDAK-DALNKMFNGSK-a2-b5 | pr\|PROT05\|RMND5A | pr\|PROT05\|RMND5A | intra-protein xl | 210307_LU | 397 | 389 | 8 | 3241.501 | 1081.508 | 3 | 2.9 | 0.48 | 0 | **37.61** |
| TKEVFHFSQAEK-QDDKVVCPR-a2-b4 | pr\|PROT04\|MAEA | pr\|PROT04\|MAEA | intra-protein xl | 210307_LU | 405 | 398 | 7 | 2703.342 | 902.122 | 3 | 3.5 | 0.21 | 0 | **37.41** |
| VQCLWCLSDGKTVLASDTHQR-VYIWHKR-a11-b6 | pr\|PROT02\|WDR26 | pr\|PROT02\|WDR26 | intra-protein xl | 210307_LU | 402 | 504 | 102 | 3611.816 | 723.371 | 5 | 0.8 | 0.17 | 0 | **37.13** |
| CELTPLKYNTER-MKFLLLQQK-a7-b2 | pr\|PROT02\|WDR26 | pr\|PROT02\|WDR26 | intra-protein xl | 210307_LU | 148 | 116 | 32 | 2808.502 | 703.133 | 4 | 3.6 | 0.33 | 0 | **37.04** |
| KHFSQAEGSQLDEVR-QDDKVVCPR-a1-b4 | pr\|PROT04\|MAEA | pr\|PROT04\|MAEA | intra-protein xl | 210307_LU | 246 | 398 | 152 | 2983.456 | 597.699 | 5 | 3.3 | 0.33 | 0 | **36.95** |
| STDQTVLEELASIKNR-IQKLVLAGR-a14-b3 | pr\|PROT01\|RanBP9 | pr\|PROT01\|RanBP9 | intra-protein xl | 210307_LU | 429 | 436 | 7 | 2937.655 | 588.539 | 5 | 1.6 | 0.35 | 0 | **36.9** |
| RETATCLAWCHDNKSR-AAQKNIDR-a14-b4 | pr\|PROT04\|MAEA | pr\|PROT04\|MAEA | intra-protein xl | 210307_LU | 209 | 61 | 148 | 3056.477 | 765.127 | 4 | 3.2 | 0.36 | 0 | **36.81** |
| VPYETLNKR-AAQKNIDR-a8-b4 | pr\|PROT04\|MAEA | pr\|PROT04\|MAEA | intra-protein xl | 210307_LU | 54 | 61 | 7 | 2171.174 | 724.732 | 3 | 1.9 | 0.43 | 0 | **36.76** |
| VLHKFSGYGQLCER-DTVQKLASDHK-a4-b5 | pr\|PROT05\|RMND5A | pr\|PROT05\|RMND5A | intra-protein xl | 210307_LU | 41 | 98 | 57 | 3071.56 | 1024.861 | 3 | 3.4 | 0.32 | 0 | **36.65** |
| LKCPYCPMEQSPGDAK-DALNKXFNGSK-a2-b5 | pr\|PROT05\|RMND5A | pr\|PROT05\|RMND5A | intra-protein xl | 210307_LU | 397 | 389 | 8 | 3257.491 | 815.381 | 4 | 1.4 | 0.35 | 0.89 | **36.65** |
| RKAVESIQAEDESAK-AAQKNIDR-a2-b4 | pr\|PROT04\|MAEA | pr\|PROT04\|MAEA | intra-protein xl | 210307_LU | 107 | 61 | 46 | 2712.414 | 679.111 | 4 | 3.6 | 0.31 | 0 | **36.58** |
| CELTPLKYNTER-AKAEWEGK-a7-b2 | pr\|PROT02\|WDR26 | pr\|PROT02\|WDR26 | intra-protein xl | 210307_LU | 148 | 172 | 24 | 2578.277 | 860.434 | 3 | 1.4 | 0.42 | 0 | **36.52** |
| YLEYLEDGKVLEALQVLR-AEWEGKGTASR-a9-b6 | pr\|PROT02\|WDR26 | pr\|PROT02\|WDR26 | intra-protein xl | 210307_LU | 132 | 178 | 46 | 3478.808 | 870.71 | 4 | 2.8 | 0.3 | 0 | **36.45** |
| MSYAEKPDEITKDEWMEK-MTDLSKGVIEEPK-a12-b6 | pr\|PROT03\|Twa1 | pr\|PROT03\|Twa1 | intra-protein xl | 210307_LU | 36 | 245 | 209 | 3812.821 | 763.572 | 5 | 3.2 | 0.23 | 0 | **36.38** |
| NIVQEDHPIMSFTISKNGR-VYIWHKR-a16-b6 | pr\|PROT02\|WDR26 | pr\|PROT02\|WDR26 | intra-protein xl | 210307_LU | 440 | 504 | 64 | 3323.734 | 831.941 | 4 | 3 | 0.17 | 0 | **36.31** |
| ELEKVLHK-VGKAIDK-a4-b3 | pr\|PROT05\|RMND5A | pr\|PROT05\|RMND5A | intra-protein xl | 210307_LU | 37 | 115 | 78 | 1862.091 | 466.531 | 4 | 2 | 0.2 | 0 | **35.96** |
| YPSSVSAITDIKR-LDHDLKHAHELR-a12-b6 | pr\|PROT06\|ARMC8b | pr\|PROT06\|ARMC8b | intra-protein xl | 210307_LU | 355 | 362 | 7 | 3056.611 | 612.33 | 5 | 2.1 | 0.21 | 0 | **35.92** |
| KHFSQAEGSQLDEVR-KAVESIQAEDESAK-a1-b1 | pr\|PROT04\|MAEA | pr\|PROT04\|MAEA | intra-protein xl | 210307_LU | 246 | 107 | 139 | 3371.659 | 675.34 | 5 | 3.3 | 0.36 | 0 | **35.9** |
| LLDKLQTYLPPSVMLPPR-CELTPLKYNTER-a4-b7 | pr\|PROT02\|WDR26 | pr\|PROT02\|WDR26 | intra-protein xl | 210307_LU | 189 | 148 | 41 | 3740.997 | 936.257 | 4 | 2.4 | 0.27 | 0 | **35.9** |
| CELTPLKYNTER-XKFLLLQQK-a7-b2 | pr\|PROT02\|WDR26 | pr\|PROT02\|WDR26 | intra-protein xl | 210307_LU | 148 | 116 | 32 | 2824.494 | 707.131 | 4 | 2.7 | 0.32 | 0 | **35.9** |
| SLNKLAQPLPMAHCANSR-EHSSDQPAAASVWKR-a4-b14 | pr\|PROT04\|MAEA | pr\|PROT04\|MAEA | intra-protein xl | 210307_LU | 346 | 144 | 202 | 3812.896 | 763.587 | 5 | 3.3 | 0.26 | 0 | **35.84** |
| ETATCLAWCHDNKSR-KHFSQAEGSQLDEVR-a13-b1 | pr\|PROT04\|MAEA | pr\|PROT04\|MAEA | intra-protein xl | 210307_LU | 209 | 246 | 37 | 3715.719 | 929.938 | 4 | 2.1 | 0.25 | 0 | **35.83** |
| XTDLSKGVIEEPK-ESTPKLAK-a6-b5 | pr\|PROT03\|Twa1 | pr\|PROT03\|Twa1 | intra-protein xl | 210307_LU | 245 | 215 | 30 | 2472.311 | 619.085 | 4 | 2.9 | 0.15 | 0 | **35.82** |
| KVSLGEGRPPVLTASR-XCSKER-a1-b4 | pr\|PROT06\|ARMC8b | pr\|PROT06\|ARMC8b | intra-protein xl | 210307_LU | 388 | 300 | 88 | 2629.375 | 658.352 | 4 | 2.7 | 0.29 | 0 | **35.73** |
| KVSLGEGRPPVLTASR-YPSSVSAITDIKR-a1-b12 | pr\|PROT06\|ARMC8b | pr\|PROT06\|ARMC8b | intra-protein xl | 210307_LU | 388 | 355 | 33 | 3239.798 | 648.967 | 5 | 3.1 | 0.34 | 0 | **35.7** |
| KVSLGEGRPPVLTASR-MCSKER-a1-b4 | pr\|PROT06\|ARMC8b | pr\|PROT06\|ARMC8b | intra-protein xl | 210307_LU | 388 | 300 | 88 | 2613.382 | 654.353 | 4 | 3.5 | 0.34 | 0 | **35.69** |
| LFDPDPQKVLQGVIDMK-NAVIGNNKQK-a8-b8 | pr\|PROT06\|ARMC8b | pr\|PROT06\|ARMC8b | intra-protein xl | 210307_LU | 45 | 62 | 17 | 3164.698 | 792.182 | 4 | 2.5 | 0.19 | 0 | **35.6** |
| MTDLSKGVIEEPK-ESTPKLAK-a6-b5 | pr\|PROT03\|Twa1 | pr\|PROT03\|Twa1 | intra-protein xl | 210307_LU | 245 | 215 | 30 | 2456.314 | 615.086 | 4 | 2.4 | 0.18 | 0 | **35.56** |
| LIMNYLVTEGFKEAAEK-MTDLSKGVIEEPK-a12-b6 | pr\|PROT03\|Twa1 | pr\|PROT03\|Twa1 | intra-protein xl | 210307_LU | 67 | 245 | 178 | 3538.83 | 885.715 | 4 | 3.1 | 0.17 | 0 | **35.53** |
| TPQCYKEDGSSKSPDCPVCSR-SLNKLAQPLPMAHCANSR-a12-b4 | pr\|PROT04\|MAEA | pr\|PROT04\|MAEA | intra-protein xl | 210307_LU | 333 | 346 | 13 | 4602.132 | 1151.541 | 4 | 3 | 0.32 | 0 | **35.51** |
| AAQKNIDRETSHVTMVVAELEK-KAVESIQAEDESAK-a4-b1 | pr\|PROT04\|MAEA | pr\|PROT04\|MAEA | intra-protein xl | 210307_LU | 61 | 107 | 46 | 4110.09 | 1028.53 | 4 | 2.6 | 0.26 | 0 | **35.38** |
| KHFSQAEGSQLDEVR-RKAVESIQAEDESAK-a1-b2 | pr\|PROT04\|MAEA | pr\|PROT04\|MAEA | intra-protein xl | 210307_LU | 246 | 107 | 139 | 3527.755 | 706.559 | 5 | 1.6 | 0.29 | 0 | **35.3** |
| YPSSVSAITDIKR-MCSKER-a12-b4 | pr\|PROT06\|ARMC8b | pr\|PROT06\|ARMC8b | intra-protein xl | 210307_LU | 355 | 300 | 55 | 2383.194 | 596.806 | 4 | 2.5 | 0.29 | 0 | **35.09** |
| LKCPYCPXEQSPGDAK-DALNKMFNGSK-a2-b5 | pr\|PROT05\|RMND5A | pr\|PROT05\|RMND5A | intra-protein xl | 210307_LU | 397 | 389 | 8 | 3257.499 | 815.383 | 4 | 4 | 0.27 | 0 | **35.06** |
| AAQKNIDRETSHVTMVVAELEK-VPYETLNKR-a4-b8 | pr\|PROT04\|MAEA | pr\|PROT04\|MAEA | intra-protein xl | 210307_LU | 61 | 54 | 7 | 3724.96 | 746 | 5 | 3.8 | 0.22 | 0 | **34.92** |
| EAAEKFR-EAAEKFR-a5-b5 | pr\|PROT03\|Twa1 | pr\|PROT03\|Twa1 | intra-protein xl | 210307_LU | 72 | 72 | 0 | 1836.94 | 460.243 | 4 | 1.8 | 0.16 | 0 | **34.89** |
| AAQKNIDRETSHVTMVVAELEK-ETATCLAWCHDNKSR-a4-b13 | pr\|PROT04\|MAEA | pr\|PROT04\|MAEA | intra-protein xl | 210307_LU | 61 | 209 | 148 | 4454.157 | 891.839 | 5 | 3.4 | 0.18 | 0 | **34.42** |
| QAAFKLYASLGANDEDIR-LDHDLKHAHELR-a5-b6 | pr\|PROT06\|ARMC8b | pr\|PROT06\|ARMC8b | intra-protein xl | 210307_LU | 373 | 362 | 11 | 3601.843 | 901.469 | 4 | 4.2 | 0.26 | 0 | **34.38** |
| MSYAEKPDEITKDEWXEK-XTDLSKGVIEEPK-a12-b6 | pr\|PROT03\|Twa1 | pr\|PROT03\|Twa1 | intra-protein xl | 210307_LU | 36 | 245 | 209 | 3844.811 | 769.97 | 5 | 3.2 | 0.2 | 0 | **34.3** |
| EHSSDQPAAASVWKR-QDDKVVCPR-a14-b4 | pr\|PROT04\|MAEA | pr\|PROT04\|MAEA | intra-protein xl | 210307_LU | 144 | 398 | 254 | 2921.42 | 731.363 | 4 | 3.7 | 0.12 | 0 | **34.1** |
| QAAFKLYASLGANDEDIRK-KVSLGEGRPPVLTASR-a5-b1 | pr\|PROT06\|ARMC8b | pr\|PROT06\|ARMC8b | intra-protein xl | 210307_LU | 373 | 388 | 15 | 3913.119 | 783.632 | 5 | 3.4 | 0.3 | 0 | **34.07** |
| DEWXEKLNNLHVQR-XSYAEKPDEITK-a6-b6 | pr\|PROT03\|Twa1 | pr\|PROT03\|Twa1 | intra-protein xl | 210307_LU | 42 | 30 | 12 | 3391.617 | 848.912 | 4 | 3.2 | 0.13 | 0 | **34.02** |
| ETATCLAWCHDNKSR-AAQKNIDR-a13-b4 | pr\|PROT04\|MAEA | pr\|PROT04\|MAEA | intra-protein xl | 210307_LU | 209 | 61 | 148 | 2900.376 | 726.102 | 4 | 3.5 | 0.17 | 0 | **33.39** |
| LIMNYLVTEGFKEAAEKFR-EAAEKFR-a17-b5 | pr\|PROT03\|Twa1 | pr\|PROT03\|Twa1 | intra-protein xl | 210307_LU | 72 | 72 | 0 | 3245.686 | 650.145 | 5 | 2.1 | 0.1 | 0 | **33.33** |
| SLNKLAQPLPMAHCANSR-EDGSSKSPDCPVCSR-a4-b6 | pr\|PROT04\|MAEA | pr\|PROT04\|MAEA | intra-protein xl | 210307_LU | 346 | 333 | 13 | 3824.782 | 957.203 | 4 | 3 | 0.45 | 0 | **32.71** |
| TPQCYKEDGSSKSPDCPVCSR-SLNKLAQPLPXAHCANSR-a12-b4 | pr\|PROT04\|MAEA | pr\|PROT04\|MAEA | intra-protein xl | 210307_LU | 333 | 346 | 13 | 4618.128 | 770.696 | 6 | 3.3 | 0.28 | 0 | **32.66** |
| SLNKLAQPLPXAHCANSR-EDGSSKSPDCPVCSR-a4-b6 | pr\|PROT04\|MAEA | pr\|PROT04\|MAEA | intra-protein xl | 210307_LU | 346 | 333 | 13 | 3840.774 | 961.201 | 4 | 2.3 | 0.52 | 0 | **32.47** |
| MSYAEKPDEITKDEWMEK-LLLWAQNELDQKK-a6-b12 | pr\|PROT03\|Twa1 | pr\|PROT03\|Twa1 | intra-protein xl | 210307_LU | 30 | 233 | 203 | 3964.959 | 992.248 | 4 | 2.9 | 0.12 | 0 | **31.64** |
| SLNKLAQPLPXAHCANSR-QDDKVVCPR-a4-b4 | pr\|PROT04\|MAEA | pr\|PROT04\|MAEA | intra-protein xl | 210307_LU | 346 | 398 | 52 | 3276.618 | 820.162 | 4 | 0.5 | 0.25 | 0 | **31.21** |
| VQCLWCLSDGKTVLASDTHQR-MSQSHEDSLTSVAWNPDGKR-a11-b19 | pr\|PROT02\|WDR26 | pr\|PROT02\|WDR26 | intra-protein xl | 210307_LU | 402 | 363 | 39 | 4855.292 | 972.066 | 5 | 3.5 | 0.12 | 0 | **30.78** |
| MSYAEKPDEITKDEWMEK-XTDLSKGVIEEPK-a12-b6 | pr\|PROT03\|Twa1 | pr\|PROT03\|Twa1 | intra-protein xl | 210307_LU | 36 | 245 | 209 | 3828.818 | 958.212 | 4 | 3.9 | 0.2 | 0 | **29.73** |

### Low-Res MS2

**Cross-linked peptide pairs in the 6mer complex identified using DSS as the cross-linking reagent and low-resolution MS/MS detection**

| Id | Protein1 | Protein2 | XLType | Spectrum | AbsPos1 | AbsPos2 | deltaAA | Mr | Mz | z | Error_rel[p | TIC | deltaS | Id-Score |
|---|---|---|---|---|---|---|---|---|---|---|---|---|---|---|
| LLLWAQNELDQKK-TKIQAQIDR-a12-b2 | pr\|PROT03\|Twa1 | pr\|PROT01\|RanBP9 | inter-protein xl | 210307_LU | 233 | 372 | | 2807.562 | 936.862 | 3 | 2.7 | 0.33 | 0 | **34.05** |
| CGYYNTAVKLAR-ILEALKVR-a9-b6 | pr\|PROT04\|MAEA | pr\|PROT05\|RMND5A | inter-protein xl | 210307_LU | 167 | 189 | | 2493.383 | 832.135 | 3 | 1.9 | 0.62 | 0 | **33.95** |
| IQKLVLAGR-EAAEKFR-a3-b5 | pr\|PROT01\|RanBP9 | pr\|PROT03\|Twa1 | inter-protein xl | 210307_LU | 436 | 72 | | 1984.154 | 662.392 | 3 | 3.5 | 0.43 | 0.73 | **33.47** |
| LASDHKDIHSSVSR-AAQKNIDR-a6-b4 | pr\|PROT05\|RMND5A | pr\|PROT04\|MAEA | inter-protein xl | 210307_LU | 104 | 61 | | 2603.349 | 868.791 | 3 | 2.8 | 0.46 | 0 | **32.9** |
| MTDLSKGVIEEPK-MCSKER-a6-b4 | pr\|PROT03\|Twa1 | pr\|PROT06\|ARMC8b | inter-protein xl | 210307_LU | 245 | 300 | | 2393.173 | 798.732 | 3 | 3.5 | 0.41 | 0 | **32.89** |
| KHFSQAEGSQLDEVR-LASDHKDIHSSVSR-a1-b6 | pr\|PROT04\|MAEA | pr\|PROT05\|RMND5A | inter-protein xl | 210307_LU | 246 | 104 | | 3418.7 | 684.748 | 5 | 3.9 | 0.46 | 0 | **32.58** |
| TKIQAQIDR-MCSKER-a2-b4 | pr\|PROT01\|RanBP9 | pr\|PROT06\|ARMC8b | inter-protein xl | 210307_LU | 372 | 300 | | 2019.03 | 674.018 | 3 | 2.8 | 0.51 | 0.34 | **32.19** |
| TKIQAQIDRFPIGDR-MCSKER-a2-b4 | pr\|PROT01\|RanBP9 | pr\|PROT06\|ARMC8b | inter-protein xl | 210307_LU | 372 | 300 | | 2704.389 | 677.105 | 4 | 3.8 | 0.41 | 0.69 | **32.13** |
| QDDKVVCPR-ILEALKVR-a4-b6 | pr\|PROT04\|MAEA | pr\|PROT05\|RMND5A | inter-protein xl | 210307_LU | 398 | 189 | | 2194.223 | 732.415 | 3 | 3.8 | 0.32 | 0 | **31.98** |
| ETATCLAWCHDNKSR-VGKAIDK-a13-b3 | pr\|PROT04\|MAEA | pr\|PROT05\|RMND5A | inter-protein xl | 210307_LU | 209 | 115 | | 2715.318 | 906.114 | 3 | 2.7 | 0.28 | 0 | **31.95** |
| EHSSDQPAAASVWKR-ILEALKVR-a14-b6 | pr\|PROT04\|MAEA | pr\|PROT05\|RMND5A | inter-protein xl | 210307_LU | 144 | 189 | | 2746.483 | 687.628 | 4 | 2.2 | 0.3 | 0 | **31.85** |
| ETATCLAWCHDNKSR-LASDHKDIHSSVSR-a13-b6 | pr\|PROT04\|MAEA | pr\|PROT05\|RMND5A | inter-protein xl | 210307_LU | 209 | 104 | | 3536.662 | 885.173 | 4 | 2.7 | 0.32 | 0 | **29.17** |
| NAVIGNNKQK-AAQKNIDR-a8-b4 | pr\|PROT06\|ARMC8b | pr\|PROT04\|MAEA | inter-protein xl | 210307_LU | 62 | 61 | | 2137.162 | 713.395 | 3 | 1 | 0.46 | 0 | **29.15** |
| MTDLSKGVIEEPK-NAVIGNNKQK-a6-b8 | pr\|PROT03\|Twa1 | pr\|PROT06\|ARMC8b | inter-protein xl | 210307_LU | 245 | 62 | | 2668.414 | 890.479 | 3 | 1.5 | 0.3 | 0.78 | **27.96** |
| MTDLSKGVIEEPK-LDHDLKHAHELR-a6-b6 | pr\|PROT03\|Twa1 | pr\|PROT06\|ARMC8b | inter-protein xl | 210307_LU | 245 | 362 | | 3066.593 | 767.656 | 4 | 4 | 0.31 | 0 | **27.89** |
| KHFSQAEGSQLDEVR-MTDLSKGVIEEPK-a1-b6 | pr\|PROT04\|MAEA | pr\|PROT03\|Twa1 | inter-protein xl | 210307_LU | 246 | 245 | | 3313.657 | 829.422 | 4 | 2.2 | 0.29 | 0.59 | **27.35** |
| LASDHKDIHSSVSR-NAVIGNNKQK-a6-b8 | pr\|PROT05\|RMND5A | pr\|PROT06\|ARMC8b | inter-protein xl | 210307_LU | 104 | 62 | | 2773.453 | 555.698 | 5 | 2.1 | 0.33 | 0 | **26.36** |
| KVSLGEGRPPVLTASR-KHFSQAEGSQLDEVR-a1-b1 | pr\|PROT06\|ARMC8b | pr\|PROT04\|MAEA | inter-protein xl | 210307_LU | 388 | 246 | | 3533.868 | 707.781 | 5 | 2.5 | 0.28 | 0 | **26.09** |
| AAQKNIDRETSHVTXVVAELEK-LASDHKDIHSSVSR-a4-b6 | pr\|PROT04\|MAEA | pr\|PROT05\|RMND5A | inter-protein xl | 210307_LU | 61 | 104 | | 4173.129 | 835.634 | 5 | 4 | 0.25 | 0.52 | **26.06** |
| LASDHKDIHSSVSR-VPYETLNKR-a6-b8 | pr\|PROT05\|RMND5A | pr\|PROT04\|MAEA | inter-protein xl | 210307_LU | 104 | 54 | | 2807.464 | 562.501 | 5 | 2.6 | 0.16 | 0 | **25.81** |
| LASDHKDIHSSVSR-QDDKVVCPR-a6-b4 | pr\|PROT05\|RMND5A | pr\|PROT04\|MAEA | inter-protein xl | 210307_LU | 104 | 398 | | 2804.398 | 702.107 | 4 | 3.7 | 0.26 | 0 | **25.33** |
| VPYETLNKR-VGKAIDK-a8-b3 | pr\|PROT04\|MAEA | pr\|PROT05\|RMND5A | inter-protein xl | 210307_LU | 54 | 115 | | 1986.118 | 497.537 | 4 | 1.6 | 0.37 | 0 | **25.26** |
| TKIQAQIDRFPIGDR-LLLWAQNELDQKK-a2-b12 | pr\|PROT01\|RanBP9 | pr\|PROT03\|Twa1 | inter-protein xl | 210307_LU | 372 | 233 | | 3492.92 | 699.592 | 5 | 3.1 | 0.13 | 0 | **24.31** |
| AAQKNIDRETSHVTMVVAELEK-LASDHKDIHSSVSR-a4-b6 | pr\|PROT04\|MAEA | pr\|PROT05\|RMND5A | inter-protein xl | 210307_LU | 61 | 104 | | 4157.135 | 693.864 | 6 | 4.1 | 0.28 | 0.36 | **24.1** |
| AAQKNIDR-ILEALKVR-a4-b6 | pr\|PROT04\|MAEA | pr\|PROT05\|RMND5A | inter-protein xl | 210307_LU | 61 | 189 | | 1993.169 | 499.3 | 4 | 0.5 | 0.13 | 0 | **22.64** |
| KVSLGEGRPPVLTASR-MTDLSKGVIEEPK-a1-b6 | pr\|PROT06\|ARMC8b | pr\|PROT03\|Twa1 | inter-protein xl | 210307_LU | 388 | 245 | | 3249.775 | 813.452 | 4 | 3.4 | 0.35 | 0 | **21.95** |
| NAVIGNNKQK-AKAEWEGK-a8-b2 | pr\|PROT06\|ARMC8b | pr\|PROT02\|WDR26 | inter-protein xl | 210307_LU | 62 | 172 | | 2140.135 | 714.386 | 3 | 3.2 | 0.34 | 0.71 | **21.41** |
| TKIQAQIDRFPIGDR-YPSSVSAITDIKR-a2-b12 | pr\|PROT01\|RanBP9 | pr\|PROT06\|ARMC8b | inter-protein xl | 210307_LU | 372 | 355 | | 3330.8 | 667.168 | 5 | 2 | 0.17 | 0 | **21.1** |
| XAVQESAAQLSMTLKVQEYPTLK-DVDGAYXTKVDLQAK-a15-b9 | pr\|PROT04\|MAEA | sp\|P04264\|K2C1_HUMA | inter-protein xl | 210307_LU | 38 | 298 | | 4388.201 | 1098.058 | 4 | 3.9 | 0.17 | 0.81 | **20.83** |
| LDHDLKHAHELR-ESTPKLAK-a6-b5 | pr\|PROT06\|ARMC8b | pr\|PROT03\|Twa1 | inter-protein xl | 210307_LU | 362 | 215 | | 2493.337 | 624.342 | 4 | 1.2 | 0.1 | 0 | **20.25** |
| NAVIGNNKQK-QDDKVVCPR-a8-b4 | pr\|PROT06\|ARMC8b | pr\|PROT04\|MAEA | inter-protein xl | 210307_LU | 62 | 398 | | 2338.213 | 585.561 | 4 | 3 | 0.21 | 0 | **20.05** |
| SLNKLAQPLPMAHCANSR-ILEALKVR-a4-b6 | pr\|PROT04\|MAEA | pr\|PROT05\|RMND5A | inter-protein xl | 210307_LU | 346 | 189 | | 3085.697 | 618.147 | 5 | 2.6 | 0.16 | 0.5 | **19.61** |
| KVSLGEGRPPVLTASR-ESTPKLAK-a1-b5 | pr\|PROT06\|ARMC8b | pr\|PROT03\|Twa1 | inter-protein xl | 210307_LU | 388 | 215 | | 2676.525 | 670.139 | 4 | 2.7 | 0.21 | 0.68 | **19.11** |
| RETATCLAWCHDNKSR-LASDHKDIHSSVSR-a14-b6 | pr\|PROT04\|MAEA | pr\|PROT05\|RMND5A | inter-protein xl | 210307_LU | 209 | 104 | | 3692.765 | 924.199 | 4 | 3 | 0.2 | 0 | **18.63** |
| KVSLGEGRPPVLTASR-QDDKVVCPR-a1-b4 | pr\|PROT06\|ARMC8b | pr\|PROT04\|MAEA | inter-protein xl | 210307_LU | 388 | 398 | | 2919.567 | 584.921 | 5 | 2.2 | 0.19 | 0 | **17.78** |
| VLQGVIDMKNAVIGNNK-SWSPKDK-a9-b5 | pr\|PROT06\|ARMC8b | pr\|PROT01\|RanBP9 | inter-protein xl | 210307_LU | 54 | 196 | | 2796.496 | 560.307 | 5 | 4 | 0.15 | 0.86 | **17.68** |
| QQTTDNNPPMKLVCGHIISR-EHSSDQPAAASVWKR-a11-b14 | pr\|PROT05\|RMND5A | pr\|PROT04\|MAEA | inter-protein xl | 210307_LU | 375 | 144 | | 4114.027 | 823.813 | 5 | 3.9 | 0.14 | 0 | **17.43** |
| YPSSVSAITDIKR-LDHDLKHAHELR-a12-b6 | pr\|PROT06\|ARMC8b | pr\|PROT06\|ARMC8b | intra-protein xl | 210307_LU | 355 | 362 | 7 | 3056.613 | 612.331 | 5 | 3 | 0.29 | 0.32 | **34.04** |
| VPYETLNKR-AAQKNIDR-a8-b4 | pr\|PROT04\|MAEA | pr\|PROT04\|MAEA | intra-protein xl | 210307_LU | 54 | 61 | 7 | 2171.171 | 724.732 | 3 | 0.7 | 0.67 | 0 | **33.9** |
| KHFSQAEGSQLDEVR-AAQKNIDR-a1-b4 | pr\|PROT04\|MAEA | pr\|PROT04\|MAEA | intra-protein xl | 210307_LU | 246 | 61 | 185 | 2782.407 | 557.489 | 5 | 2.5 | 0.38 | 0.75 | **33.02** |
| LASDHKDIHSSVSR-IKDTVQK-a6-b2 | pr\|PROT05\|RMND5A | pr\|PROT05\|RMND5A | intra-protein xl | 210307_LU | 104 | 93 | 11 | 2519.341 | 630.843 | 4 | 2.6 | 0.39 | 0 | **32.64** |
| TPKDAASVR-IVSKGR-a3-b4 | pr\|PROT01\|RanBP9 | pr\|PROT01\|RanBP9 | intra-protein xl | 210307_LU | 221 | 248 | 27 | 1739.992 | 581.005 | 3 | 1.2 | 0.45 | 0 | **32.49** |
| LASDHKDIHSSVSR-VGKAIDK-a6-b3 | pr\|PROT05\|RMND5A | pr\|PROT05\|RMND5A | intra-protein xl | 210307_LU | 104 | 115 | 11 | 2418.292 | 605.581 | 4 | 2.1 | 0.46 | 0.48 | **32.14** |
| RKAVESIQAEDESAK-AAQKNIDR-a2-b4 | pr\|PROT04\|MAEA | pr\|PROT04\|MAEA | intra-protein xl | 210307_LU | 107 | 61 | 46 | 2712.412 | 679.111 | 4 | 2.9 | 0.42 | 0.61 | **32.12** |
| RETATCLAWCHDNKSR-AAQKNIDR-a14-b4 | pr\|PROT04\|MAEA | pr\|PROT04\|MAEA | intra-protein xl | 210307_LU | 209 | 61 | 148 | 3056.472 | 765.126 | 4 | 1.6 | 0.42 | 0 | **32.06** |
| STDQTVLEELASIKNR-IQKLVLAGR-a14-b3 | pr\|PROT01\|RanBP9 | pr\|PROT01\|RanBP9 | intra-protein xl | 210307_LU | 429 | 436 | 7 | 2937.655 | 735.422 | 4 | 1.7 | 0.41 | 0.82 | **31.94** |
| EAAEKFR-EAAEKFR-a5-b5 | pr\|PROT03\|Twa1 | pr\|PROT03\|Twa1 | intra-protein xl | 210307_LU | 72 | 72 | 0 | 1836.942 | 613.322 | 3 | 2.9 | 0.1 | 0 | **31.79** |
| RETATCLAWCHDNKSR-KHFSQAEGSQLDEVR-a14-b1 | pr\|PROT04\|MAEA | pr\|PROT04\|MAEA | intra-protein xl | 210307_LU | 209 | 246 | 37 | 3871.825 | 775.373 | 5 | 3.4 | 0.38 | 0 | **30.81** |
| CELTPLKYNTER-AKAEWEGK-a7-b2 | pr\|PROT02\|WDR26 | pr\|PROT02\|WDR26 | intra-protein xl | 210307_LU | 148 | 172 | 24 | 2578.28 | 860.435 | 3 | 2.5 | 0.49 | 0.69 | **30.79** |
| YLEYLEDGKVLEALQVLR-AEWEGKGTASR-a9-b6 | pr\|PROT02\|WDR26 | pr\|PROT02\|WDR26 | intra-protein xl | 210307_LU | 132 | 178 | 46 | 3478.815 | 870.712 | 4 | 4.8 | 0.35 | 0 | **30.36** |
| CELTPLKYNTER-MKFLLLQQK-a7-b2 | pr\|PROT02\|WDR26 | pr\|PROT02\|WDR26 | intra-protein xl | 210307_LU | 148 | 116 | 32 | 2808.498 | 703.132 | 4 | 2.3 | 0.42 | 0.72 | **30.36** |
| QDDKVVCPR-AAQKNIDR-a4-b4 | pr\|PROT04\|MAEA | pr\|PROT04\|MAEA | intra-protein xl | 210307_LU | 398 | 61 | 337 | 2168.104 | 543.034 | 4 | 1.7 | 0.29 | 0 | **29.91** |
| ESTPKLAK-EAAEKFR-a5-b5 | pr\|PROT03\|Twa1 | pr\|PROT03\|Twa1 | intra-protein xl | 210307_LU | 215 | 72 | 143 | 1860.006 | 621.01 | 3 | 3.7 | 0.1 | 0 | **29.32** |
| AAQKNIDRETSHVTMVVAELEK-KHFSQAEGSQLDEVR-a4-b1 | pr\|PROT04\|MAEA | pr\|PROT04\|MAEA | intra-protein xl | 210307_LU | 61 | 246 | 185 | 4336.19 | 1085.055 | 4 | 3.4 | 0.28 | 0 | **29.3** |
| KAVESIQAEDESAK-AAQKNIDR-a1-b4 | pr\|PROT04\|MAEA | pr\|PROT04\|MAEA | intra-protein xl | 210307_LU | 107 | 61 | 46 | 2556.311 | 853.111 | 3 | 3 | 0.32 | 0 | **29.21** |
| ETATCLAWCHDNKSR-KHFSQAEGSQLDEVR-a13-b1 | pr\|PROT04\|MAEA | pr\|PROT04\|MAEA | intra-protein xl | 210307_LU | 209 | 246 | 37 | 3715.722 | 929.938 | 4 | 3.1 | 0.3 | 0 | **29.19** |
| RETATCLAWCHDNKSR-RKAVESIQAEDESAK-a14-b2 | pr\|PROT04\|MAEA | pr\|PROT04\|MAEA | intra-protein xl | 210307_LU | 209 | 107 | 102 | 3801.829 | 761.374 | 5 | 3.2 | 0.29 | 0.33 | **28.84** |
| LKCPYCPMEQSPGDAK-DALNKXFNGSK-a2-b5 | pr\|PROT05\|RMND5A | pr\|PROT05\|RMND5A | intra-protein xl | 210307_LU | 397 | 389 | 8 | 3257.499 | 815.383 | 4 | 3.7 | 0.38 | 0.69 | **28.73** |
| YPSSVSAITDIKR-XCSKER-a12-b4 | pr\|PROT06\|ARMC8b | pr\|PROT06\|ARMC8b | intra-protein xl | 210307_LU | 355 | 300 | 55 | 2399.19 | 600.805 | 4 | 3 | 0.25 | 0 | **28.65** |
| KAVESIQAEDESAK-VPYETLNKR-a1-b8 | pr\|PROT04\|MAEA | pr\|PROT04\|MAEA | intra-protein xl | 210307_LU | 107 | 54 | 53 | 2760.425 | 921.149 | 3 | 2.5 | 0.3 | 0 | **28.37** |
| RETATCLAWCHDNKSR-KAVESIQAEDESAK-a14-b1 | pr\|PROT04\|MAEA | pr\|PROT04\|MAEA | intra-protein xl | 210307_LU | 209 | 107 | 102 | 3645.726 | 912.439 | 4 | 2.8 | 0.4 | 0 | **28.09** |
| KHFSQAEGSQLDEVR-VPYETLNKR-a1-b8 | pr\|PROT04\|MAEA | pr\|PROT04\|MAEA | intra-protein xl | 210307_LU | 246 | 54 | 192 | 2986.523 | 996.515 | 3 | 2.7 | 0.34 | 0 | **28.02** |
| LDHDLKHAHELR-NAVIGNNKQK-a6-b8 | pr\|PROT06\|ARMC8b | pr\|PROT06\|ARMC8b | intra-protein xl | 210307_LU | 362 | 62 | 300 | 2705.443 | 902.822 | 3 | 2.7 | 0.27 | 0 | **27.97** |
| TKEVFHFSQAEK-QDDKVVCPR-a2-b4 | pr\|PROT04\|MAEA | pr\|PROT04\|MAEA | intra-protein xl | 210307_LU | 405 | 398 | 7 | 2703.34 | 676.843 | 4 | 2.8 | 0.31 | 0 | **27.25** |
| VLHKFSGYGQLCER-DTVQKLASDHK-a4-b5 | pr\|PROT05\|RMND5A | pr\|PROT05\|RMND5A | intra-protein xl | 210307_LU | 41 | 98 | 57 | 3071.556 | 1024.86 | 3 | 2 | 0.34 | 0 | **27.2** |
| DKPIEMQLTSAK-XCSKER-a2-b4 | pr\|PROT06\|ARMC8b | pr\|PROT06\|ARMC8b | intra-protein xl | 210307_LU | 258 | 300 | 42 | 2323.128 | 581.79 | 4 | 2.3 | 0.26 | 0 | **27.11** |
| YPSSVSAITDIKR-MCSKER-a12-b4 | pr\|PROT06\|ARMC8b | pr\|PROT06\|ARMC8b | intra-protein xl | 210307_LU | 355 | 300 | 55 | 2383.193 | 596.806 | 4 | 2.1 | 0.2 | 0 | **26.84** |
| KHFSQAEGSQLDEVR-RKAVESIQAEDESAK-a1-b2 | pr\|PROT04\|MAEA | pr\|PROT04\|MAEA | intra-protein xl | 210307_LU | 246 | 107 | 139 | 3527.761 | 706.56 | 5 | 3.4 | 0.32 | 0.36 | **26.14** |
| MTDLSKGVIEEPK-ESTPKLAK-a6-b5 | pr\|PROT03\|Twa1 | pr\|PROT03\|Twa1 | intra-protein xl | 210307_LU | 245 | 215 | 30 | 2456.317 | 615.087 | 4 | 3.4 | 0.21 | 0.85 | **26.09** |
| EHSSDQPAAASVWKR-QDDKVVCPR-a14-b4 | pr\|PROT04\|MAEA | pr\|PROT04\|MAEA | intra-protein xl | 210307_LU | 144 | 398 | 254 | 2921.417 | 731.362 | 4 | 2.6 | 0.17 | 0 | **26.06** |
| MTDLSKGVIEEPK-KVKYPK-a6-b3 | pr\|PROT03\|Twa1 | pr\|PROT03\|Twa1 | intra-protein xl | 210307_LU | 245 | 236 | 9 | 2345.298 | 587.332 | 4 | 2.7 | 0.23 | 0.59 | **25.97** |
| AAQKNIDRETSHVTMVVAELEK-RKAVESIQAEDESAK-a4-b2 | pr\|PROT04\|MAEA | pr\|PROT04\|MAEA | intra-protein xl | 210307_LU | 61 | 107 | 46 | 4266.188 | 854.245 | 5 | 1.8 | 0.28 | 0.26 | **25.78** |
| AEWEGKGTASR-SKLLDK-a6-b2 | pr\|PROT02\|WDR26 | pr\|PROT02\|WDR26 | intra-protein xl | 210307_LU | 178 | 185 | 7 | 2031.068 | 678.03 | 3 | 2 | 0.29 | 0 | **25.58** |
| KVSLGEGRPPVLTASR-MCSKER-a1-b4 | pr\|PROT06\|ARMC8b | pr\|PROT06\|ARMC8b | intra-protein xl | 210307_LU | 388 | 300 | 88 | 2613.377 | 654.352 | 4 | 1.6 | 0.27 | 0 | **25.37** |
| ELEKVLHK-VGKAIDK-a4-b3 | pr\|PROT05\|RMND5A | pr\|PROT05\|RMND5A | intra-protein xl | 210307_LU | 37 | 115 | 78 | 1862.09 | 466.53 | 4 | 1.4 | 0.26 | 0 | **25.32** |
| SLNKLAQPLPMAHCANSR-TPQCYKEDGSSK-a4-b6 | pr\|PROT04\|MAEA | pr\|PROT04\|MAEA | intra-protein xl | 210307_LU | 346 | 327 | 19 | 3543.705 | 709.749 | 5 | 4 | 0.28 | 0 | **25.22** |
| AAQKNIDRETSHVTMVVAELEK-KAVESIQAEDESAK-a4-b1 | pr\|PROT04\|MAEA | pr\|PROT04\|MAEA | intra-protein xl | 210307_LU | 61 | 107 | 46 | 4110.094 | 1028.531 | 4 | 3.7 | 0.27 | 0 | **24.21** |
| TPQCYKEDGSSKSPDCPVCSR-SLNKLAQPLPXAHCANSR-a12-b4 | pr\|PROT04\|MAEA | pr\|PROT04\|MAEA | intra-protein xl | 210307_LU | 333 | 346 | 13 | 4618.125 | 1155.539 | 4 | 2.6 | 0.29 | 0 | **24.15** |
| KVSLGEGRPPVLTASR-NAVIGNNKQK-a1-b8 | pr\|PROT06\|ARMC8b | pr\|PROT06\|ARMC8b | intra-protein xl | 210307_LU | 388 | 62 | 326 | 2888.628 | 963.884 | 3 | 2.9 | 0.26 | 0 | **24.08** |
| LFDPDPQKVLQGVIDMK-NAVIGNNKQK-a8-b8 | pr\|PROT06\|ARMC8b | pr\|PROT06\|ARMC8b | intra-protein xl | 210307_LU | 45 | 62 | 17 | 3164.699 | 792.183 | 4 | 2.9 | 0.24 | 0 | **24.07** |
| SLNKLAQPLPXAHCANSR-TPQCYKEDGSSK-a4-b6 | pr\|PROT04\|MAEA | pr\|PROT04\|MAEA | intra-protein xl | 210307_LU | 346 | 327 | 19 | 3559.695 | 890.932 | 4 | 2.5 | 0.21 | 0 | **24.07** |
| KVSLGEGRPPVLTASR-KVSLGEGRPPVLTASR-a1-b1 | pr\|PROT06\|ARMC8b | pr\|PROT06\|ARMC8b | intra-protein xl | 210307_LU | 388 | 388 | 0 | 3469.988 | 579.339 | 6 | 4.4 | 0.29 | 0 | **24.04** |
| DKPIEMQLTSAK-NAVIGNNKQK-a2-b8 | pr\|PROT06\|ARMC8b | pr\|PROT06\|ARMC8b | intra-protein xl | 210307_LU | 258 | 62 | 196 | 2582.381 | 861.802 | 3 | 2.9 | 0.19 | 0 | **23.77** |
| KHFSQAEGSQLDEVR-KAVESIQAEDESAK-a1-b1 | pr\|PROT04\|MAEA | pr\|PROT04\|MAEA | intra-protein xl | 210307_LU | 246 | 107 | 139 | 3371.658 | 675.339 | 5 | 3 | 0.33 | 0 | **23.69** |
| ILEALKVR-VGKAIDK-a6-b3 | pr\|PROT05\|RMND5A | pr\|PROT05\|RMND5A | intra-protein xl | 210307_LU | 189 | 115 | 74 | 1808.116 | 453.037 | 4 | 1.5 | 0.19 | 0 | **23.53** |
| ETATCLAWCHDNKSR-AAQKNIDR-a13-b4 | pr\|PROT04\|MAEA | pr\|PROT04\|MAEA | intra-protein xl | 210307_LU | 209 | 61 | 148 | 2900.375 | 726.102 | 4 | 3.3 | 0.16 | 0.65 | **23.47** |
| MSYAEKPDEITKDEWMEK-MTDLSKGVIEEPK-a12-b6 | pr\|PROT03\|Twa1 | pr\|PROT03\|Twa1 | intra-protein xl | 210307_LU | 36 | 245 | 209 | 3812.823 | 954.213 | 4 | 3.7 | 0.21 | 0 | **23.25** |
| MTDLSKGVIEEPK-XTDLSKGVIEEPK-a6-b6 | pr\|PROT03\|Twa1 | pr\|PROT03\|Twa1 | intra-protein xl | 210307_LU | 245 | 245 | 0 | 3045.564 | 1016.196 | 3 | 4.7 | 0.19 | 0 | **23.17** |
| SLNKLAQPLPXAHCANSR-EDGSSKSPDCPVCSR-a4-b6 | pr\|PROT04\|MAEA | pr\|PROT04\|MAEA | intra-protein xl | 210307_LU | 346 | 333 | 13 | 3840.777 | 961.202 | 4 | 3 | 0.31 | 0 | **23.03** |
| ETATCLAWCHDNKSR-KAVESIQAEDESAK-a13-b1 | pr\|PROT04\|MAEA | pr\|PROT04\|MAEA | intra-protein xl | 210307_LU | 209 | 107 | 102 | 3489.624 | 1164.216 | 3 | 2.9 | 0.17 | 0 | **22.87** |
| KVSLGEGRPPVLTASR-YPSSVSAITDIKR-a1-b12 | pr\|PROT06\|ARMC8b | pr\|PROT06\|ARMC8b | intra-protein xl | 210307_LU | 388 | 355 | 33 | 3239.798 | 648.967 | 5 | 3.1 | 0.22 | 0 | **22.79** |
| SLNKLAQPLPMAHCANSR-EHSSDQPAAASVWKR-a4-b14 | pr\|PROT04\|MAEA | pr\|PROT04\|MAEA | intra-protein xl | 210307_LU | 346 | 144 | 202 | 3812.897 | 763.587 | 5 | 3.4 | 0.26 | 0 | **22.13** |
| ETATCLAWCHDNKSR-RKAVESIQAEDESAK-a13-b2 | pr\|PROT04\|MAEA | pr\|PROT04\|MAEA | intra-protein xl | 210307_LU | 209 | 107 | 102 | 3645.728 | 912.44 | 4 | 3.6 | 0.22 | 0 | **22.11** |
| AAQKNIDRETSHVTMVVAELEK-VPYETLNKR-a4-b8 | pr\|PROT04\|MAEA | pr\|PROT04\|MAEA | intra-protein xl | 210307_LU | 61 | 54 | 7 | 3724.96 | 621.834 | 6 | 3.8 | 0.31 | 0.53 | **22.1** |
| KVSLGEGRPPVLTASR-DKPIEMQLTSAK-a1-b2 | pr\|PROT06\|ARMC8b | pr\|PROT06\|ARMC8b | intra-protein xl | 210307_LU | 388 | 258 | 130 | 3163.736 | 633.755 | 5 | 2.9 | 0.34 | 0 | **22.04** |
| TPKDAASVR-SWSPKDK-a3-b5 | pr\|PROT01\|RanBP9 | pr\|PROT01\|RanBP9 | intra-protein xl | 210307_LU | 221 | 196 | 25 | 1928.006 | 643.676 | 3 | 2.8 | 0.17 | 0 | **22.02** |
| LLDKLQTYLPPSVMLPPR-CELTPLKYNTER-a4-b7 | pr\|PROT02\|WDR26 | pr\|PROT02\|WDR26 | intra-protein xl | 210307_LU | 189 | 148 | 41 | 3741.001 | 936.258 | 4 | 3.3 | 0.22 | 0 | **21.98** |
| QAAFKLYASLGANDEDIR-LDHDLKHAHELR-a5-b6 | pr\|PROT06\|ARMC8b | pr\|PROT06\|ARMC8b | intra-protein xl | 210307_LU | 373 | 362 | 11 | 3601.84 | 901.468 | 4 | 3.3 | 0.22 | 0 | **21.75** |
| TPQCYKEDGSSKSPDCPVCSR-SLNKLAQPLPMAHCANSR-a12-b4 | pr\|PROT04\|MAEA | pr\|PROT04\|MAEA | intra-protein xl | 210307_LU | 333 | 346 | 13 | 4602.13 | 1151.54 | 4 | 2.5 | 0.3 | 0 | **21.57** |
| MSYAEKPDEITKDEWXEK-XTDLSKGVIEEPK-a12-b6 | pr\|PROT03\|Twa1 | pr\|PROT03\|Twa1 | intra-protein xl | 210307_LU | 36 | 245 | 209 | 3844.814 | 769.971 | 5 | 4 | 0.28 | 0.73 | **21.34** |
| QAAFKLYASLGANDEDIRK-KVSLGEGRPPVLTASR-a5-b1 | pr\|PROT06\|ARMC8b | pr\|PROT06\|ARMC8b | intra-protein xl | 210307_LU | 373 | 388 | 15 | 3913.119 | 783.632 | 5 | 3.2 | 0.22 | 0.61 | **21.34** |
| SLNKLAQPLPMAHCANSR-VPYETLNKR-a4-b8 | pr\|PROT04\|MAEA | pr\|PROT04\|MAEA | intra-protein xl | 210307_LU | 346 | 54 | 292 | 3263.7 | 816.933 | 4 | 2.9 | 0.25 | 0.71 | **21.3** |
| SLNKLAQPLPMAHCANSR-EDGSSKSPDCPVCSR-a4-b6 | pr\|PROT04\|MAEA | pr\|PROT04\|MAEA | intra-protein xl | 210307_LU | 346 | 333 | 13 | 3824.78 | 1275.935 | 3 | 2.6 | 0.3 | 0 | **21.15** |
| AAQKNIDRETSHVTMVVAELEK-ETATCLAWCHDNKSR-a4-b13 | pr\|PROT04\|MAEA | pr\|PROT04\|MAEA | intra-protein xl | 210307_LU | 61 | 209 | 148 | 4454.159 | 891.84 | 5 | 4 | 0.24 | 0 | **20.79** |
| MSYAEKPDEITKDEWMEK-XTDLSKGVIEEPK-a12-b6 | pr\|PROT03\|Twa1 | pr\|PROT03\|Twa1 | intra-protein xl | 210307_LU | 36 | 245 | 209 | 3828.816 | 766.771 | 5 | 3.3 | 0.25 | 0.86 | **20.71** |
| MSYAEKPDEITKDEWXEK-MTDLSKGVIEEPK-a12-b6 | pr\|PROT03\|Twa1 | pr\|PROT03\|Twa1 | intra-protein xl | 210307_LU | 36 | 245 | 209 | 3828.815 | 766.771 | 5 | 2.9 | 0.15 | 0.75 | **20.46** |
| AAQKNIDRETSHVTXVVAELEK-KAVESIQAEDESAK-a4-b1 | pr\|PROT04\|MAEA | pr\|PROT04\|MAEA | intra-protein xl | 210307_LU | 61 | 107 | 46 | 4126.086 | 826.225 | 5 | 2.9 | 0.26 | 0.88 | **20.43** |
| KVSLGEGRPPVLTASR-XCSKER-a1-b4 | pr\|PROT06\|ARMC8b | pr\|PROT06\|ARMC8b | intra-protein xl | 210307_LU | 388 | 300 | 88 | 2629.375 | 658.352 | 4 | 2.8 | 0.14 | 0 | **20.43** |
| AAQKNIDRETSHVTXVVAELEK-RKAVESIQAEDESAK-a4-b2 | pr\|PROT04\|MAEA | pr\|PROT04\|MAEA | intra-protein xl | 210307_LU | 61 | 107 | 46 | 4282.188 | 857.446 | 5 | 3.1 | 0.16 | 0.73 | **19.99** |
| LKCPYCPXEQSPGDAK-DALNKMFNGSK-a2-b5 | pr\|PROT05\|RMND5A | pr\|PROT05\|RMND5A | intra-protein xl | 210307_LU | 397 | 389 | 8 | 3257.497 | 815.382 | 4 | 3.3 | 0.22 | 0 | **19.82** |
| KHFSQAEGSQLDEVR-QDDKVVCPR-a1-b4 | pr\|PROT04\|MAEA | pr\|PROT04\|MAEA | intra-protein xl | 210307_LU | 246 | 398 | 152 | 2983.453 | 995.492 | 3 | 2.5 | 0.12 | 0.64 | **19.32** |
| KVSLGEGRPPVLTASR-LDHDLKHAHELR-a1-b6 | pr\|PROT06\|ARMC8b | pr\|PROT06\|ARMC8b | intra-protein xl | 210307_LU | 388 | 362 | 26 | 3286.802 | 822.708 | 4 | 3.5 | 0.19 | 0 | **18.9** |
| SLNKLAQPLPXAHCANSR-QDDKVVCPR-a4-b4 | pr\|PROT04\|MAEA | pr\|PROT04\|MAEA | intra-protein xl | 210307_LU | 346 | 398 | 52 | 3276.629 | 656.334 | 5 | 3.9 | 0.2 | 0 | **17.73** |
| QQTTDNNPPMKLVCGHIISR-ILEALKVR-a11-b6 | pr\|PROT05\|RMND5A | pr\|PROT05\|RMND5A | intra-protein xl | 210307_LU | 375 | 189 | 186 | 3386.826 | 847.714 | 4 | 2.9 | 0.11 | 0 | **17.28** |
| TDDNCIVLKTLPCLVR-TDDNCIVLKTLPCLVR-a9-b9 | pr\|PROT06\|ARMC8b | pr\|PROT06\|ARMC8b | intra-protein xl | 210307_LU | 289 | 289 | 0 | 3970.05 | 993.52 | 4 | 2.6 | 0.28 | 0 | **16.16** |

**Description of the column headers**

| | |
|---|---|
| *Id* | Assigned peptides and cross-linking sites within the *peptide* sequences. The longer peptide is designated as (a)lpha, the shorter as (b)eta. |
| *Protein1* | SwissProt/UniProt accession number and identifier of the protein 1 (containing peptide desginated as alpha). |
| *Protein2* | SwissProt/UniProt accession number and identifier of the protein 2 (containing peptide desginated as beta). |
| *XLType* | Intra- or inter-protein, sometimes also ambiguous. |
| *Spectrum* | References the actual MS/MS spectra that were used for the assignment. |
| *AbsPos1* | Position in the *protein* sequence of protein 1. |
| *AbsPos2* | Position in the *protein* sequence of protein 2. |
| *deltaAA* | Distance in the primary sequence (makes only sense for intra-proteins cross-links). |
| *Mr* | Molecular mass calculated from experimental m/z and z (neutral mass). |
| *Mz* | Experimentally observed mass-to-charge ratio of the precursor ion in Da. |
| *z* | Experimentally observed precursor charge. |
| *Error [ppm]* | Deviation between experimental and theoretical mass in ppm. |
| *TIC* | Fraction of the total ion current covered by assigned fragment ions. |
| *delta score* | Relative score for the top hit in relation to the second best hit for this spectrum, calculated as Id-score of 2nd best hit divided by Id-score of best hit. "0" means no second best hit exists. |
| *Id-score* | Identification score as assigned by xQuest. The higher, the better. |
| *X* | in the amino acid sequence denotes an oxidized Met residue |
